# Mechanistic insights into Enterocin C targeting the undecaprenyl phosphate recycling protein BacA

**DOI:** 10.1101/2025.05.12.652377

**Authors:** Victor Folcher, Rodolphe Auger, Clara Louche, Pascale Serror, Thierry Touzé

## Abstract

*Enterococcus faecalis* is an important opportunistic pathogen responsible for healthcare-associated infections. It is intrinsically resistant to various antibiotics, particularly to cephalosporins and vancomycin, creating an urgent need for alternative therapeutics. In this context, bacteriocins warrant investigations as a potential source of medical-use antibiotics. Herein, we demonstrate that Enterocin C, a class IIb two-peptide bacteriocin, specifically targets the membrane embedded undecaprenyl phosphate recycling protein BacA from enterococci as a cell surface receptor. Using biochemical and biophysical methods, supported by AlphaFold2 modelling and mutagenesis, we deciphered the EntC’s molecular interaction pattern with its target, marking the first mechanistic insight of a two-peptide bacteriocin. The two peptides act cooperatively at nanomolar concentrations to interact with the outward-open catalytic pocket of BacA: the peptide EntC1 docks deeply into the catalytic site, inhibits BacA’s enzymatic activity and enables the binding of peptide EntC2, eliciting membrane permeabilization, eventually leading to cell death. A comparative analysis with LcnG, a homologous bacteriocin, reveals a conserved interaction pattern, paving the way for bioengineering of these bacteriocins and developing tailored antimicrobial strategies.

## Introduction

The emergence of multidrug-resistant (MDR) and extensively drug-resistant (XDR) bacterial strains presents an unprecedented challenge to humanity that is expected to increase in the years to come. The World Health Organization (WHO) identified this issue as one of the top ten most critical global health concerns, stressing the urgent need for research into therapeutic alternatives (Naghavi *et al*, 2024). The pathogenic bacteria responsible for life-threatening nosocomial infections and developing antibiotic resistance are collectively termed “ESKAPE pathogens”, which encompass *Enterococcus* sp. (*Enterococcus faecium* and *Enterococcus faecalis*), *Staphylococcus aureus*, *Klebsiella pneumoniae*, *Acinetobacter baumannii*, *Pseudomonas aeruginosa*, and *Enterobacter* species (Venkateswaran *et al*, 2023). In response to this escalating antibiotic resistance crisis, bacteriocins are emerging as promising alternatives. These antimicrobial peptides are produced by bacteria across all phyla through ribosomal synthesis (Sugrue *et al*, 2024). They are widespread in nature and exhibit diversity in their structures and mechanisms of action. They show a narrow to broad inhibitory activity and are featured by high antibacterial activities, causing bacterial cell death at nanomolar to picomolar concentrations, with minimal or no cytotoxicity effects on eukaryotic cells. Despite their numerous advantages, only a few bacteriocins have progressed beyond preclinical trials, and none are currently employed in human therapy, which can be explained by a lack of understanding of their mechanisms of action. Elucidating these mechanisms is therefore critical in selecting them for the treatment of antibiotic-resistant bacterial infections.

Bacteriocins are categorized into two classes based on whether their founding peptides undergo post-translational modifications (class I) or not (class II) (Sugrue *et al*, 2024). Class IIb bacteriocins comprise two linear peptides that permeabilize cell membranes of target cells through mechanisms that remain to be elucidated (Nissen-Meyer *et al*, 2010, 2011). Examples of class IIb bacteriocins include the best studied Plantaricin JK and four homologous bacteriocins, Lactococcin G (LcnG), lactococcin Q (LcnQ), Enterocin 1071 (Ent1071) and Enterocin C (EntC), among at least 15 class IIb bacteriocins. All these bacteriocins apparently act via a cell surface receptor-mediated process conferring them high specificity of action. Genomic analysis of resistant clones suggests that Plantaricin JK targets a protein from the amino acid-polyamine-organocation (APC) family (Oppegård *et al*, 2016; Ekblad *et al*, 2017). LcnG, produced by *Lactococcus lactis,* consists of peptides LcnG_α_ (39 amino acids) and LcnG_β_ (35 amino acids). While these peptides are ineffective individually, they exhibit antibacterial activity at nanomolar concentrations against *L. lactis* strains when combined in a 1:1 ratio (Nissen-Meyer *et al*, 1992; Moll *et al*, 1996a; Kjos *et al*, 2014). LcnG selectively permeabilizes the cell membranes to monovalent cations (Moll *et al*, 1998, 1996a). Whole-genome sequencing of LcnG-resistant clones revealed stop codons or frameshifts mutations in the *bacA* gene indicating that the protein BacA likely constitutes the cell surface receptor of LcnG (Kjos *et al*, 2014). BacA is a membrane-embedded protein, which is widely conserved in bacteria. It is involved in the recycling of the undecaprenyl phosphate (C_55_-P), a lipid carrier enabling peptidoglycan subunit translocation across the plasma membrane **(Fig. EV1)** (El Ghachi *et al*, 2004). C_55_-P forms the so-called lipid II when bound to the peptidoglycan disaccharide-pentapeptide subunit. Following the subunit transfer to the nascent peptidoglycan, the lipid carrier is released as undecaprenyl pyrophosphate (C_55_-PP), which must be recycled to sustain peptidoglycan, and other cell wall glycopolymers, biosynthesis (Manat *et al*, 2014). The recycling consists of C_55_-PP dephosphorylation followed by the flip of C_55_-P to the inner side of the membrane. Either BacA or a protein from the PAP2 superfamily catalyze C_55_-PP dephosphorylation (El Ghachi *et al*, 2005). According to its crystal structure, which highly resembles that of transporter proteins, BacA was hypothesized to also catalyze the flip of C_55_-P (El Ghachi *et al*, 2018; Workman *et al*, 2018). Recent studies have identified additional membrane proteins with C_55_-P flippase activity, including members of the DedA family and DUF368-containing proteins (Roney & Rudner, 2023; Sit *et al*, 2023). BacA from *L. lactis* may act as a docking site for LcnG, potentially leading to pore formation. C_55_-P recycling is essential for bacterial viability; therefore, the cytotoxicity of LcnG may also originate from the inhibition of BacA activity. However, *L. lactis* encodes PAP2 enzymes that may perform redundant activity with BacA as observed in *E. coli* and *Bacillus subtilis* (Touzé *et al*, 2008; Zhao *et al*, 2016), so BacA activity may not be essential *per se* in this bacterium. In the present study, we have investigated the mechanism of action of Enterocin C (EntC), an LcnG homologue produced by *E. faecalis* C901 strain (**Fig. 1A**). EntC is composed of EntC1 and EntC2 peptides, and it is active against the opportunistic pathogen *E. faecalis* (Maldonado-Barragán *et al*, 2009). We demonstrated that EntC also relies on BacA to exert its antibacterial activity and its binding mode to BacA was investigated in depth using complementary *in vitro* approaches from which, in combination with AlphaFold2 modelling and mutagenesis, we could infer a model of action. Our findings demonstrate that EntC1 binds to the outward-open catalytic pocket of BacA, inhibiting its enzymatic activity and enabling the further binding of EntC2 in a cooperative manner. This latter interaction stabilizes the tripartite complex and triggers membrane permeabilization likely by anchoring the peptides, as one functional unit, deeply into the lipid bilayer, ultimately leading to cell death. We herein describe, for the first time, the binding mode of a two-peptide bacteriocin to its membrane-embedded target paving the way for the further development and bioengineering of this class II bacteriocin for narrow-spectrum antibiotics with minimal impact on microbiota.

**Fig. 1.**
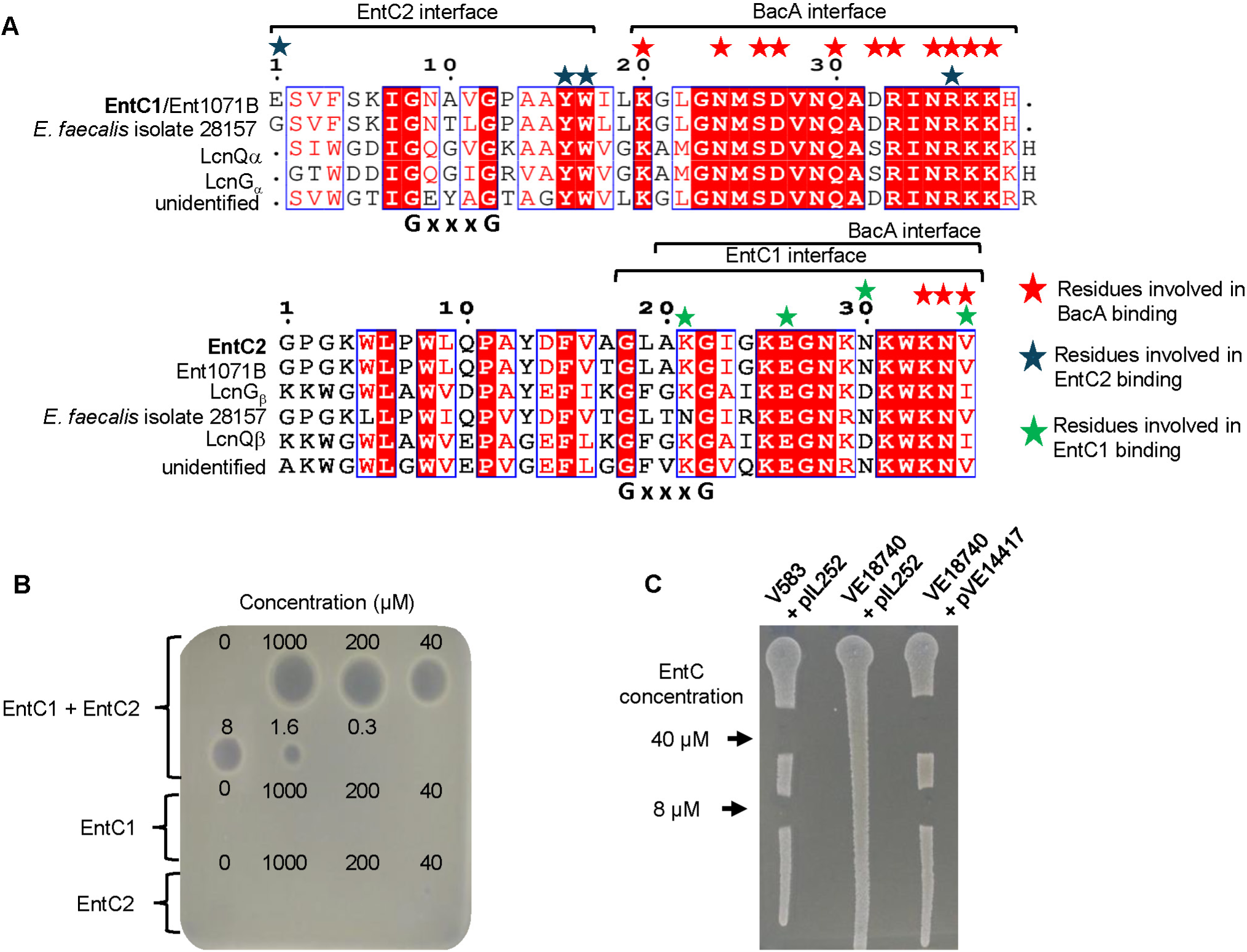
Sequence alignment and activity measurement of EntC. **(A)** Sequence alignment of EntC-related peptides. The interface regions and residues involved in binding, according to AlphaFold2 models, are indicated above the sequences (red stars indicate residues involved in BacA binding, blue stars for EntC2 binding and green stars for EntC1 binding). Blue boxes highlight similar sequences, identical sequences are filled in red. The GxxxG motif common to all aligned peptides is highlighted below the sequences. **(B)** Dose effect of EntC peptides on *E. faecalis* V583 strain. 6 µL-droplets of peptide solutions at the indicated concentrations are spotted on a lawn of *E. faecalis* V583 cells. Results are representative of three independent experiments. **(C)** Activity measurement of EntC on V583 and VE18740 (Δ*bacA*) cells carrying plasmids pIL252 (empty vector) or pVE14417 (*bacA_ef_*-carrying pIL252 vector). The indicated cells were patched on BHI agar plate and a drop of EntC solution at the indicated concentration was spotted. Results are representative of three independent experiments.

## Results

### BacA is targeted by EntC peptides

To study the mechanism of action of EntC, EntC1 and EntC2 mature peptides, *i.e.* without their N-terminal signal sequence (**Fig. 1A**), were obtained by chemical synthesis. Spot-on-lawn assays demonstrate the susceptibility of the clinical *E. faecalis* vancomycin-resistant V583 strain to the synthetic peptides when they were combined at equimolar concentrations in the micromolar range (**Fig. 1B**). In contrast, neither peptide exhibited antibacterial activity when applied individually up to 1 mM. The minimum inhibitory concentration (MIC) of EntC on V583 strain was determined in different culture media demonstrating varying MIC values: 100 nM in BHI, 50 nM in M17 and 2 nM in MRS (**Appendix Fig. S1**). This finding aligns with previous reports according to which growth in MRS medium optimizes bacteriocins efficacy in comparison to other media for a yet unknown reason (Yang *et al*, 2018; Todorov & Dicks, 2009). To establish whether BacA is the target of EntC, the *bacA*-deleted strain VE18740 was generated. VE18740 strain displays a similar growth phenotype as the wild-type strain in laboratory conditions and it exhibited a full resistance to EntC (**Fig. 1C**). The susceptibility of VE18740 strain to EntC was fully restored upon the expression of *bacA* from a plasmid (**Fig. 1C**). Our data demonstrate that the peptides exert a synergistic antibacterial activity against *E. faecalis* that is fully dependent on BacA protein.

### EntC inhibits the enzymatic activity of BacA

To investigate the mode of binding of EntC to BacA, an N-terminal His_6_-tagged BacA from *E. faecalis* (BacA*_ef_*) was overproduced in *E. coli,* solubilized in *n*-dodecyl-β-D-maltoside (DDM) detergent and purified to homogeneity by affinity purification followed by gel filtration (**Fig. EV2A, B**). To assess the functionality of the purified BacA*_ef_*, its C_55_-PP phosphatase activity was characterized. The optimal pH for BacA*_ef_* was determined between pH 6 and 8 (**Fig. EV2C**) and the presence of EDTA inhibited its activity **(Fig. EV2D**), as previously reported with BacA from *E. coli* (BacA*_ec_*) (Manat *et al*, 2015). The activity was restored upon the addition of CaCl_2_ following EDTA treatment, and to a lesser extent by MgCl_2_, MnCl_2_ and CoCl_2_ (**Fig. EV2E, F**). The specific activity was determined to be 27.0 ± 4.6 µmol/min/mg in the optimal conditions as compared to 11.3 ± 1.3 µmol/min/mg for BacA*_ec_*. When added at 50 µM, both EntC1 and EntC mixture (EntC1 and EntC2 in 1:1 molar ratio) completely inhibited BacA*_ef_*, while no effect was observed with EntC2 alone (**Fig. 2A**). EntC and EntC1 exhibited a dose-dependent inhibition of BacA, with IC_50_ values of 0.4 ± 0.1 µM and 0.8 ± 0.2 µM, respectively (**Fig. 2A**). These data demonstrate that EntC1 directly binds BacA*_ef_* and blocks its catalytic cycle. Notably, a decrease of the IC_50_ value was observed in the presence of both peptides as compared to EntC1 with a significant p-value (0.0007). The activity of BacA*_ec_*, which shares 41% sequence identity with BacA*_ef_* (**Fig. EV3**), was not inhibited by EntC up to 50 µM, highlighting the specificity of this bacteriocin.

**Fig. 2.**
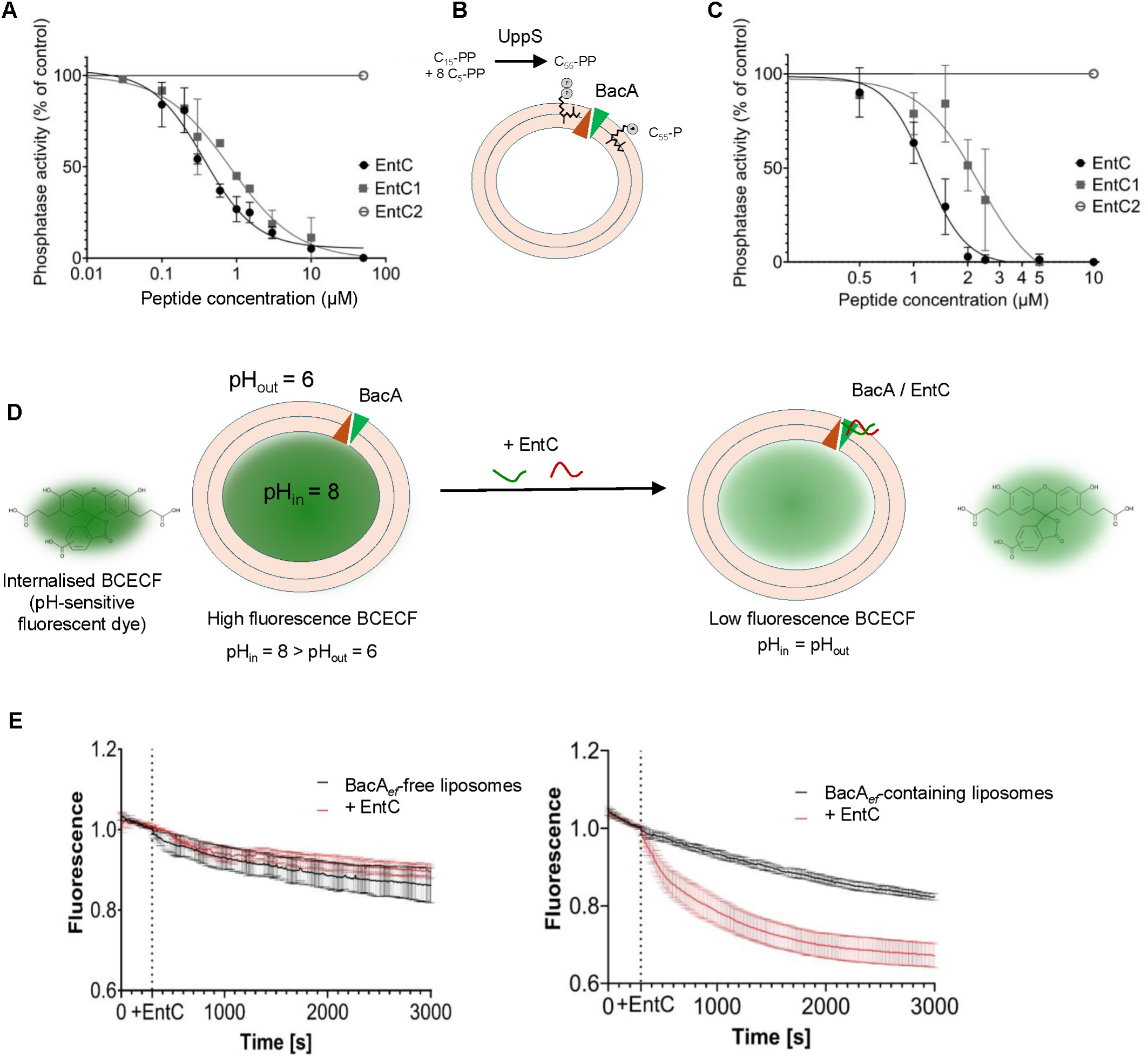
Effect of EntC on BacA*_ef_* C_55_-PP phosphatase activity and membrane integrity. (**A**) Dose effect of EntC peptides on BacA C_55_-PP phosphatase activity in DDM micelles. (**B**) Schematic representation of UppS-coupled C_55_-PP phosphatase assay in liposome. (**C**) Dose effect of EntC peptides on BacA C_55_-PP phosphatase activity in liposome. (**D**) Schematic representation of the permeabilization assay using the BCECF pH sensitive dye. BCECF is internalized in the lumen of liposomes at internal pH = 8 and liposomes are dispersed in buffer at pH = 6 to assess membrane integrity over time after the addition of EntC. (**E**) Membrane permeabilization activity of EntC. Fluorescence measurements are normalized to values recorded prior to the addition of 5 µM EntC (dashed line). The panel shows results obtained with BCECF-containing liposomes without (left panel) or with BacA*_ef_* reconstituted into the lipid bilayer (right panel). Results are the average of three independent experiments and error bars indicate standard deviations.

To assess EntC activity towards BacA*_ef_* in a membrane-like environment, the latter was reconstituted in liposomes. A coupled enzymatic assay was developed using *E. coli* UppS (C_55_-PP synthase) and water-soluble C_55_-PP precursors in order to synthesise C_55_-PP in the liposome-containing reaction mixture and to allow its partition in the lipid bilayer (**Fig. 2B**). Using BacA-free liposomes as a control, the synthesis of C_55_-PP was observed upon the addition of UppS and C_55_-PP precursors. A fraction of this C_55_-PP was further converted into C_55_-P with BacA-containing liposomes assessing the proper folding of BacA in the lipid bilayer and allowing activity measurement. We then showed that EntC and EntC1 fully inhibited BacA*_ef_* activity when added at 5 µM, while EntC2 had no effect (**Fig. 2C**). A dose-dependent inhibition of BacA*_ef_* by EntC and EntC1 was observed with IC_50_ values of 1.2 ± 0.1 µM and 2.2 ± 0.3 µM, respectively (**Fig. 2C**), which closely mirrors the effect of the peptides observed in micelles. At the highest concentration tested (10 µM), neither EntC nor EntC1 had an inhibitory effect on the activity of BacA*_ec_* when similarly reconstituted in liposomes.

### EntC permeabilizes BacA-containing liposomes

LcnG was previously described to cause membrane permeabilization in *L. lactis* cells (Moll *et al*, 1996a). To investigate whether EntC is able to disrupt lipid bilayers in a BacA-dependent manner, we used a pH-sensitive non-permeant fluorescent dye, BCECF, that was internalized in the lumen of liposomes at internal pH 8. Then, BCECF fluorescence was monitored over time after diluting the liposomes in a lower pH buffer (pH 6) (**Fig. 2D**). BacA-liposomes and protein-free liposomes displayed the same slow decrease of BCECF fluorescence over time in the absence of EntC (1.10^-4^ sec^-1^) (**Fig. 2E**). The addition of 5 µM of EntC to BacA*_ef_*-liposomes (peptide to lipid molar ratio, c.a. 1:1300) strongly enhanced the quenching of BCECF by the lower external pH (decrease rate about 6.10^-4^ sec^-1^), while it had no effect on protein-free liposomes (**Fig. 2E**). In order to determine whether the presence of any membrane protein would sensitize the liposomes to EntC, we also tested PgpB-liposomes. PgpB is a PAP2 C_55_-PP phosphatase from *E. coli* with a completely different fold and catalytic mechanism as compared to BacA (Workman *et al*, 2018; Fan *et al*, 2014; El Ghachi *et al*, 2018). In this case, EntC had no permeabilization effect (**Appendix Fig. S2**). These results indicate that EntC dissipates the pH gradient across the membrane of liposomes in a BacA-dependent manner, supporting that EntC directly parasitizes BacA to elicit the permeabilization of target cell membranes.

### EntC peptides form a complex with BacA in 1:1 stoichiometry

In order to highlight the formation of complexes between EntC peptides and BacA, *in vitro* crosslinking experiments were performed using the amine-reactive, membrane-permeable and non-cleavable crosslinker disuccinimidyl suberate (DSS). The peptides and BacA were incubated at equimolar concentrations (i.e. 17 µM) in the presence of a molar excess of DSS (100- or 1000-fold excess), the crosslinking reaction was quenched, the mixture was resolved by SDS-PAGE and BacA was revealed by western blotting. As shown in **Fig. 3A**, in the presence of EntC and DSS, two faint bands appeared above the BacA monomer, which likely correspond to BacA complexed with one and two peptides, respectively, according to their apparent molecular mass. In the presence of EntC1 alone, a strong and unique band was observed above the BacA monomer with a molecular mass corresponding to a BacA:EntC1 complex with a 1:1 stoichiometry (**Fig. 3A**). When EntC2 was added alone, a very faint band was also observed above BacA, slightly below the BacA:EntC1 complex, which is relevant to their molecular mass difference. Notably, covalent dimer of BacA were also observed at c.a. 60 kDa in the presence of DSS (**Fig. 3A**). BacA*_ec_* was previously found to crystallize as dimers by Workman et al. but whether the dimers were biologically relevant was not determined (Workman *et al*, 2018). Herein, the fact that BacA*_ef_* also dimerizes in DDM micelles supports a physiologically relevant oligomerization. Interestingly, in the presence of EntC1, the amount of covalent BacA dimers was weakened by fourfold as judged from densitometry analysis (p-value = 0.005) and no BacA dimers cross-linked with EntC1 were detected (**Fig. 3A**). Together, these results show that both peptides are able to bind BacA separately, but with a much greater propensity of binding for EntC1, and EntC1 apparently interferes with BacA dimerization. The band corresponding to the covalent BacA:EntC1 complex was further analyzed by mass spectrometry after trypsin digestion. As expected, peptides from both BacA and EntC1 were recovered, with 55% and 69% coverage, respectively (**Appendix Fig. S3**). Unfortunately, no cross-linked BacA:EntC1 peptide, that could have helped us to determine their interface, was recovered, presumably due to limited coverage.

**Fig. 3.**
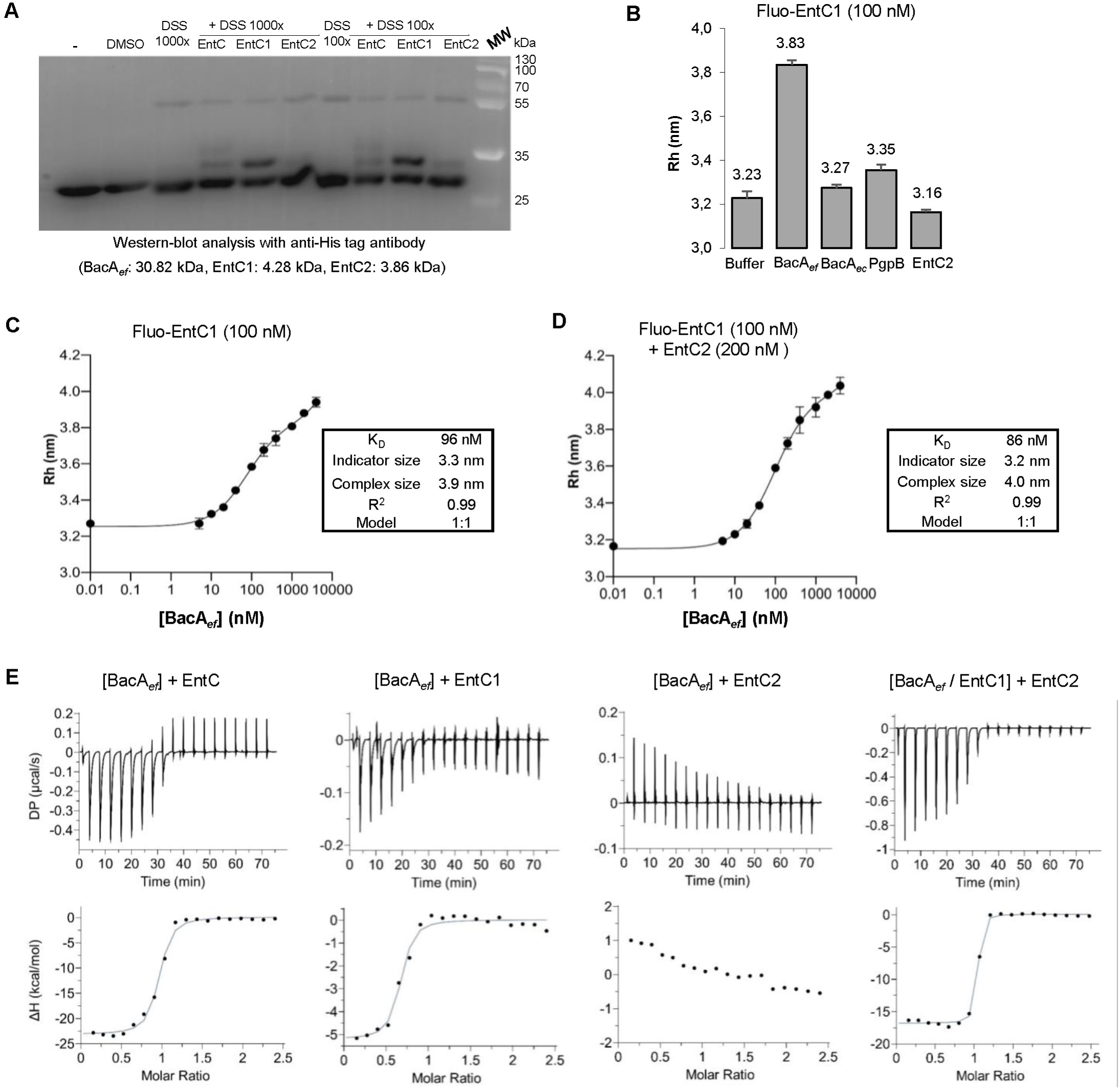
Interaction measurement between BacA*_ef_* and EntC peptides. (**A**) DSS Crosslinking of BacA*_ef_* sample in the absence or the presence of EntC peptides added at equimolar concentrations (17 µM) at two different protein to DSS ratio. (**B**) Flow-Induced Dispersion Analysis (FIDA) for the determination of hydrodynamic radii (Rh) of Fluorescein-labelled EntC1 in the absence or the presence of unlabelled proteins or EntC2. Results are the average of three independent experiments and error bars indicate standard deviations. **(C,D)** FIDA determination of the dissociation constant of Fluo-EntC1 towards BacA*_ef_* in the absence (**C**) and in the presence of EntC2 (**D**). Results are the average of three independent experiments and error bars indicate standard deviations. **(E)** Isothermal Titration Calorimetry (ITC) scans for BacA*_ef_* titration with EntC peptides. Upper panels, ITC thermogram responses for the titration of EntC peptides (400 µM) into of the indicated mixture indicated between brackets (32 µM). Lower panels, heat profile from peak integration of the ITC thermograms. The data shown are representative of experiments performed in triplicate.

### EntC1 binds BacA*_ef_* with high affinity

Flow-Induced Dispersion Analysis (FIDA) is used to determine the hydrodynamic radius (Rh) of proteins or peptides based on the fact that the laminar flow profile of particles through narrow capillaries (75 μm diameter sized) is governed by their Rh (Monti *et al*, 2024). The Rh is then determined from the measurement of the flow profile of a fluorescently labelled species. FIDA can be used to monitor protein-protein interaction through changes in the flow profile of the fluorescent species upon interaction with partners (≥ 10% increment of the Rh can be detected). To evaluate the BacA*_ef_*:EntC interaction, EntC peptides were then individually and randomly labelled at primary amino groups with fluorescein. Unfortunately, the labelling of EntC2 was not successful in contrast to EntC1 labelling (see materials and methods). The dispersion of fluorescein-EntC1 (Fluo-EntC1) in detergent-free solution yielded an Rh of 0.69 nm, and the peptide was found to adhere to the capillary as judged from the deviation of the Taylorgram from the ideal Gaussian shape expected for pure dispersion (**Appendix Fig. S4**). Upon dispersion of Fluo-EntC1 in a buffer containing 0.1 % DDM, an Rh of 3.2 nm was yielded and no adherence to the capillary was noted (**Appendix Fig. S4, Fig. 3B**). This size is likely representative of a particle corresponding to the peptide within a DDM micelle as judged from the size increment. Previous studies using circular dichroism and NMR spectroscopy have revealed that both LcnG peptides were unstructured in water, but became structured upon exposure to micelles or liposomes (Rogne *et al*, 2008; Hauge *et al*, 1998). Our results strongly suggest that EntC1 is sharing this feature; EntC1 interacts with micelles to the contact of which it may get, at least partially, folded. Upon dispersion of EntC1 in a buffer containing BacA*_ef_* micelles, an Rh of 3.8 nm was yielded, demonstrating the binding of EntC1 to BacA*_ef_*-micelles according to the significant size increment, while no significant change of EntC1 Rh was observed in the presence BacA*_ec_* or PgpB micelles (**Fig. 3B**). The addition of EntC2 did not induce an increase of EntC1 Rh. However, due to the small size of the peptide compared to the size of the micelle, we could not infer whether EntC2 interacted with EntC1-containing micelles or not (**Fig. 3B**). We then measured the dispersion of Fluo-EntC1 upon titration with BacA*_ef_* micelles (**Fig**. **3C**). The titration curve was best fitted with a 1:1 binding model yielding a *K*_d_ of 96 nM. The addition of EntC2 in the latter experiment at twice the concentration of EntC1, displayed a similar binding curve, best fitted with 1:1 binding model, and yielding a *K*_d_ of 86 nM (**Fig**. **3D**).

### EntC peptides bind cooperatively with BacA*_ef_*

As EntC2 binding parameters could not be determined by FIDA, the binding was also assessed in solution by Isothermal Titration Calorimetry (ITC). The peptides, prepared in DDM solution, were titrated in a solution of BacA*_ef_* micelles. The titration of EntC into a BacA*_ef_* solution exhibited an exothermic binding curve, best fitted with single binding site model and providing a *K*_d_ of 196 nM (**Fig**. **3E** **and Table 1**). The binding was governed by an important enthalpy change (ΔH −23.1 kcal/mol) with a strong entropy penalty (-TΔS of 14.1 kcal/mol). As a control, no binding was observed upon titration of EntC into BacA*_ec_* (**Appendix Fig. S5**). The titration of EntC1 into BacA*_ef_* also exhibited an exothermic reaction providing a *K*_d_ of 306 nM, with a similar free energy change (ΔG −9 kcal/mol) but a much smaller magnitude of heat release (ΔH = −5.4 kcal/mol) and a favorable entropy change (-TΔS −3.4 kcal/mol) (**Fig**. **3E** **and Table 1**). No binding was observed upon titration of EntC2 with BacA*_ef_*, nor titration of EntC2 with EntC1 (**Appendix Fig**. **S5**). These data demonstrate a favorable binding of EntC1 to BacA*_ef_*, which is consistent with the inhibition of BacA activity and FIDA analyzes. It also suggests that EntC2 binds to the BacA*_ef_*:EntC1 pre-complex as the thermodynamic parameters significantly differ when EntC2 was added together with EntC1, while it does not apparently bind directly to BacA*_ef_*, nor to EntC1. Indeed, the titration of EntC2 into a mixture of BacA*_ef_* and EntC1 at equimolar concentration displayed an exothermic binding curve featured by a favorable enthalpy change (ΔH - 16.8 kcal/mol) together with entropy penalty (-TΔS of 6.5 kcal/mol) (**Fig. 3E**). Specifically, the formation of a pre-complex enables the subsequent binding of EntC2. The affinity of this second binding event is high, about 10-fold higher (21 nM) than EntC1 binding, and the stoichiometry was very close to 1:1. Consistently, the titration of EntC1 into a mixture containing BacA*_ef_* and EntC2 displayed a similar binding pattern as the one obtained with titration of EntC into BacA*_ef_* (**Appendix Fig**. **S5 and Table 1**).

**Table 1.**
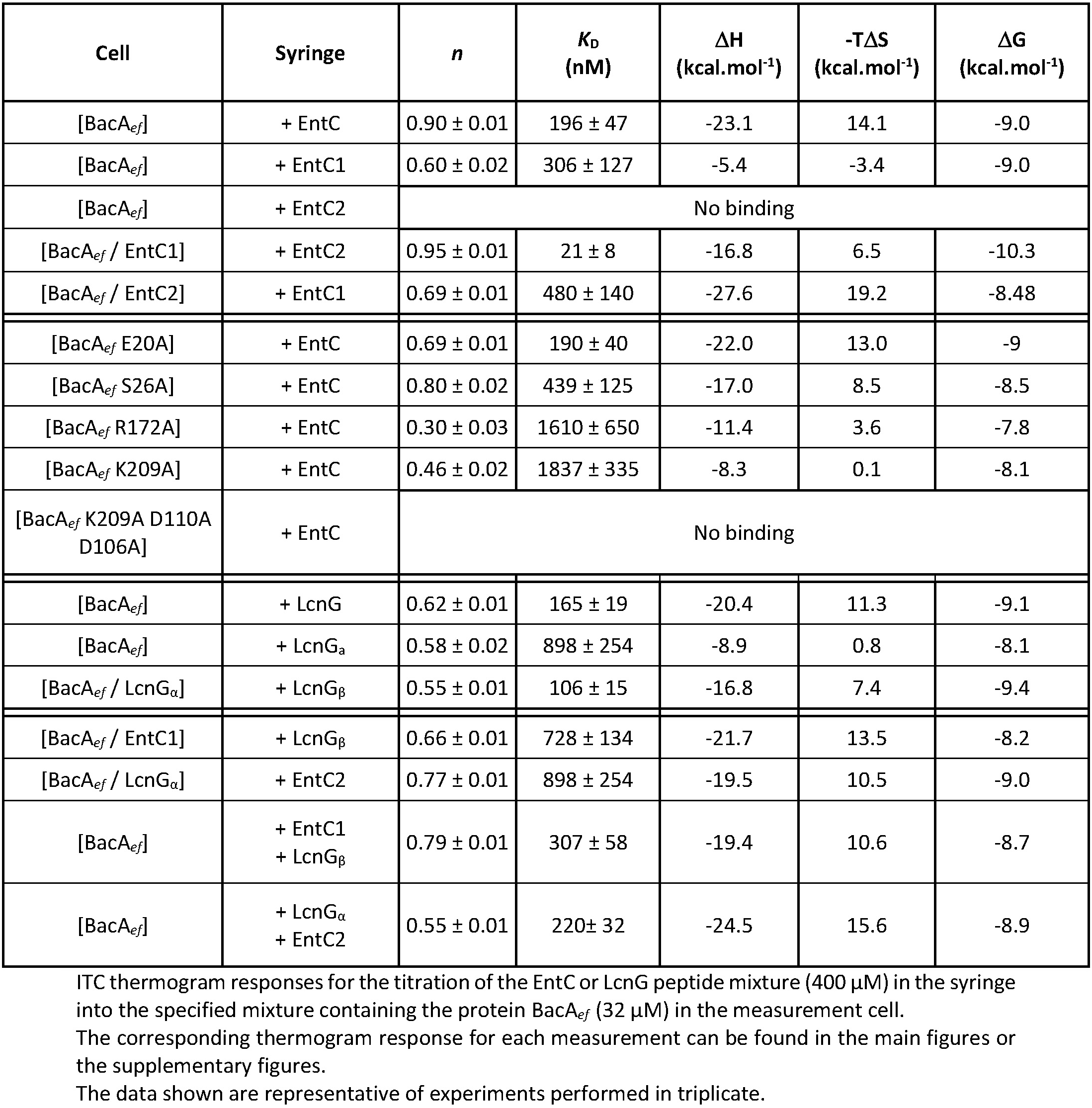
ITC scans thermodynamic parameters.

### BacA*_ef_* is gradually and highly stabilized upon binding with EntC peptides

ITC measurements strongly suggests that EntC binding stabilizes BacA*_ef_* according to the enthalpy driven binding associated with a strong entropy penalty. To further investigate the extent of this stabilization, we measured the melting temperature (T_m_) and enthalpy of denaturation (ΔH) of BacA*_ef_* in the absence and presence of peptides using Differential Scanning Calorimetry (DSC) (**Fig. 4**). Importantly, none of the peptides themselves (individually or combined) displayed a DSC signal when prepared in DDM solution at 66 µM. The thermogram of BacA*_ef_* at 33 µM displayed a Tm of 66.3°C and a ΔH of 26.7 kcal/mol (**Fig. 4**). The addition of EntC2 at twice the concentration of BacA*_ef_* did not change the DSC signal, which correlates with the lack of interaction as emphasized by ITC. In contrast, the thermogram of BacA*_ef_* in the presence of EntC1 displayed a single endothermal peak with a significant increase of the T_m_ and ΔH to 73°C and 45.3 kcal/mol, respectively (**Fig. 4**). The addition of complete EntC further enhanced the stabilization of the complex yielding a T_m_ of 83.8°C and a ΔH of 83.7 kcal/mol (**Fig. 4**). Whether the peptides fold upon contact with the micelles and further gain folding upon binding to BacA*_ef_* is an open question. It is not possible to evaluate the contribution of peptide structuring to these measurements, particularly regarding the ΔH value, as it reflects the overall bonding energy of all components within the complex. Nevertheless, these data clearly demonstrate a substantial gain in stability of BacA*_ef_* upon binding with the bacteriocin with a gradual effect with respect to the sequential binding of EntC peptides.

**Fig. 4.**
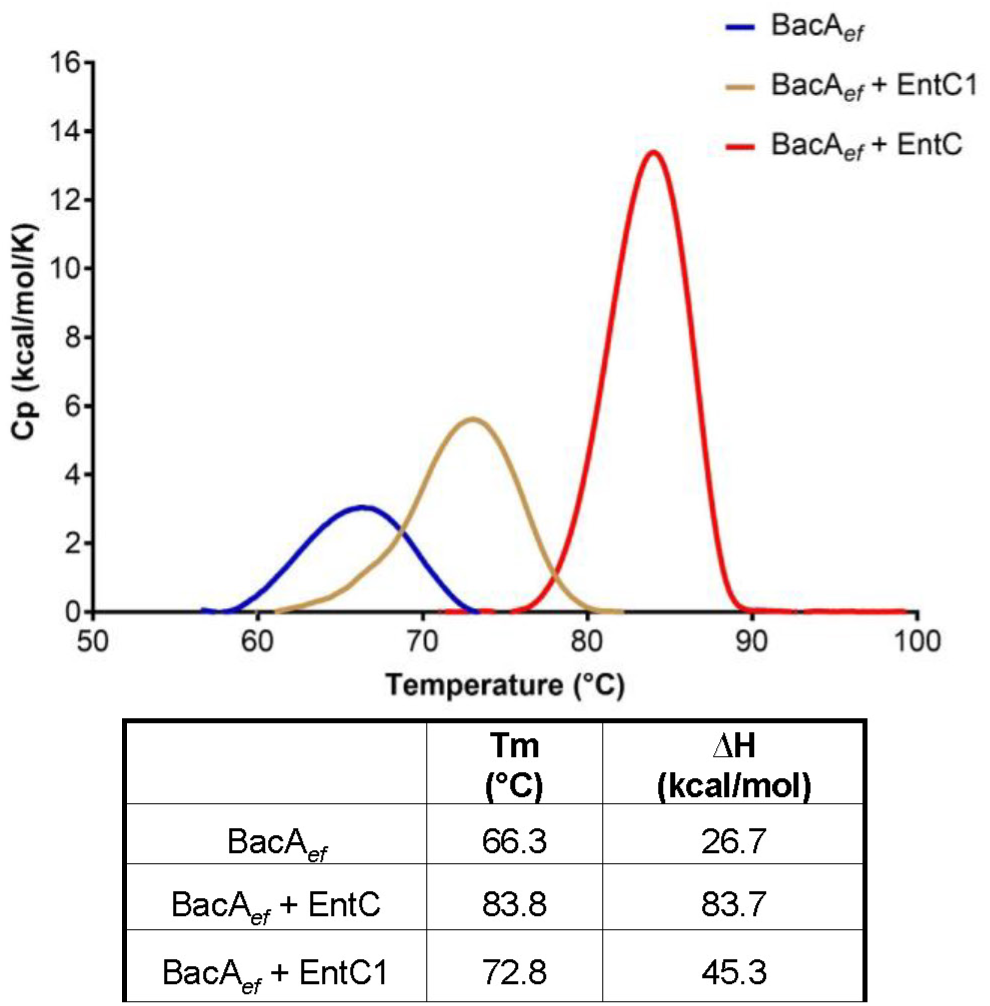
Stability enhancement of BacA*_ef_* by EntC peptides. Upper panel, representative DSC thermograms of BacA*_ef_* (32 µM) in the absence and the presence of EntC peptides (64 µM). Lower table, Tm and thermodynamic parameters. DSC scan were performed at 60°C/h between 30 °C and 100 °C. The data shown are representative of experiments performed in triplicate.

### AlphaFold2 models a reliable BacA:EntC complex

We used AlphaFold2 multimer (Evans *et al*, 2021) to predict the binding mode of EntC peptides to BacA*_ef_*. First, a model of BacA*_ef_* was generated with high confidence (**Appendix Fig. S6A,B**): the predicted template modelling (pTm) score was 0.92 and the predicted Local Distance Difference Test (pLDDT) score was 95.52 (**Appendix Fig. S6C**). The BacA*_ef_* model shares high similarity with the BacA*_ec_* crystallographic structure, displaying an RMSD of 1.35 Å over all C_α_ atoms (**Appendix Fig. S6D)**. According to this model, BacA*_ef_* consists of ten membrane-embedded α-helices, six of which are transmembrane (H3-5, H8-10), while shorter helices H1 and H2, as well as H6 and H7, form two reentrant helix-loop-helix motifs arranged in an inverted manner with respect to the membrane plane. The two short loops from these motifs come in close contact at the middle of the protein, where they form the bottom of a cavity that is open to the outer side of the membrane as well as towards the hydrophobic core of the lipid bilayer between TM helices H4 and H8 forming a V-shape entrance, which is lined by hydrophobic residues. Previous studies of BacA*_ec_* have identified E21, S27, and R174 as the catalytic residues, which, together with other conserved residues, are located within the helix-loop-helix motifs, at the bottom of the cavity (El Ghachi *et al*, 2018; Manat *et al*, 2015). Sequence alignment and structure superposition identified the corresponding catalytic residues in BacA*_ef_* as E20, S26, and R172 (**Fig**. **EV3**).

The structure of EntC peptides were not confidently predicted, either individually or combined (**Appendix Fig. S7**). In contrast, AlphaFold2 has confidently predicted a BacA*_ef_*:EntC1 complex (**Fig. 5A and Appendix Fig. S8A, B**) with an average pLDDT of 91.06, a pTM score of 0.89, and an ipTM (interface pTM score) score of 0.82. The confidence was only reduced for the N-terminal part of EntC1, which was not predicted to be in contact with BacA. According to this model, EntC1 consists of three short α-helices h1 V_3_-V_11_, h2 G_12_-M_25_ and h3 D_27_-K_38_ (**Fig. 5A**). The h2 and h3 helices are docking deeply within the outward-open cavity of BacA. The helix h2 is passing through the V-shaped entrance establishing hydrophobic contacts with TM helices H4 and H8 where they are both kinked at invariant proline residues. The helix h1 and a portion of h2 then protrude out of BacA cavity at the midplane of the membrane (**Fig. 5**). The h2 and h3 helices form a right angle close to BacA catalytic site, where EntC1-D27 forms hydrogen bonds with R172 and S26 catalytic residues of BacA (**Fig**. **6A**). Other charged residues form the C-terminus of EntC1 form hydrogen bonds and salt bridges with BacA residues from TM H3 (E47), H4 (D106 and D110) and H8 (K209) that are exposed at the upper side of the cavity entrance (**Fig**. **6B** **and Appendix Table S1**).

**Fig. 5.**
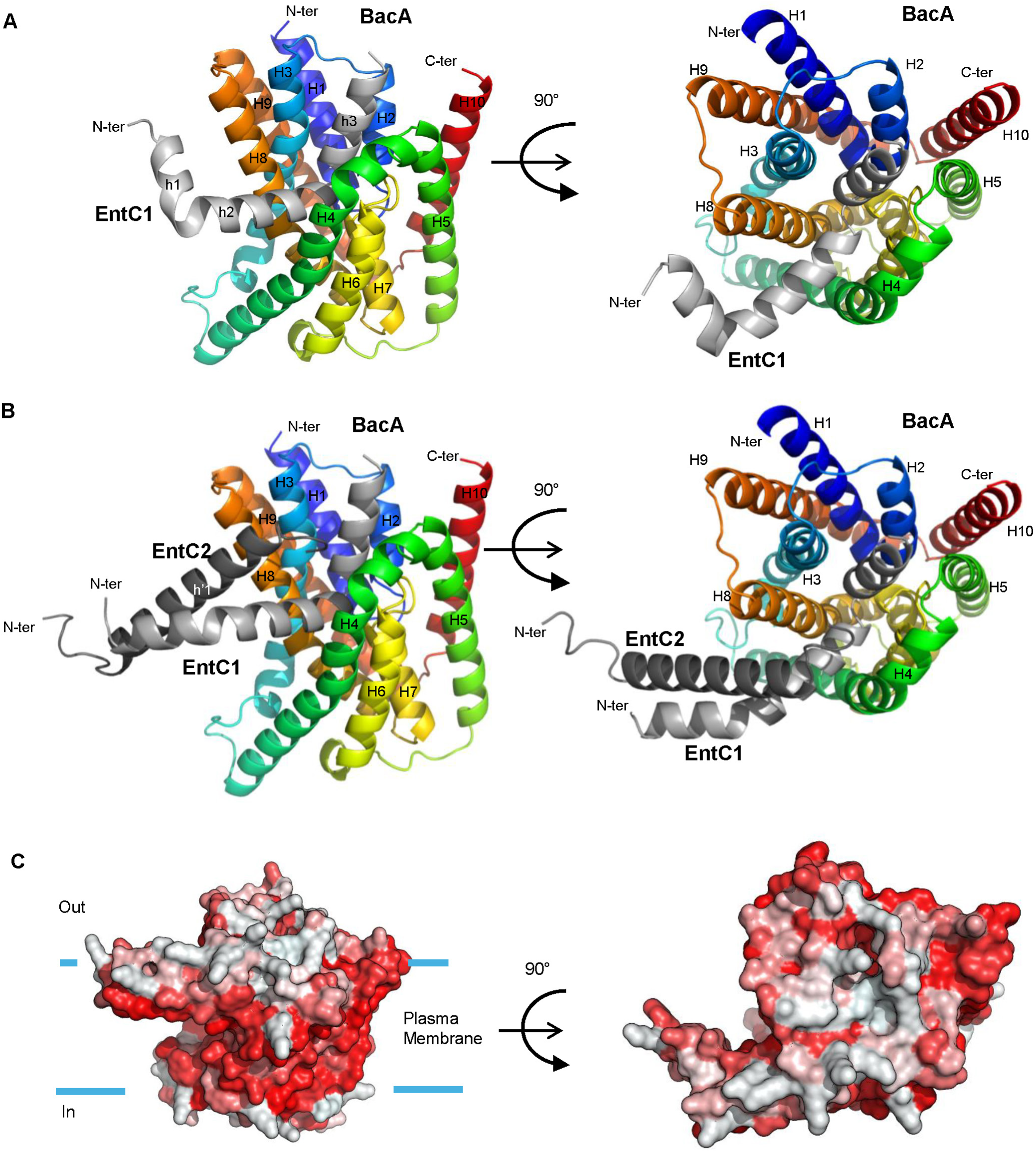
AlphaFold2 modelling of BacA*_ef_* in complex with EntC peptides. (**A,B**) Cartoon representation of the best structural model of BacA*_ef_* in complex with EntC1 (**A**) and with EntC (**B**). BacA*_ef_* is colored in rainbow from the N- (blue) to the C-terminal (red), EntC1 is colored in light grey, EntC2 is colored in grey. (**C**) Surface representation of the complex with hydrophobic residues according to the Eisenberg hydrophobicity scale colored in red. The likely membrane boundaries are indicated by blue bars.

**Fig. 6.**
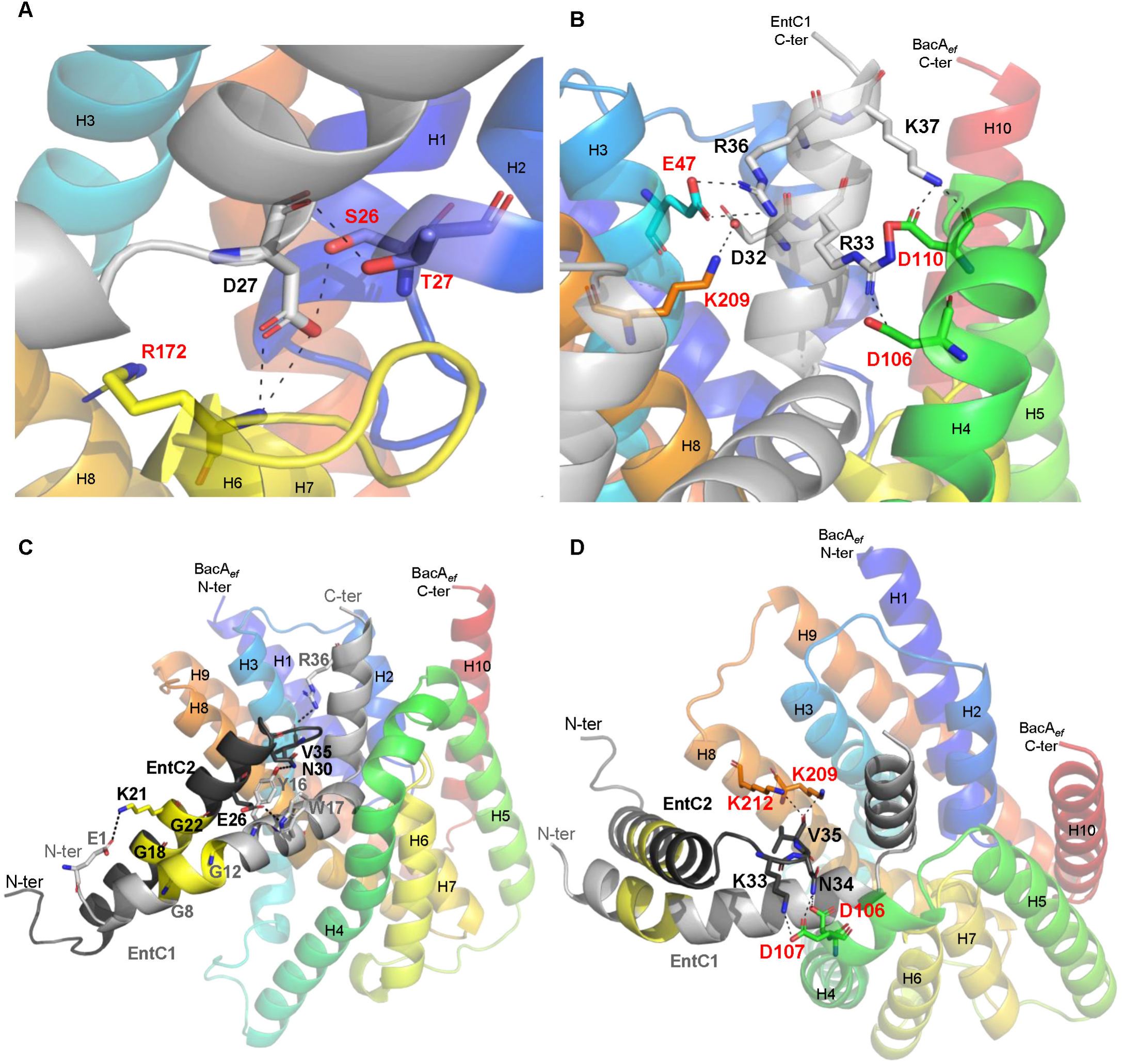
Details on Interfacial regions within AlphaFold2 models of BacA*_ef_* in complex with EntC peptides. (**A, B)** Zooms at the interface region between BacA*_ef_* and EntC1. (**A**) Zoom at the interface region nearby the catalytic site of BacA. (**B**) Zoom at the interface region at the upper-side entrance of BacA catalytic pocket. **(C, D)** Details of the interface regions formed between EntC2 and EntC1 (**C**) or BacA*_ef_* (**D**). Residues involved in hydrogen bonds or salt bridges (dashed lines), according to PISA analysis, are represented in stick and colored according to the atom (N in blue, O in red).

AlphaFold2 did not predict a complex between BacA*_ef_* and EntC2. In contrast, a complex between BacA*_ef_* and both EntC peptides simultaneously was modelled with a high pLDDT score of 94.44, a pTm score of 0.91, and an ipTM score of 0.89 (**Fig**. **5B, C** **and Appendix Fig. S8C**). In contrast to BacA*_ef_*-EntC1 model, the N-terminal part of EntC1 (helices h1 and h2), which is depicted to interact with the cognate EntC2 peptide, was modelled as an extended α-helix with a much greater confidence throughout the helix (pLDDT > 90). The C-terminal part of EntC1 remains in the same position within BacA cavity upon EntC2 binding. EntC2 forms a single α-helix (h1’ P_7_-N_30_) and only its non-structured N-terminal part, Gly1 to Trp5, displayed a lower confidence. There is no major structural difference between BacA*_ef_* in complex with EntC1 or both peptides (RMSD = 0.138 Å). According to this model, EntC2 interacts with BacA via its C-terminal K33, N34 and V35 residues forming hydrogen bonds and salt bridges with side chains of residues located at the outer ends of TM H4 (D106 and D107) and H8 (K209 and K212), as well as hydrophobic contacts between EntC2 A20, I23 and W32 with TM H8 F207, F211 and A215 at the entrance of the cavity (**Fig. 6C, D and Appendix Table S1**). A GxxxG motif is conserved in both peptides of all known class IIb bacteriocins (Oppegård *et al*, 2008; Nissen-Meyer *et al*, 2010) (**Fig. 1A**). These motifs were demonstrated to be essential for the antibacterial activity of the bacteriocins, where they were hypothesized to drive tight peptide-peptide interaction (Senes *et al*, 2000, 2004). In line with this, the G_8_xxxG_12_ motif from EntC1 is in very close contact with the G_18_xxxG_22_ motif from EntC2 (**Fig. 6C, D**). The N-terminal to mid-region of EntC peptides, featured by this tight helix-helix contact, are likely deeply embedded in the membrane hydrophobic acyl core, which is relevant with their amphipathic nature as emphasized by helical wheel projections and the calculation of their hydrophobic moment by Heliquest software (**Appendix Fig. S9**). According to the model, EntC binding to BacA induced only a minor change in the positioning of the BacA α-helices without significant restructuring (**Appendix Fig. S10**) (RMSD = 0.538 Å). The most notable change is a small shift of the ends of TM α-helices that are exposed to the outer side of the membrane leading to a small dilation of the aperture, but the protein cavity remains sealed towards the cytoplasmic side of the membrane.

### Specific BacA residues determine EntC binding

To assess the requirement for an active BacA for EntC activity and to highlight key residues involved in the formation of a complex, we generated BacA catalytic inactive variants through alanine substitution of E20, S26 and R172 residues. The variants were assessed by a functional complementation assay using the *E. coli* BWTetra-Ts*bacA* strain, which lacks all chromosomal genes encoding C_55_-PP phosphatase and carries a copy of *bacA_ec_* on a thermosensitive plasmid (Manat *et al*, 2015). This strain grows normally at 30 °C but lyses at 42 °C due to the arrest of C_55_-P recycling unless it is complemented *in trans*. The growth of this strain was fully restored at 42 °C upon the expression of *bacA_ef_* from a plasmid, while the mutation of any catalytic residue invalidated the complementation as expected (**Appendix Table S2**). The susceptibility of the *E. faecalis bacA* null mutant, VE18740 strain, to EntC was restored with a plasmid expressing wild-type *bacA_ef_* or any of the three catalytic site mutant alleles, although a slight decrease of the susceptibility was observed with R172A variant as judged from spot-on-lawn assays (**Fig. EV4A**). In conclusion, BacA functionality is not required for EntC antibacterial activity and although S26 and R172 are, according to AlphaFold2, involved in the contact, their mutation do not abolish BacA targeting. We also measured the bactericidal effect of EntC on VE18740 cells expressing the latter variants. When applied for 30 min on exponentially growing cells, 0.5 µM of EntC was found to kill more than 99% of the cells expressing wild-type BacA*_ef_* (**Table 2**). The expression of the variant E20A did not significantly change the susceptibility to EntC as compared to the wild-type, while the expression of S26A and R172A variants significantly increased the viability, up to 6.8 % and 7.8 %, respectively. The BacA*_ef_* variants were purified to further assess EntC binding. As anticipated, these variants were almost completely lacking any C_55_-PP phosphatase activity (**Appendix Table S2**). Titrations of EntC to BacA*_ef_*E20A displayed a very similar binding pattern as that determined with wild-type BacA. In contrast, consistently with the model, the titration of S26A and R172A variants displayed a gradual effect with an increase of the apparent *K*_d_, a decrease in the heat release upon binding, and a reduction in the entropy penalty (**Table 1 and Fig. EV4B**).

**Table 2.**
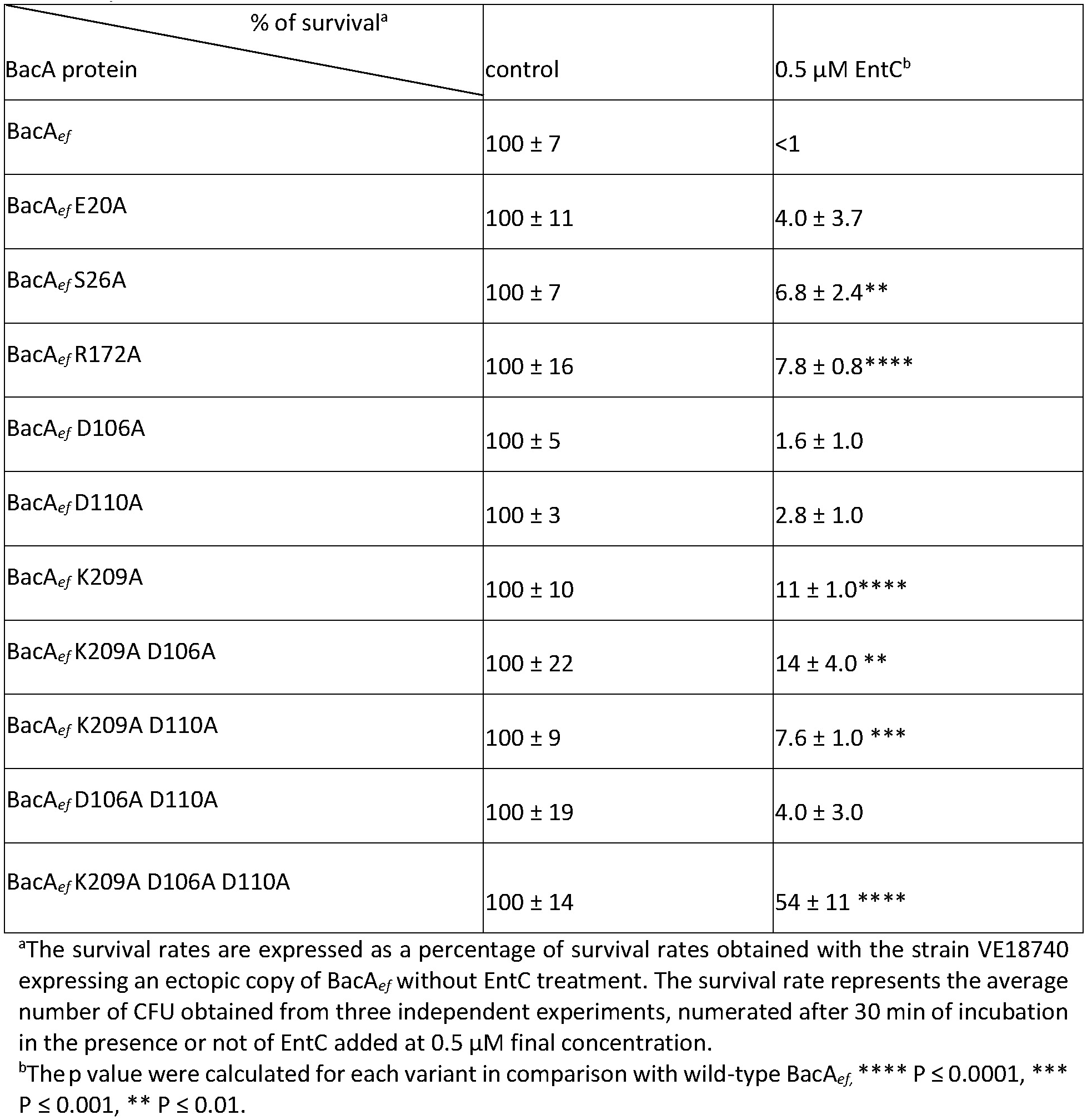
Survival rates of *E. faecalis* VE18740 strain (Δ*bacA*) expressing BacA variants upon incubation with 0.5 µM EntC for 30 min.

We also performed alanine substitution of D106, D110 and K209 residues of BacA*_ef_* since they were found to be involved in contacts with both EntC1 and EntC2 (**Appendix Table S1**). In contrast to the catalytic mutants, all the variants tested (combinations of single, double and triple mutations) complemented the *E. coli* BWTetra-Ts*bacA* strain, demonstrating the functionality of the variants (**Appendix Table S2**). Upon their expression in VE18740 strain, only the K209A variants caused a small reduction of susceptibility as compared to the wild-type BacA as judged from spot-on-lawn assays (**Fig. EV5A)**. When we measured the bactericidal effect of EntC on BacA variants-expressing cells, the K209A mutation resulted in a significant gain of survival rate, up to 11 % as compared to the wild-type, while the D106A and D110A mutations did not (**Table 2**). The double mutants (K209A-D106A and K209A-D110A) did not significantly increase survival rates as compared to K209A variant, while the triple mutant (K209A-D106A-D110A) displayed 54% of survival rate (**Table 2**). The BacA*_ef_*K209A and BacA*_ef_* triple mutants were purified. They both displayed 10% of residual C_55_-PP phosphatase activity (**Appendix Table S2**), demonstrating a significant role of K209 residue in BacA activity, although this residual activity was sufficient for *in vivo* complementation. EntC titration on BacA*_ef_*K209A revealed an impaired interaction characterized by a higher *K*_d_ and a reduction of heat content release, and no binding was observed upon titration of EntC on the triple mutant (**Table 1 and Fig. EV5B**). Since the lack of interaction could originate from an impaired stability of the latter variant, its thermal stability was compared to wild-type BacA using DSC (**Fig. EV5C**). The higher Tm and similar ΔH values of the variant as compared to the wild-type BacA indicated that the lack of interaction was not due to impaired stability.

### Cross-reactivity of EntC-related bacteriocins

To date, only a limited number of EntC-related bacteriocins have been identified, they are categorized as Enterocins and Lactococcins based on the producing cells (**Fig. 1A**). To investigate their specificity of action, we examined the cross-reactivity of LcnG and EntC peptides. Chemically synthesized LcnG peptides (61% identity with EntC) were obtained. *L. lactis* IL1403 exhibited sensitivity to both bacteriocins, while *E. faecalis* was only susceptible to EntC (**Fig**. **7A**). Alignment of BacA from these two species revealed conservation of residues involved in EntC:BacA interaction according to AlphaFold2 (**Fig. EV3 and Appendix Table S1**). *In vitro*, LcnG inhibited BacA*_ef_* C_55_-PP phosphatase activity in a dose-dependent manner similar to EntC (IC_50_ value of LcnG measured at 0.8 µM) (**Fig**. **7B**). LcnG_α_ (EntC1 homologue) also inhibited BacA*_ef_* activity with a comparable IC_50_ value (1 µM), while LcnG_β_ had no effect. Unfortunately, we were unable to produce a stable form of BacA from *L. lactis* for a comprehensive *in vitro* comparative study.

**Fig. 7.**
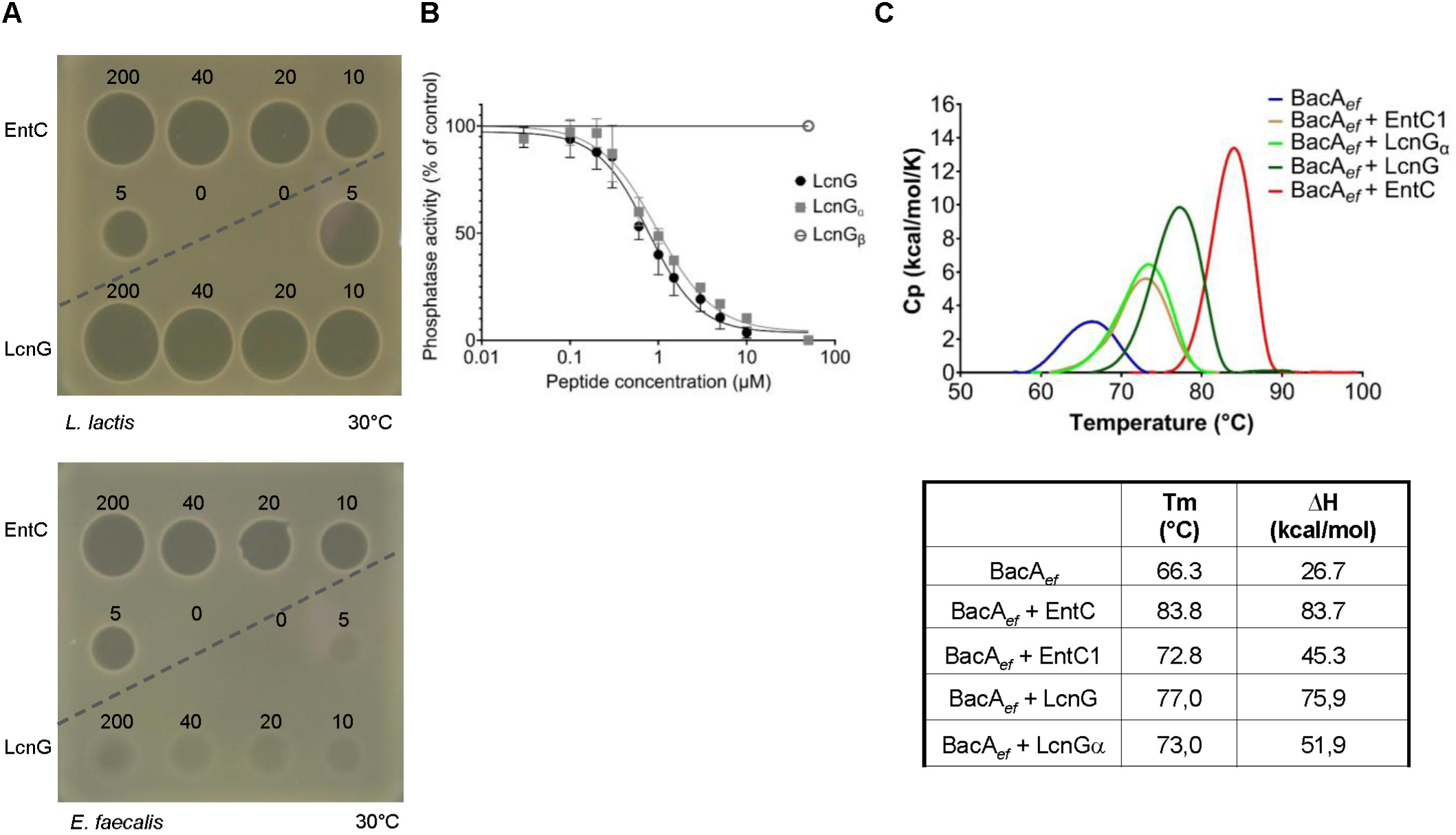
Cross-activity of EntC and LcnG peptides. (**A**) Activity measurement of EntC and LcnG peptides towards *L. lactis* and *E. faecalis* cells. (**B**) Dose effect of LcnG peptides on the C_55_-PP phosphatase activity of BacA*_ef_* in detergent. Results are the average of three independent experiments and error bars indicate standard deviations. (**C**) Stability measurement of BacA*_ef_* in the absence or the presence of different peptide solutions. Upper panel, representative DSC thermograms of BacA*_ef_* (32 µM) incubated with different peptide mixtures (64 µM), lower table, T_m_ and thermodynamic parameters.

We further assessed LcnG binding to BacA*_ef_* by ITC (**Table 1 and Appendix Fig. S11**). LcnG titration on BacA*_ef_* exhibited an exothermic reaction with thermodynamic parameters similar to EntC titration, characterized by a substantial enthalpy release (−20.4 kcal/mol) and a large entropy penalty (11.3 kcal/mol), and a *K*_d_ of 165 nM. LcnG_α_ titration exhibited a significantly higher *K*_d_ compared to the complete bacteriocin (898 nM). As anticipated, LcnG_β_ titration showed no interaction with BacA. Titration of LcnG_β_ on the BacA*_ef_* + LcnG_α_ mixture displayed a strong enthalpy-driven interaction (ΔH of −16.8 kcal/mol), with an entropy penalty (-TΔS of 7.4 kcal/mol) and the *K*_d_ was lower (106 nM) as compared to LcnG_α_. Overall, these results demonstrate a binding pattern of LcnG to BacA*_ef_* similar to EntC, although LcnG was found to be inactive against *E. faecalis*.

We further assessed the structural similarity of the complexes formed by BacA*_ef_* with both bacteriocins by DSC (**Fig**. **7C**). The addition of LcnG_α_ to BacA*_ef_* resulted in an increase in BacA T_m_ and ΔH in a manner analogous to EntC1. The presence of both LcnG peptides further increased T_m_ and ΔH values, albeit not to the same extent as EntC. Despite LcnG binds to BacA with similar thermodynamic parameters as EntC, thermal stability measurements revealed significant differences between the complexes they form, with differences of 6.8°C (T_m_) and 7.8 kcal/mol (ΔH). AlphaFold2 generated a high-confidence model of BacA*_ef_* in complex with LcnG, exhibiting an average pLDDT of 95.04, a pTM score of 0.92, and an ipTM of 0.9 (**Appendix Fig. S12**). This model indicates that LcnG is positioned in a nearly identical manner as to EntC in the BacA outward-open cavity, with an RMSD of 0.2 Å. A notable difference between the models of the complexes, is the lack of an ionic bond between LcnG_α_ and K209 residue from BacA*_ef_* due to the lack of an acidic residue in LcnG_α_ as compared to D32 from EntC1 (**Appendix Fig. S13**).

### Activity of chimera bacteriocins

To elucidate the determinants for the specificity of these bacteriocins, chimera bacteriocins were evaluated for activity. The LcnG_α_-EntC2 chimera (LcnG_α_ and EntC2 in 1:1 molar ratio) exhibited no activity against either *E. faecalis* or *L. lactis* (**Fig. 8A**) suggesting that EntC2 either cannot bind to BacA:LcnG_α_ pre-complex (regardless of BacA origin) or does not form an active complex. Notably, this peptide combination lacks the interaction between residue E1 of EntC1 (lacking in LcnG_α_) and K21 of EntC2 (**Fig. 1A and Fig. 6C**). Previous mutational studies have demonstrated that residue E1 in Ent1071A (Ent1071 is a close homologue of EntC) is essential for the bacteriocin’s activity. Moreover, the absence of a negative charge at position 1 in peptide LcnG_α_ has been shown to impair activity when paired with EntC1071B (Oppegård *et al*, 2007). Conversely, the EntC1-LcnG_β_ chimera exhibited a similar antibacterial activity to EntC and LcnG against *L. lactis*, and it also displayed a moderate activity against *E. faecalis* in contrast to LcnG, which lacks activity against this bacterium. This suggests that LcnG_β_ can interact with the BacA:EntC1 pre-complex in a cytotoxic manner as evidenced by *L. lactis* susceptibility. We analyzed the binding of the chimeras to BacA*_ef_* using ITC (**Table 1 and Appendix Fig. S14**).

**Fig. 8.**
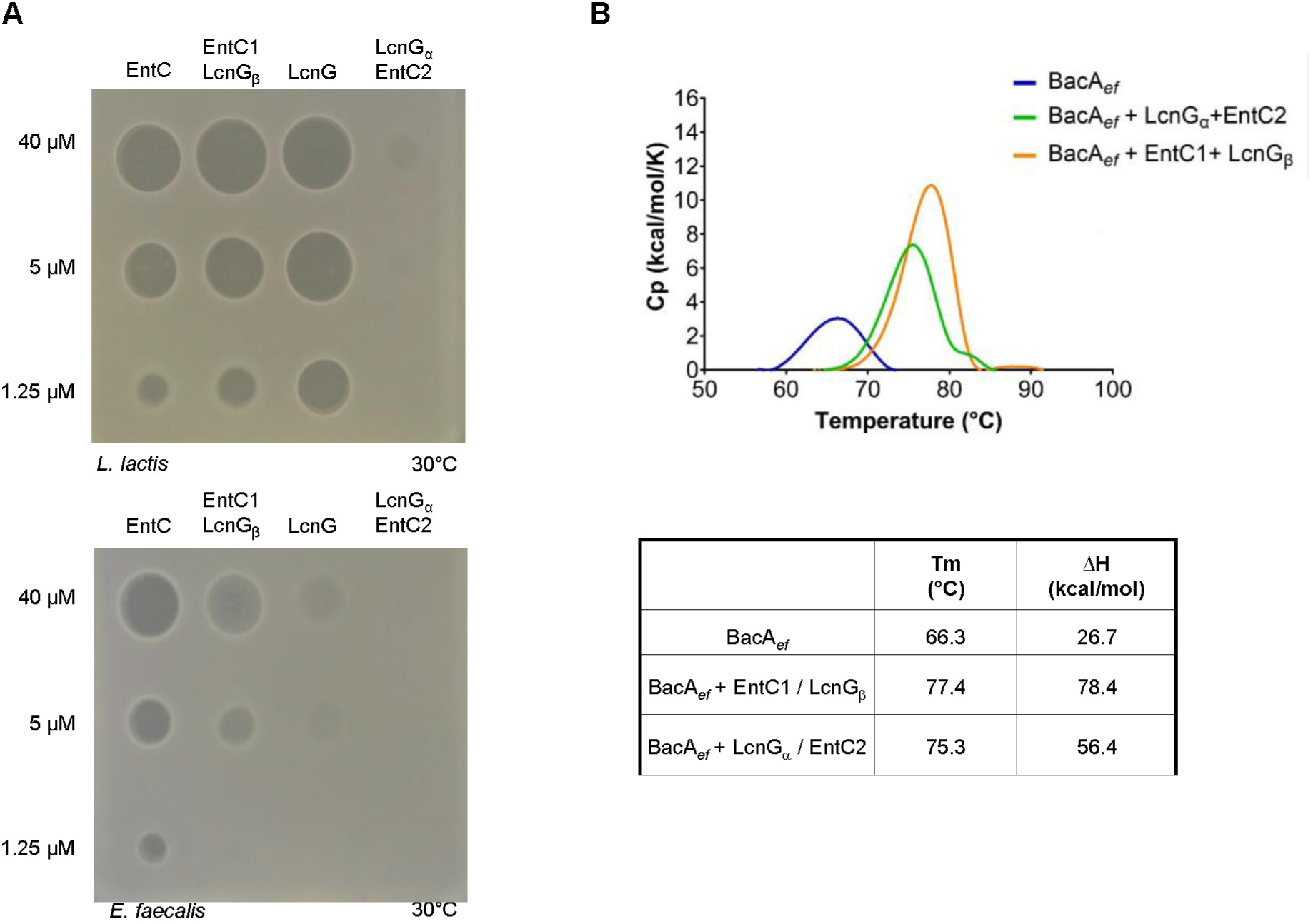
Activity of chimeric bacteriocins. (**A**) Activity measurement of EntC and LcnG chimera on *L. lactis* (upper panel) and *E. faecalis* (lower panel) strains. The data shown are representative of experiments performed in triplicate. (**B**) Stability measurement of BacA*_ef_* in the absence or in the presence of bacteriocin chimera. Upper panel, representative DSC thermograms of BacA*_ef_* (32 µM) incubated with chimera (64 µM), lower table, T_m_ and thermodynamic parameters.

The titration of the inactive LcnG_α_-EntC2 chimera on BacA*_ef_*, or titration of EntC2 to a BacA*_ef_* + LcnG_α_ mixture, both exhibited exothermic binding curves, displaying similar *K*_d_ and thermodynamic parameters to native bacteriocins. Similarly, the titration of EntC1-LcnG_β_ on BacA*_ef_*, or titration of LcnG_β_ to a BacA*_ef_* + EntC1 mixture, demonstrated exothermic binding curves. These findings demonstrate that binding of the peptides to BacA is not the sole critical factor, but the conformation or dynamics of the tripartite complexes within the membrane, especially the protruding parts of the bacteriocins, likely plays a key role in their antibacterial activities.

We investigated the resulting complex conformation by measuring the T_m_ and enthalpy of denaturation (**Fig**. **8B**). The addition of the inactive LcnG_α_-EntC2 chimera to BacA*_ef_* increased BacA*_ef_’s* T_m_ and ΔH values of by 9°C and 29.7 kcal/mol, respectively, similar to the effect of LcnG_α_ on BacA. The EntC1-LcnG_β_ chimera had a more pronounced effect on BacA*_ef_*, raising T_m_ and ΔH values by 11.1°C and 51.7 kcal/mol, comparable to LcnG’s impact on BacA. These results infer distinct conformational states or dynamics induced by the different combinations. EntC1 and LcnG_α_ exhibited the least effect, while EntC caused the most significant stabilization, which correlates with their toxic activity against *E. faecalis*.

## Discussion

In the present work, we have demonstrated that EntC, and the related bacteriocin LcnG, specifically target the C_55_-P recycling protein BacA through a direct and high-affinity binding. This binding event elicits the dissipation of proton gradient as observed when EntC was added to BacA*_ef_*-containing liposomes, supporting the hypothesis that the membrane permeabilization is the primary cause for the death of the target cells. It further demonstrates that BacA is required and sufficient for EntC to trigger membrane disruption. AlphaFold modelling and interaction assays demonstrate that EntC1 initiates the binding by docking to the outward-open cavity of BacA, establishing intermolecular bonds with deeper BacA catalytic residues. This binding mode of EntC1 to BacA likely mimics the mode of access of C_55_-PP substrate from the outer leaflet of the lipid bilayer. The negatively charged headgroup of C_55_-PP must interact with the active site residues, especially with positively charged R174, localized at the bottom of the catalytic pocket, which corresponds to the midplane of the membrane. The C_55_-PP lipid tail should then exit the cavity along the hydrophobic groove shaped by TM helices H4 and H8, and protrude dynamically within the lipid bilayer. Although, a complex of BacA with its substrate has not yet been resolved, this mode of substrate binding was strongly supported by the presence of monoolein lipid, presumably mimicking C_55_-PP, within the active site pocket in the crystal structure of BacA*_ec_* (Workman *et al*, 2018). According to its binding site, EntC1 binding must then block substrate access and thus inhibit BacA catalysis as it was herein observed experimentally. ITC measurements of EntC1 binding to BacA reveal an entropy-driven binding event, suggesting the release of ordered water molecules from the binding interface and/or the maintenance of conformational flexibility within the bipartite complex. This aligns with the modelled position of EntC1 within BacA’s cavity, where the binding of EntC1 to the hydrophobic pass leading to the cavity would promote the release of ordered water molecules, contributing to the overall increase of the entropy (Mosebi, 2022; Teilum *et al*, 2009; Krimmer & Klebe, 2015). Moreover, according to our model, the N-terminal to mid-region of EntC1 is protruding out of BacA within the hydrophobic core of the lipid bilayer, likely conferring a high degree of freedom to this region (Teilum *et al*, 2009).

As demonstrated by ITC, the binding of EntC1 enables the subsequent docking of EntC2 to BacA. This observation is consistent with AlphaFold2 modelling, which indicates that EntC2 interacts peripherally in the tripartite complex, establishing contacts with the protruding N-terminal to mid-region of EntC1 as well as with residues from BacA positioned at the ends of TM helices forming the V-shape opening towards the catalytic pocket. This second binding event is characterized by a substantial heat release and a significant entropy penalty. The close helix-helix contacts between the two peptides may significantly reduce the flexibility of EntC1, potentially accounting for the entropy cost and for the strong stabilization of the tripartite complex observed by DSC. It is also plausible that the peptides may gain structuring as they interact with BacA, which may contribute to these enthalpy-driven binding events.

According to the model, the binding of EntC peptides with BacA involves their respective mid-region to C-terminus (residues 20-39 in EntC1 and 21-35 in EntC2), which exhibit high sequence similarity between the related bacteriocins (c.a. 80% identity), while their N-terminal halves are more divergent (c.a. 40% identity) (**Fig. 1A**). This observation aligns with the fact that EntC and LcnG peptides, as well as any chimeric combinations, were found to interact with BacA_ef_ yielding similar thermodynamic parameters. Nevertheless, EntC is the only bacteriocin displaying antibacterial activity against *E. faecalis,* suggesting that beyond their similar docking to BacA, the conformation or dynamics of the tripartite complex, particularly the tight helix-helix motif of the bacteriocin that protrudes within the lipid bilayer, is critical. This likely accounts for the lack of consistency between BacA binding and antibacterial activity. The reason for LcnG’s inactivity against *E. faecalis* remains unclear, considering that the docking to BacA*_ef_* is conserved (except for a salt bridge between the peptide and BacA*_ef_* K209 residue) and that the N-terminal parts of the two peptides are compatible by nature. These results suggest that the membrane features may be critical for the activity of the bacteriocins and that the composition or dynamics may differ between the membranes of *E. faecalis* and *L. lactis,* potentially accounting for the observed discrepancy. Obviously, the N-terminal halves of the peptides have evolved more rapidly than their C-terminal counterparts, which is consistent with the fact that their evolution is not constrained by binding to the cell surface receptor. Consequently, cognate peptides have co-evolved in their N-terminal halves independently to BacA (**Fig. 1A**). Evidently, the docking of the peptides via their C-terminal halves does not confer the killing activities *per se*, since similar docking patterns do not necessarily result in antibacterial activities, as exemplified by the lack of activity of LcnG or chimera against *E. faecalis*. Therefore, their protruding halves, whose folding does not strictly depend on BacA, may be pivotal for the killing activity.

The cognate peptides interact primarily via their respective GxxxG sequences, which have been previously identified as essential for the antibacterial activity of all class IIb bacteriocins (Nissen-Meyer *et al*, 2011), despite the fact that these bacteriocins do not apparently share a common cell surface receptor. This further supports the hypothesis that this common helix-helix motif of class IIb bacteriocins is not directly involved in the recognition of the receptor but rather interacts with the membrane to facilitate membrane permeabilization. In EntC/LcnG bacteriocins, this tight helix-helix motif protrudes deeply in the lipid bilayer due to the docking of the C-terminal parts of the peptides to the outward-open cavity of BacA, passing via the V-shaped entrance leading to the acyl core of the membrane. The docking of the peptides to BacA likely provides the driving force required for the peptides to insert deeply within the lipid bilayer and cause membrane permeabilization. These GxxxG motifs constitute a prevalent transmembrane helical dimerization interface in which the C_α_-H from two glycine residues from one helix form hydrogen bonds with the carbonyl from the nearby helix, enabling very close contact of the membrane-embedded helices with minimal steric hindrance (Teese & Langosch, 2015). Recent studies have highlighted the role of GxxxG motifs in the toxicity and pore formation activity of amyloid peptides through the strong stabilization of ion-channel-like pores (Rando *et al*, 2023). Notably, the docking of EntC provides substantial stabilization of the tripartite complex and the degree of stabilization was found to correlate with the level of antibacterial activity when comparing the effect of native and chimera bacteriocins. The conformation of a tripartite complex with the least degree of freedom in the membrane apparently provides the optimal support for membrane permeabilization. As previously emphasized, a negative charge at position 1 in EntC1 was shown to be essential for EntC activity, and the lack of equivalent residue in LcnG_α_ accounted for the lack of activity when paired with Ent1071B (equivalent to EntC2) (Oppegård *et al*, 2007). This observation aligns with the lack of antibacterial activity of LcnG_α_-EntC2 chimera. Interestingly, LcnG_β_ has a negative charge at position 10 that is apparently functionally equivalent to E1 from EntC1, as the detrimental effect of removing E1 from EntC1 was neutralized by introducing a negative charge at position 10 in Ent1071B (Oppegård *et al*, 2007). In our model, E1 from EntC1 is forming an ionic bond with K21 from EntC2, and position 10 in EntC2 is in very close proximity to E1 (**Fig. 6C**), suggesting that introducing a negative charge at this position could establish the same ionic pairing with K21, thereby compensating the removal of E1. Since this region of the peptides is expected to reside in the acyl core of the bilayer, we can hypothesize that this ionic bond is essential to maintain and stabilize the peptides anchored to the bilayer through the neutralization of this charged residues. These charged residues positioned within the acyl core may also facilitate proton leakage across the membrane. Overall, these observations support a key role of the N-terminal halves of the peptides in membrane destabilization. As demonstrated herein, the peptide EntC1 directly interacts with BacA*_ef_* and EntC2 subsequently binds to the BacA:EntC1 bipartite complex. However, several evidence suggests that both peptides may initially interact when they come into contact with the membrane surface, prior to their docking to BacA, although this peptide-peptide interaction could not be elucidated through biophysical approaches. Due to the small size of the peptides, their interaction is challenging to measure using ITC or FIDA, as these techniques lack the sensitivity to detect the likely low enthalpy change and minimal molecular size variation upon interaction. The prior interaction of the peptides with each other is supported by the lower *K*_d_ values measured when titrating EntC or LcnG to BacA*_ef_* as compared to EntC1 and LcnG_α_, as well as the lower IC_50_ value measured for the inhibition of BacA*_ef_* activity by EntC as compared to EntC1. Furthermore, this reasoning aligns with previous NMR and circular dichroism (CD) spectroscopy studies of LcnG peptides. These studies have demonstrated that both LcnG peptides are unstructured in water, and become structured, displaying helical content primarily in their N-terminal halves, when exposed separately to membrane-like environments such as micelles and liposomes. CD-spectroscopy studies then revealed a significant enhancement of their helical content when both peptides were combined with the membrane-like environment, strongly suggesting that the peptides interact to form one functional unit (Rogne *et al*, 2008; Nissen-Meyer *et al*, 2010; Hauge *et al*, 1998). The interaction of the peptides together at the contact of the membrane surface would then enable a single binding event with BacA through lateral diffusion at the membrane surface rather than relying on diffusion in the aqueous environment and sequential interactions of the peptides with their receptor. This mode of action may be critical for their antibacterial activity by enhancing the rate of binding to the cell surface receptor via the reduction of the dimensionality that has been featured as membrane catalysis to overcome the diffusion control barrier (Castanho & Fernandes, 2006). While peptide-peptide interaction and the cooperative structuring of the peptides at the contact of the membrane warrant further investigation, it is evidently insufficient to mediate the permeabilization of the membrane. The subsequent docking of the bacteriocin to BacA likely enable deeper insertion of the peptides in the membrane, which must be pivotal for membrane disruption.

At present, a limited number of mechanisms for membrane permeabilization by bacteriocins have been elucidated. The lantibiotic Nisin functions by lipid II binding, resulting in the formation of water-filled pores in a barrel stave model, wherein peptides are oriented perpendicular to the membrane plane (Moll *et al*, 1996b; Héchard & Sahl, 2002). An alternative mode of action, applicable to certain lantibiotics, is the wedge model (Driessen *et al*, 1995; Li *et al*, 2021). In this model, an interaction occurs between the cationic part of the bacteriocin with the anionic head groups of phospholipids, enabling the hydrophobic residues of the bacteriocin to insert into the membrane acyl core, causing local deformation of the lipid bilayer (Pérez-Ramos *et al*, 2021). For class II bacteriocins, the only known mechanism involves class IIa bacteriocins interacting with the transmembrane mannose phosphotransferase system (Man-PTS) protein. In the case of the pediocin PA-1, the bacteriocin penetrates the Man-PTS protein core domain. The N-terminal region interacts with the surface domain of the protein, while the C-terminal domain half penetrates the membrane through the protein. This interaction opens the Man-PTS in a wedge-like manner, forming a pore across the membrane leading to cell death (Zhu *et al*, 2022). In comparison to these mechanisms, the action of EntC and its homologues appears to adopt a distinct mode of action. Our findings indicate that the inhibition of BacA activity is not the primary cause of toxicity, since BacA is not essential in *E. faecalis* as demonstrated here, likely due to its redundancy with other C_55_-PP phosphatases and C_55_-P flippases. Instead, our findings strongly suggest that the lethal activity of these bacteriocins is attributable to membrane permeabilization. This is consistent with the high potency of these bacteriocins, as membrane permeabilization depends on a low number of peptides, unlike what is expected when cell death is promoted by the inhibition of a protein activity.

Given the amphipathic nature of the peptides and the mode of docking to BacA, EntC likely resides deep within the lipid bilayer but rather in a parallel orientation with respect to the membrane plane, without crossing the entire membrane, contrary to previous hypotheses (Nissen-Meyer *et al*, 2010). EntC insertion could subsequently induce a local thinning or deformation of the membrane, resulting in an imbalance of charge, area, and surface tension that may drive the local disruption of the bilayer structure and cause leakage. This mechanism was hypothesized to be the basis for the antibacterial activity of several amphipathic peptides that affect cell membrane integrity, known as interfacial activity model (Wimley, 2010; Grage *et al*, 2016). Our study provides novel mechanistic insights into bacteriocin-target interactions and identify BacA as a novel and vulnerable antibacterial target. These discoveries significantly advance our understanding of class II bacteriocins and open avenues for the rational design of peptide-based antimicrobials with enhanced specificity and potency.

## Methods

### Products

The oligonucleotides were provided and DNA sequencing was performed by Eurofins Genomics. *n*-dodecyl-β-D-maltopyranoside (DDM) was provided by ThermoScientific. Farnesyl pyrophosphate (C_15_-PP) and isopentenyl pyrophosphate (C_5_-PP) were from Sigma, and [^14^C]C_5_-PP ((1.48-2.22 GBq)/mmol) was from Perkin Elmer. DOPC, DOPE and DOPG lipids were provided by Avanti Polar Lipids. C_55_-PP synthase UppS was purified as previously described (El Ghachi *et al*, 2004). [^14^C]C_55_- PP was prepared from it precursors,C_15_-PP and [^14^C]C_5_-PP, upon reaction with UppS according to the published procedure (El Ghachi *et al*, 2004). DNA purification kits were from Macherey-Nagel. Peptides were synthesised by GenScript at purity level over 90 %. All other materials were of reagent grade and obtained from commercial sources. *E. coli* strains were grown in 2xYeast Extract and Tryptone (2YT) medium, at 30, 37 or 42°C with aeration and agitation (200 rpm). Solid media were obtained by adding 15 g/L agar. *E. faecalis* strains were grown in brain heart infusion (BHI) medium at 30 or 37 °C in static conditions, and when specified in M17 supplemented with 0.5% (w/v) glucose (M17G) or MRS. *L. lactis* strain Il1403 was grown in M17G at 30°C in static conditions. The antibiotics ampicillin (Amp) and chloramphenicol (Cm) were used at 100 µg/mL and 25 µg/mL, respectively. Erythromycin (Ery) was used at 3 µg/mL when used for *E. faecalis* and 150 µg/mL for *E. coli*.

### Deletion of *bacA_ef_*

The VE18740 strain (Δ*bacA_ef_*) was generated by double-crossover recombination in the VE18379 strain (Furlan *et al*, 2019) using pVE14414, a pGhost9-derivative plasmid (Maguin *et al*, 1992). Two DNA fragments encompassing the 5’- and 3’-ends of the *bacA_ef_* gene were amplified by PCR from VE18379 chromosomal DNA with primer pairs OEF1066-OEF1067 and OEF1068-OEF1069, respectively (**Appendix Table S3**). The fragments were then cloned by Gibson assembly into pGhost9 at the EcoRV site, yielding pVE14414. A markerless in-frame *bacA_ef_* deletion mutant was selected as previously described (Brinster *et al*, 2007), and the deletion of *bacA_ef_* was confirmed by sequencing of the genomic DNA of the strain.

### Cloning

For complementation experiments of VE18740 strain, *bacA_ef_* was cloned in a low copy number *E. coli*-*E. faecalis* shuttle plasmid. The entire gene was amplified using primers OEF1170-OEF1171 (**Appendix Table S3**) and cloned by Gibson assembly into pIL252 at the SmaI site, yielding plasmid pVE14417. To facilitate targeted mutagenesis, the p15A replication origin amplified with OEF1145-OEF1146 primers and digested with NheI restriction enzyme was introduced into pVE14417 at the XbaI site. The resulting pVE14420 plasmid was electroporated into VE18740 strain, yielding the VE18749 strain. VE18740 strain harboring empty pIL252 plasmid was used as a control. For recombinant expression of BacA in the *E. coli* C43(*DE3*) strain, the *bacA_ef_* gene was amplified using oligonucleotides BacA1-BacA2 (**Appendix Table S3**). The resulting fragment was inserted at BamHI and HindIII sites in pET2130 vector (derived of pET21d expression vector from Novagen, His_6_-Tag in N-ter, T7 promoter, Amp^R^) generating pBacA*_ef_*His. Plasmid pTrcBac30 was used for the expression of N-terminal His_6_-tagged BacA*_ec_* (El Ghachi *et al*, 2004), while plasmid pPgpBHis was used for the expression of N-terminal His_6_-tagged PgpB (Touzé *et al*, 2008). Variants were produced by site-directed mutagenesis using the QuickChange II Site-Directed Mutagenesis Kit (Agilent Technologies) with the original plasmid as a template and a pair of oligonucleotides (Eurofins Genomics) containing the mutation (**Appendix Table S3**). Mutations were confirmed by sequencing the entire *bacA* gene. Electrotransformation of *E. coli* and *E. faecalis* was carried out as previously described (Dower *et al*, 1988; Dunny *et al*, 1991).

### EntC or LcnG agar-assay plate on *E. faecalis* or *L. lactis*

VE18740 *E. faecalis* and IL1403 *L. lactis* strains were grown in BHI and M17G media, respectively. The selective antibiotic Ery was added if necessary (if strain carries a plasmid of interest). When absorbance at 600nm (A_600_) reached 0.6, the culture was diluted to 10⁶ CFU/mL in the corresponding media containing 0.75% agar. The mix was spread on an agar plate of the corresponding media and 6 µL-droplets of EntC or LcnG peptides solutions, at various concentrations, were spotted. The plates were incubated overnight at 30°C or 37°C.

### Survival test on *E. faecalis*

A culture at exponential growth (A_600_ ∼0.4, corresponding to about 3 x 10^8^ CFU/mL) of VE18740 strain transformed by the desired plasmid was diluted to 10^6^ CFU/mL in BHI in the presence or absence of EntC at 0.5 µM. After 30 min of incubation at 37°C, serial dilutions were plated on BHI-Ery agar plates and incubated overnight at 37°C to determine the viable cell counts. The rate of survival was expressed as a percentage of CFU obtained after exposure to EntC as compared with the control without EntC.

### Expression of recombinant Proteins

The His_6_-tagged proteins and variants were expressed in the *E. coli* C43(*DE3*) cells. The cells were grown in 2YT medium supplemented with ampicillin at 37°C until A_600_ reaches 1. Protein production was then induced by the addition of 1 mM IPTG and the culture was placed at 22 °C and rotated at 180 rpm, for 16 h. Cells were harvested by centrifugation at 4000 × *g* for 20 min at 4°C and washed by resuspension in buffer A (20 mM HEPES, pH 7.5, 0.4 M NaCl, 10% (v/v) glycerol). The bacterial pellet was stored at −20°C until protein purification.

### Purification of membrane proteins

Cells were resuspended in buffer A and lysed using a French press. The membrane fraction was obtained after ultra-centrifugation at 186,000 × *g* for 45 min at 4°C. The membrane pellet was resuspended in 20 ml of buffer B (buffer A + 2% (w/v) DDM) and incubated for 2 h with agitation at 4°C for solubilization, followed by centrifugation at 165,000 × *g* for 25 min at 4°C. One mL of Ni^2+^- nitrilotriacetic acid agarose resin pre-equilibrated in buffer C (buffer A + 0.1% DDM) was added to the supernatant, and the mixture was incubated overnight at 4°C under agitation. It was then transferred into a chromatography column Poly-prep® (Biorad) allowing the resin coupled with the proteins to settle, and was further washed with buffer C. The column was then washed with 30 mL of buffer C + 10 mM imidazole. Proteins were eluted in 1mL-fractions with buffer C containing 400 mM imidazole. Fractions were quantified by absorbance at 280 nm and analyzed on SDS-PAGE (12% acrylamide). Those containing the target protein were subjected to gel filtration on a Superdex 200 pg column in buffer D (buffer C with NaCl at 0.2 M) at 1 ml/min flow rate. 1 mL-fractions were collected, quantified by absorbance at 280 nm and analyzed by SDS-PAGE coupled Coomassie blue staining or western blotting using an anti-His tag antibody. Protein fractions were concentrated using a Vivaspin™ 20 concentrator with a 50 kDa cut-off up to 1 mg/ml as determined by the absorbance at 280 nm using extinction coefficients of 44 920 M^−1^ cm^−1^ and 25 440 M^−1^ cm^−1^ for BacA*_ef_* and BacA*_ec_*, respectively. Samples were stored at −20°C. ITC and DSC measurements were carried out with a BacA sample at 1 mg/ml dialyzed overnight against the following buffer: 20 mM Tris-HCl, pH 7.5, 0.2 M NaCl and 0.1% DDM.

### C_55_-PP phosphatase assay in detergent

The C_55_-PP phosphatase assays were performed in a 10-μL reaction mixture containing 20 mM Tris-HCl, pH 7.4, 400 µM CaCl_2_, 150 mM NaCl, 0.1 % (w/v) DDM and 50 μM radiolabeled [^14^C]C_55_-PP (900 Bq). The BacA concentration was adjusted in order to achieve less than 30 % substrate hydrolysis. The mixture was incubated for 10 min at 37 °C and the reaction was stopped by freezing in liquid nitrogen. Substrates and products were separated by thin layer chromatography on precoated silica gel 60 plates (Merck) using 1-propanol/ammonium hydroxide/water (6:3:1, v/v/v) as the mobile phase. Radioactive spots were located and quantified using a radioactivity scanner (model Multi-Tracermaster LB285, Berthold-France). When the phosphatase activity was tested at different pH values, the following buffers were used at 20 mM: sodium acetate (pH 4-7) or Tris-HCl (pH 7-8). In the EDTA assay, the protein was pre-treated with 200 µM EDTA. For pH and EDTA assays, CaCl_2_ was omitted from the reaction buffer.

### Liposomes preparation

Liposomes were prepared by rehydrating a lipid film of DOPC: DOPG: DOPE (3 mg : 0.75 mg : 1.25 mg) after 4 h of desiccation by adding 1 ml of buffer (20 mM HEPES, pH 7.5, 50 mM NaCl, 15% glycerol) with 6 mM BCECF and HEPES at pH 8 for permeability tests. The lipid suspension underwent 10 freeze-thaw cycles in liquid nitrogen and was extruded through a 100 nm polycarbonate membrane. Protein reconstitution was performed by destabilizing the liposomes with 5.5 mM Triton X-100 for 15 min. The protein was added at a final concentration of 15 µg/ml and the mixture was incubated at 4°C for 15 min. Detergent removal was achieved for 5 h at 4°C with 25 mg of Biobeads prehydrated with agitation, repeated 4 times, followed by ultracentrifugation at 16,000 × g for 20 min at 4°C to harvest proteoliposomes in the pellet. Liposomes containing BCECF were purified using a PD-10 column (GE Healthcare). Eluates (500 µL) were collected and analyzed by fluorescence measurement (λ_ex_ 485nm, λ_em_ 530nm) and by western blot using peroxidase coupled anti-His tag antibody (Sigma-Aldrich) diluted 1:2000 to identify fractions containing proteoliposomes with internalized BCECF.

### C_55_PP phosphatase assay in liposomes

Reaction mixture contained 20 µM C_15_-PP, 20 µM [^14^C]C_5_-PP, and 180 µM C_5_-PP in 20 mM HEPES, pH 7.5, 50 mM NaCl, and 15% glycerol. For each kinetics assay, 5 µL of the reaction mixture was diluted with 5 µL of proteoliposomes supplemented with UppS at a final concentration of 0.2 mg/mL. The mixture was incubated at 37°C for 10 min and the reaction was then stopped by plunging the tube in liquid nitrogen. Substrates and products were separated and quantified by thin layer chromatography as previously described.

### Proteoliposome permeabilisation assay

BCECF-containing liposomes were diluted in buffer 20 mM HEPES, pH 6, 50 mM NaCl, 15% glycerol in a black Corning Flat Bottom Polystyrol 96 well plate. The fluorescence was then monitored for 1h with an excitation wavelength of 485 nm and an emission wavelength of 530 nm before and after addition of 5 µM EntC. Fluorescence measurements are normalised based on a value of 1, which was attributed to the last fluorescence value before EntC addition.

### Crosslinking experiments

Crosslinking experiments was performed using the amine-reactive, membrane-permeable, and non-cleavable crosslinker disuccinimidyl suberate (DSS) purchased by Sigma. EntC peptides and BacA protein were mixed at equimolar concentrations (17 µM) in buffer D in the presence of an excess of DSS crosslinker (1000-fold or 100-fold) in a final reaction volume of 20 µL. The reaction was incubated for 30 min at 4°C and any unreacted DSS was quenched with 50 mM Tris-HCl. After separation on 12% acrylamide gel, proteins were electrotransferred onto polyvinylidene difluoride (PVDF) membrane and BacA was detected using peroxidase-coupled anti-His tag antibody by chemiluminescence detection using the ChemiDoc MP system (BioRad).

### Functional complementation tests

The thermosensitive *E. coli* strain BWTetra-Ts*bacA* (El Ghachi *et al*, 2005) was transformed by heat shock with plasmids pET2130 carrying the *bacA_ef_* gene or its mutated variants and the functional complementation test was performed as previously described (Manat *et al*, 2015).

### Flow-Induced dispersion analysis (FIDA)

The experiments were performed on a FIDA 1 instrument equipped with a 480 nm LED induced fluorescence detector and a FIDA permanently coated capillary (FIDA Biosystems ApS, Copenhagen Denmark). The sample tray and the capillary chamber were temperature-controlled at 25°C. The EntC1 peptide was first labelled using the FL 480 Fida1 Protein Labelling Kit according to the manufacturer’s instructions. Briefly, the EntC1 peptide was incubated overnight at 4°C with 5-fold molar excess of FAM NHS ester reactive dye. The free dye was removed using two successive size exclusion chromatography and the purification efficiency was checked using the FIDA1 instrument. The degree of labelling was determined by measuring the absorbance at both 280 nm et 495 nm and was found to be 6%. All protein samples were prepared in the assay buffer 20 mM Tris-HCl pH7.4, 200 mM NaCl and 0.1% DDM. All the binding experiments were carried out using the premix method to maintain binding equilibrium during the run. Following the FIDA experimental procedure, the capillary was first rinsed and equilibrated with the assay buffer at 3500 mbar for 2 min. Next the unlabelled protein was filled at 3500 mbar for 20 sec, after which the premix sample (Fluorescein-EntC1 preincubated with the unlabelled protein) was injected at 50 mbar for 10 sec. Finally, the indicator was flowed towards the fluorescence detector with the analyte sample at 400 mbar for 180 sec. For binding curves, the FL-EntC1 indicator was fixed at 100 nM and titrated with 0-4000 nM BacA*_ef_* analyte in the absence or presence of 200 nM EntC2 peptide. The Taylorgrams were processed using the FIDA data analysis software (version 2.32) to determine the apparent hydrodynamics radius (Rh) and fit binding curves to the best binding model.

### Isothermal titration microcalorimetry measurements (ITC)

Isothermal titration microcalorimetry experiments were performed with an ITC200 isothermal titration calorimeter from MicroCal (Malvern Panalytical, Malvern, UK). The experiments were carried out at 20 °C. The protein concentration in the microcalorimeter cell (0.2 mL) was 33 μM. Nineteen 2- μL injections of peptides at 400 μM were performed at intervals of 240 s while stirring at 500 rpm. The experimental data were fitted to theoretical titration curves with the software supplied by MicroCal (ORIGIN®, Northampton, MA, USA). This software uses the relationship between the heat generated by each injection and ΔH (enthalpy change in kcal/mol), *K*_a_ (the association binding constant in M^−1^), n (the number of binding sites), total protein concentration and free and total ligand concentrations.

### Differential scanning calorimetry (DSC)

Thermal stability of BacA protein in different mixtures of peptides was performed by DSC on a MicroCal model VP-DSC (Malvern Panalytical, Malvern, UK). BacA (33 µM) was mixed with different mixtures of peptides at twice BacA concentration (i.e. 66 µM). Each measurement was preceded by a baseline scan with the standard buffer. Scans were performed at 1 K/min (60°C/h) temperature ramp between 30 °C and 100 °C. The heat capacity of the buffer was subtracted from that of the protein sample before analysis. The thermodynamic parameters were determined by fitting the data to the following equation:

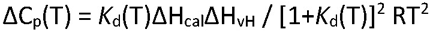

where *K*_d_ is the equilibrium constant for a two-state process, ΔH_vH_ the enthalpy calculated based on a two-state process and ΔH_cal_ the measured enthalpy.

### Alphafold2 modelling

Alphafold2 v2.3 modelling of the predicted 3D protein structures was performed via the AlphaFold@I2BC server accessible from the I2BC (Institute for Integrative Biology of the Cell). Multiple Sequence Alignments (MSAs) relies on MMseqs2 (Mirdita *et al*, 2019) while pairwise representation of amino acids is generated using colabfold v1.5.2 (Mirdita *et al*, 2022). Multimeric protein structures were predicted using AlphaFold-Multimer (Evans *et al*, 2021). The structural model of the tripartite complex BacA/EntC1/EntC2 is available in ModelArchive at https://modelarchive.org/doi/10.5452/ma-2xesy (Tauriello *et al*, 2025).

## Author contribution

**Victor FOLCHER**: Resources; Data curation; Formal analysis; Validation; Investigation; Methodology; Writing—original draft; Writing-review and editing. **Rodolphe AUGER**: Resources; Data curation; Formal analysis; Validation; Investigation; Methodology; Writing-review. **Clara LOUCHE**: Formal analysis; Validation; Investigation; Methodology. **Pascale SERROR**: Conceptualization; Resources; Supervision; Validation; Methodology; Writing—review and editing. **Thierry TOUZE**: Conceptualization; Data curation; Formal analysis; Supervision; Funding acquisition; Validation; Investigation; Methodology; Project administration; Writing-review and editing.

## Acknowledgements

We thank the BIOI2 platform from I2BC for making the ColabFold pipeline easily accessible. The present work has benefited from the facility of macromolecular interactions (PIM platform) of I2BC, supported by French Infrastructure for Integrated Structural Biology (FRISBI). We thank Magali Aumont-Nicaise from the platform PIM for assistance with the DSC and ITC analyses. This work has benefited from the facilities and expertise of the I2BC proteomic platform SICaPS, supported by IBiSA, Ile de France Region, Plan Cancer, CNRS and Paris-Sud University, for nano-LC-MSMS analyses. We thank FidaBio for providing the FIDA device. We thank Françoise Le Bohec for the construction of the *E. faecalis* strain VE18740 (Δ*bacA_ef_*) and plasmid pVE14417.

## Data Availability

The structural model is available in ModelArchive

at https://modelarchive.org/doi/10.5452/ma-2xesy.

## Figures Expanded View

**Fig. EV1.**
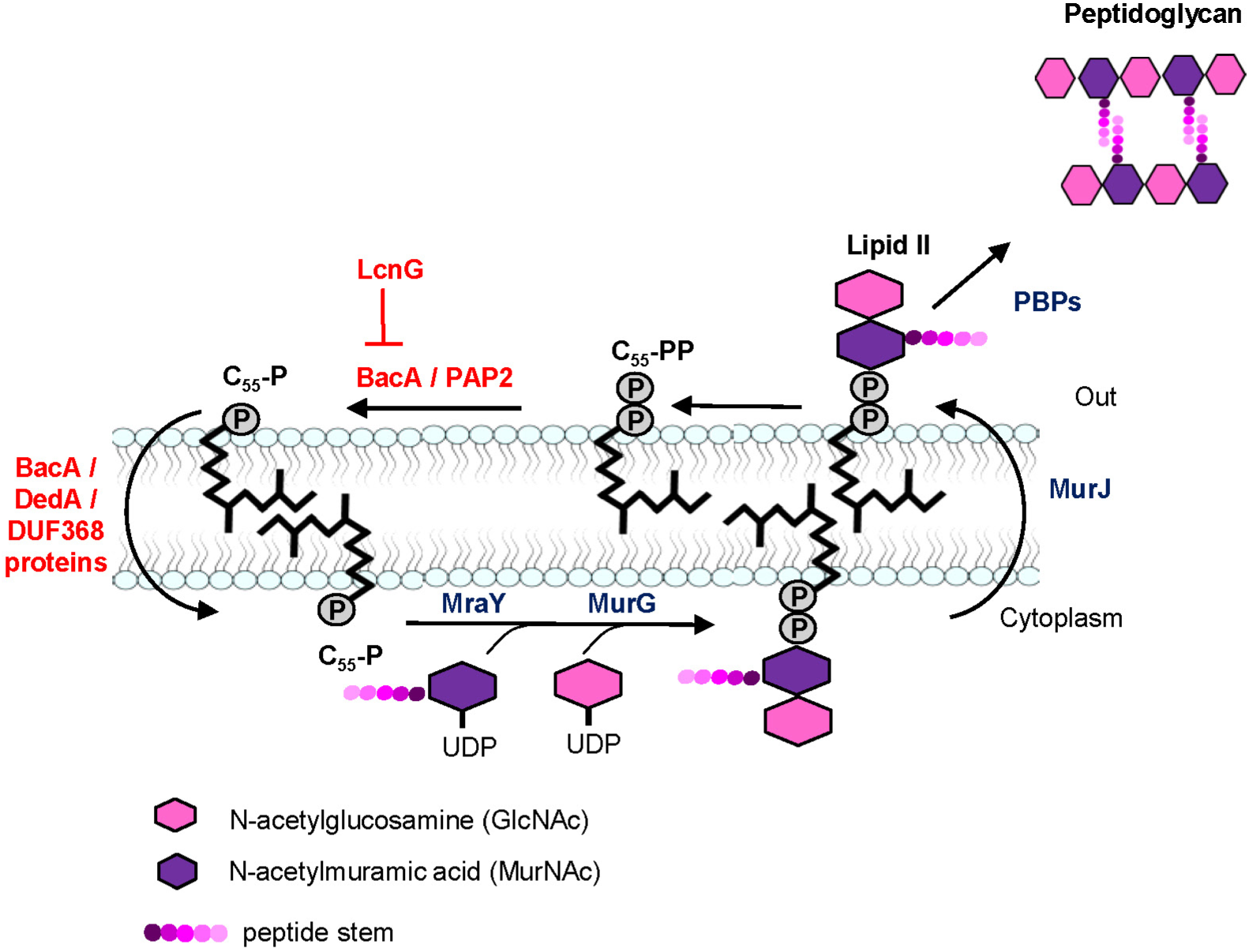
Schematic representation of lipid II cycle. Peptidoglycan synthesis starts in the cytoplasm with the successive transfer of phospho-MurNAc-pentapeptide and GlcNAc moieties from cytosoluble nucleotide precursors onto the lipid carrier undecaprenyl-phosphate (C_55_-P), yielding lipid II. The membrane intermediate is then translocated across the plasma membrane by the flippase MurJ. The peptidoglycan subunit is subsequently transferred from lipid II to the nascent peptidoglycan strand releasing the lipid carrier in its pyrophosphorylated form, C_55_-PP. the C_55_-PP is then recycled by a dephosphorylation reaction catalyzed by either BacA or a member of the type 2 phosphatidic acid phosphatase (PAP2) superfamily and a flip back to the inner side, hypothetically by the action of BacA, DedA proteins or DUF-368-containing proteins. Overall, these successive steps constitute the so-called lipid II cycle.

**Fig. EV2.**
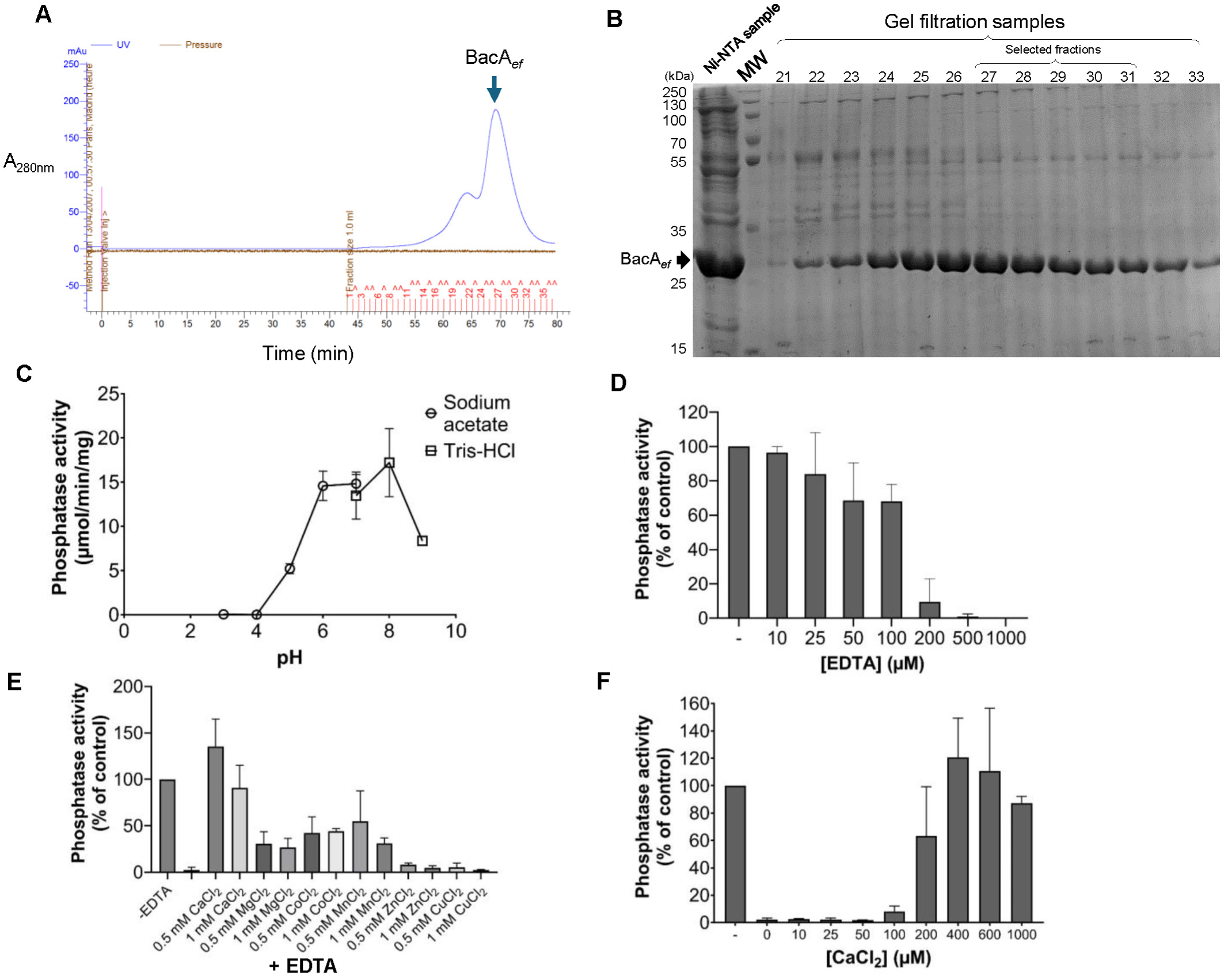
Purification and biochemical characterization of BacA*_ef_*. (**A**) Gel filtration chromatogram on a Superdex 200 resin of BacA*_ef_* sample protein obtained from affinity purification on Ni^2+^-NTA-agarose. Elution was performed at a flow rate of 1 mL/min, coupled with UV spectrophotometry at 280 nm (A_280nm_) measured at the column outlet (blue line). The pressure during the size-exclusion chromatography is represented by the brown line. The collected samples are indicated by red ticks, along with their corresponding numbers. (**B**) SDS-PAGE analysis of 10 µL protein samples obtained from affinity purification and gel filtration chromatography. The samples were prepared in 1x Laemmli buffer and separated on a 12% acrylamide gel. Proteins were revealed using Coomassie Blue staining. The molecular weight (MW) marker is indicated on the left and BacA*_ef_* is indicated by a black arrow. The samples that were selected for further studies were pooled and concentrated up to 1 mg/mL, using a VivaSpin filter, before their storage at −20°C. (**C**) BacA*_ef_* C_55_-PP phosphatase activity measured at various pH. Activity measurements were performed in sodium acetate buffer for pH ranging from 3 to 7 (round dots), and Tris-HCl buffer for pH ranging from 7 to 9 (square dots). (**D**) Dose-dependent effect of EDTA on BacA*_ef_* C_55_-PP phosphatase activity. The activity was measured at pH 7.5 in Tris-HCl buffer in the presence of 0.1% DDM. (**E**) Restoration of BacA*_ef_* activity by different divalent cations following treatment with 500 µM EDTA. (**F**) Dose-dependent effect of CaCl₂ on the activity of EDTA-treated BacA*_ef_*. Activity was measured at pH 7.5 in Tris-HCl buffer with 0.1% DDM.

**Fig. EV3.**
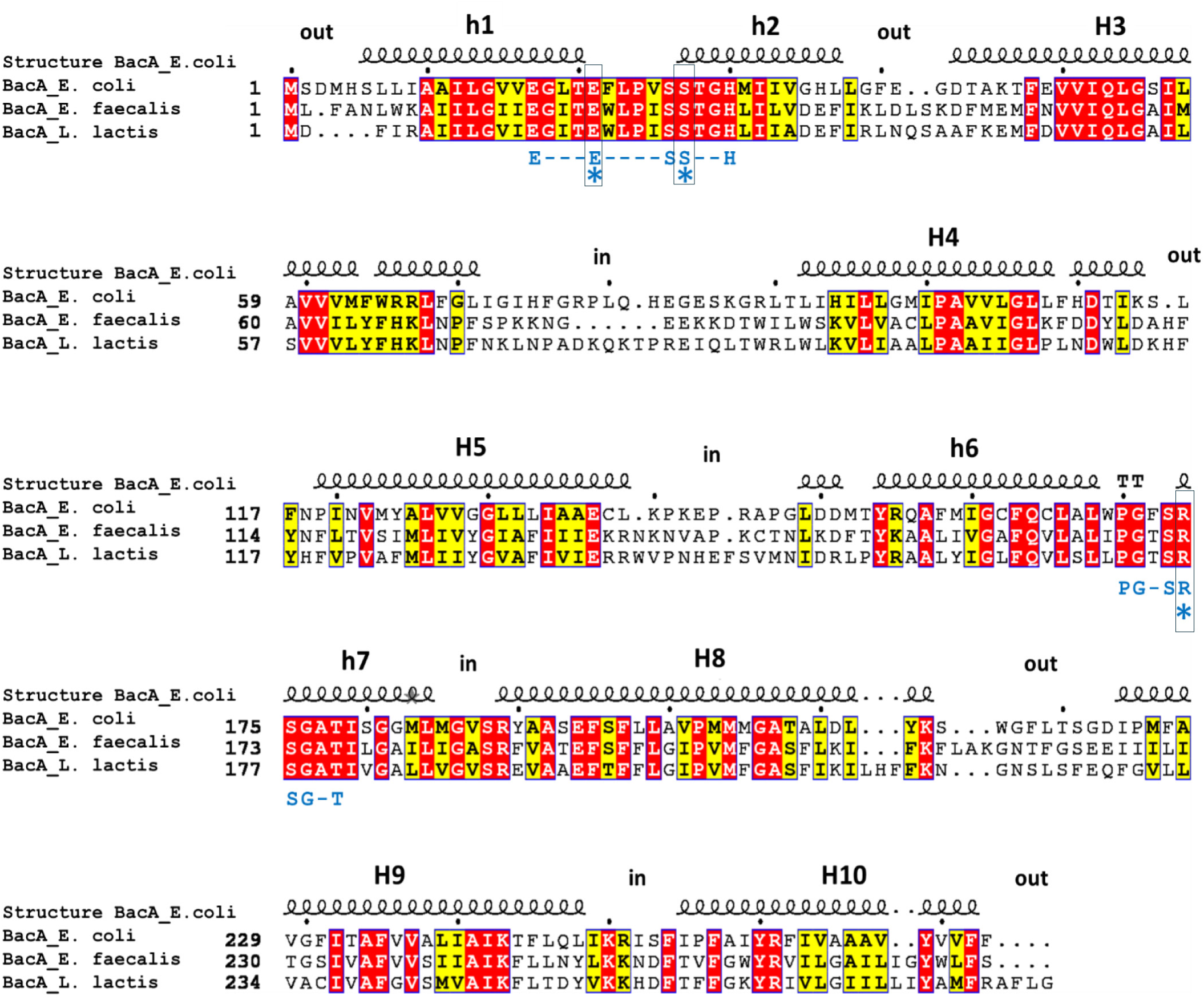
Sequence alignment of BacA homologues from *E. coli*, *E. faecalis* and *L. lactis*. Identical and similar residues in BacA homologues are marked with red and yellow fills, respectively. The BacA consensus sequence is indicated in blue below the sequence alignment and active site residues are indicated by stars. Secondary structural elements and the position of membrane-embedded helices (H1 to H10), according to BacA*_ec_* crystallographic structure, are indicated above the sequence.

**Fig. EV4.**
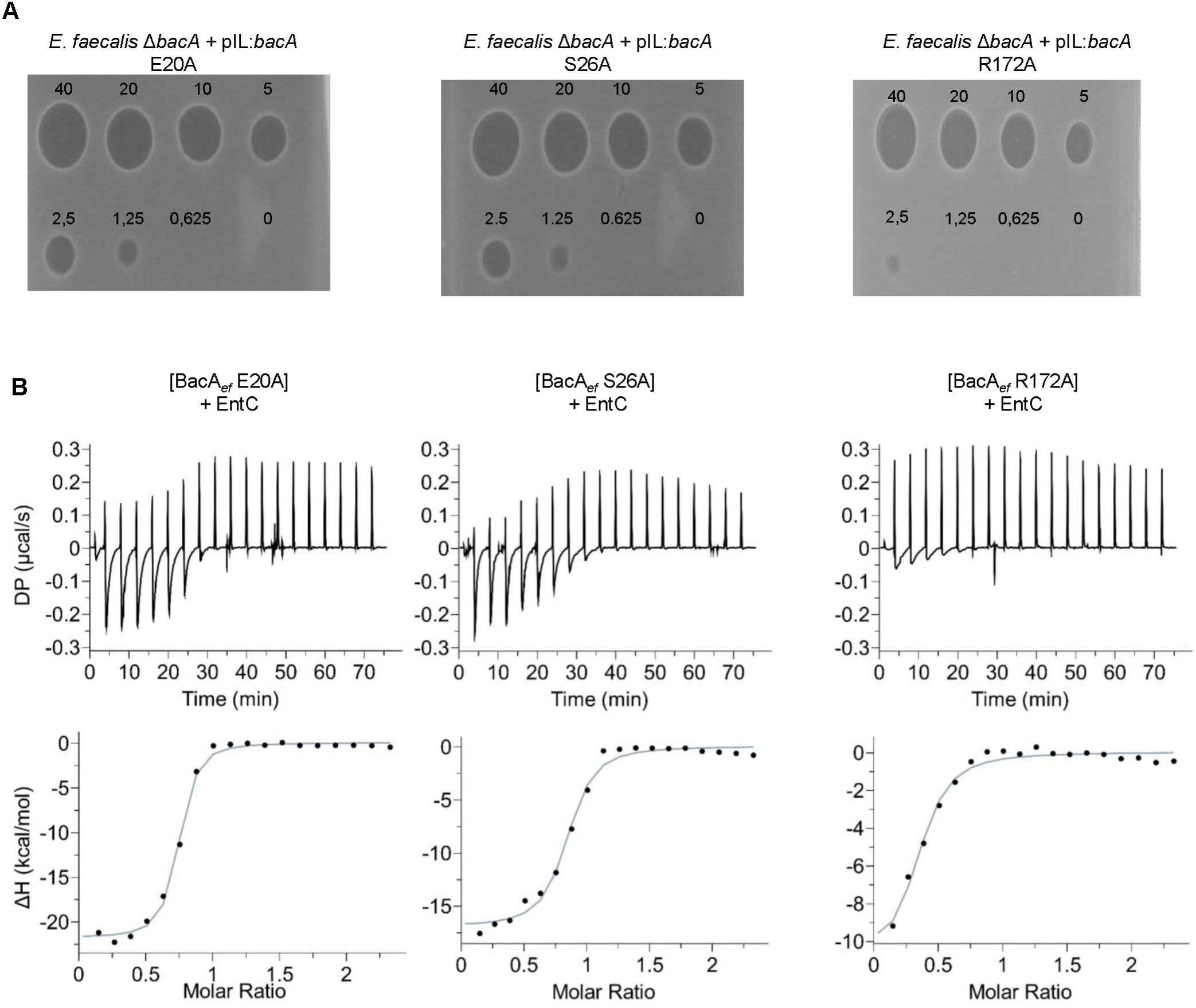
Effect of substitution of BacA*_ef_* catalytic residues on the activity of EntC. (**A**) Spot-on-lawn assays of EntC on a lawn of *E. faecalis* Δ*bacA* cells expressing catalytically inactive variants of BacA*_ef_*. (**B**) ITC scans of BacA*_ef_* variants titration with EntC. Upper panels, raw ITC thermograms from the titration of BacA*_ef_* variants (32 µM) with EntC bacteriocin (400 µM). Lower panels, heat profile from peak integration of ITC thermograms to determine the thermodynamic parameters of the interaction. The data shown are representative of experiments performed in triplicate.

**Fig. EV5.**
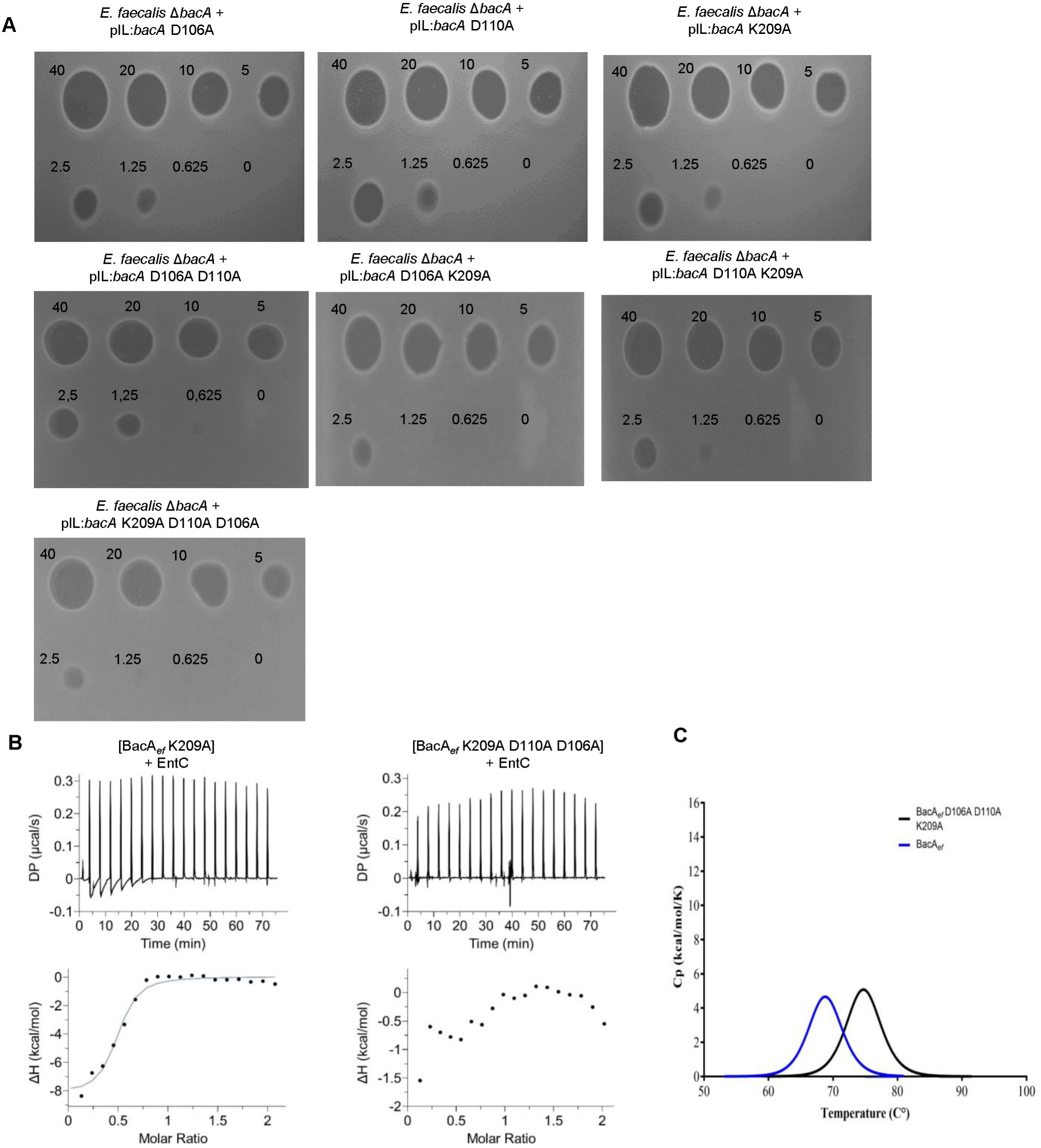
Effect of substitution of specific BacA*_ef_* residues on the activity of EntC. **(A)** Spot-on-lawn assays of EntC on a lawn of *E. faecalis* Δ*bacA* cells expressing various BacA*_ef_* variants. **(B)** ITC scans of BacA*_ef_* variants titration with EntC. Upper panels, raw ITC thermograms from the titration of BacA*_ef_* variants (32 µM) with EntC bacteriocin (400 µM); Lower panels, heat profile from peak integration of the ITC thermograms to determine the thermodynamic parameters of the interaction. The data shown are representative of experiments performed in triplicate. **(C)** Stability measurement of BacA*_ef_* and BacA*_ef_* D106A D110A K209A variant. Representative DSC thermograms of protein (32 µM).

