## supplemental data for "Mechanistic insights into Enterocin C targeting the undecaprenyl phosphate recycling protein BacA"

### Slide 1
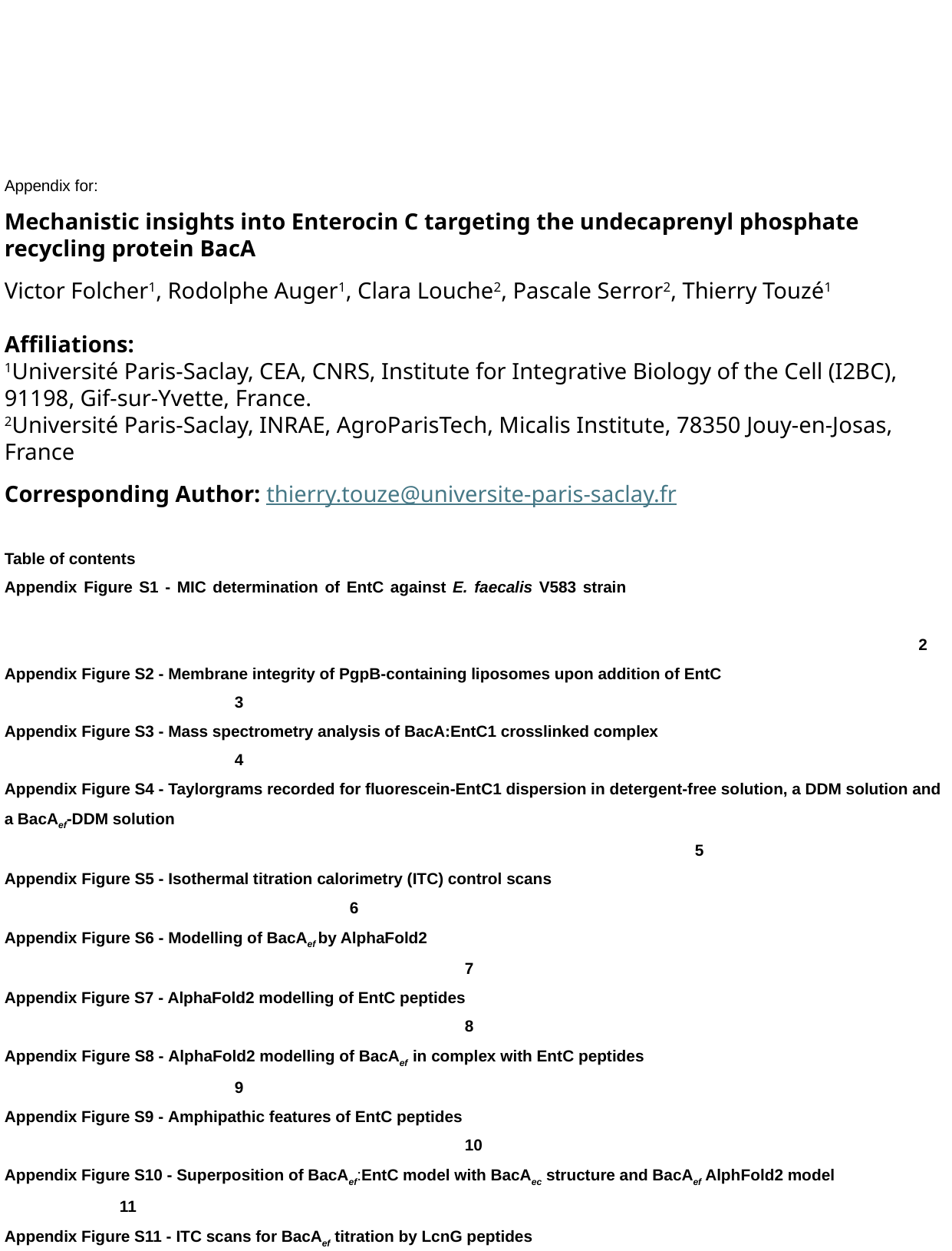

Appendix for:
Mechanistic insights into Enterocin C targeting the undecaprenyl phosphate recycling protein BacA
Victor Folcher^1^, Rodolphe Auger^1^, Clara Louche^2^, Pascale Serror^2^, Thierry Touzé^1^
Affiliations:
^1^Université Paris-Saclay, CEA, CNRS, Institute for Integrative Biology of the Cell (I2BC), 91198, Gif-sur-Yvette, France.
^2^Université Paris-Saclay, INRAE, AgroParisTech, Micalis Institute, 78350 Jouy‑en‑Josas, France
Table of contents
Appendix Figure S1 - MIC determination of EntC against E. faecalis V583 strain 		2
Appendix Figure S2 - Membrane integrity of PgpB-containing liposomes upon addition of EntC 				3
Appendix Figure S3 - Mass spectrometry analysis of BacA:EntC1 crosslinked complex 					4
Appendix Figure S4 - Taylorgrams recorded for fluorescein-EntC1 dispersion in detergent-free solution, a DDM solution and a BacA_ef_-DDM solution 													5
Appendix Figure S5 - Isothermal titration calorimetry (ITC) control scans 							6
Appendix Figure S6 - Modelling of BacA_ef_ by AlphaFold2 									7
Appendix Figure S7 - AlphaFold2 modelling of EntC peptides								8
Appendix Figure S8 - AlphaFold2 modelling of BacA_ef_ in complex with EntC peptides 					9
Appendix Figure S9 - Amphipathic features of EntC peptides 								10
Appendix Figure S10 - Superposition of BacA_ef_:EntC model with BacA_ec_ structure and BacA_ef_ AlphFold2 model		11
Appendix Figure S11 - ITC scans for BacA_ef_ titration by LcnG peptides 							12
Appendix Figure S12 - AlphaFold2 model of BacA_ef_ with LcnG peptides. 							13
Appendix Figure S13 - Details on Interfacial regions within AlphaFold2 models of BacA_ef_ in complex with LcnG peptides	14
Appendix Figure S14 - ITC scans for BacA_ef_ titration by EntC/LcnG chimera 							15
Appendix Table S1 - Found Interfaces in BacA_ef_:EntC tripartite complex according to PISA 					16
Appendix Table S2 - in vivo and in vitro functionality of BacA_ef_ variants							17
Appendix Table S3 - Oligonucleotides used in this studya									18

### Slide 2
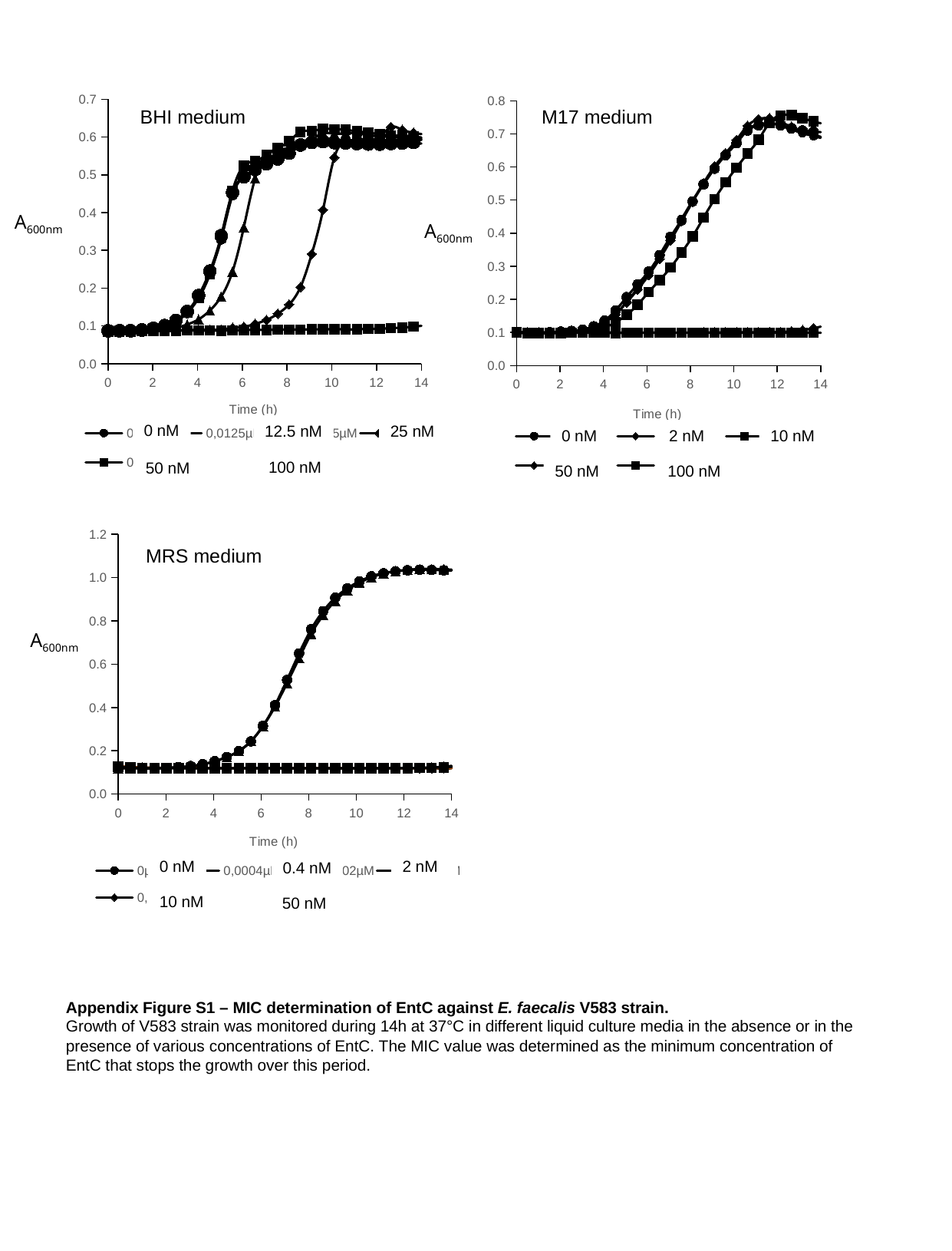

#### Chart
| Category | 0µM | 0µM | 0,002µM | 0,01µM | 0,05µM | 0,1µM |
|---|---|---|---|---|---|---|M17 medium
0 nM
2 nM
10 nM
50 nM
100 nM
#### Chart
| Category | 0 | 0 | 0,0125µM | 0,025µM | 0,05µM | 0,1µM |
|---|---|---|---|---|---|---|BHI medium
A_600nm_
A_600nm_
0 nM
12.5 nM
25 nM
100 nM
50 nM
#### Chart
| Category | 0µM | 0,0004µM | 0,002µM | 0,01µM | 0,05µM | 0,05µM |
|---|---|---|---|---|---|---|MRS medium
A_600nm_
0 nM
2 nM
0.4 nM
10 nM
50 nM
Appendix Figure S1 – MIC determination of EntC against E. faecalis V583 strain.
Growth of V583 strain was monitored during 14h at 37°C in different liquid culture media in the absence or in the presence of various concentrations of EntC. The MIC value was determined as the minimum concentration of EntC that stops the growth over this period.

### Slide 3
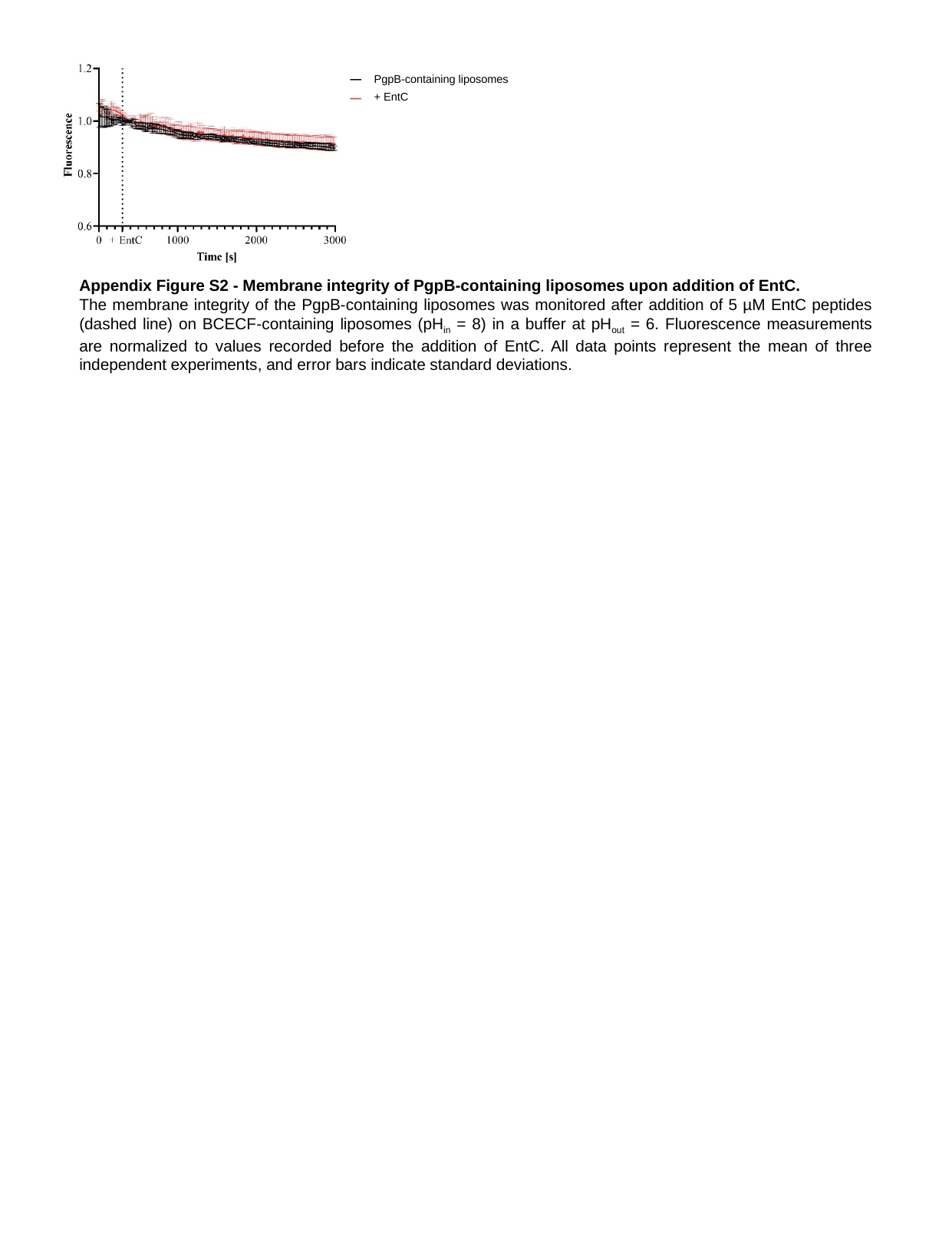

PgpB-containing liposomes
+ EntC
Appendix Figure S2 - Membrane integrity of PgpB-containing liposomes upon addition of EntC.
The membrane integrity of the PgpB-containing liposomes was monitored after addition of 5 µM EntC peptides (dashed line) on BCECF-containing liposomes (pH_in_ = 8) in a buffer at pH_out_ = 6. Fluorescence measurements are normalized to values recorded before the addition of EntC. All data points represent the mean of three independent experiments, and error bars indicate standard deviations.

### Slide 4
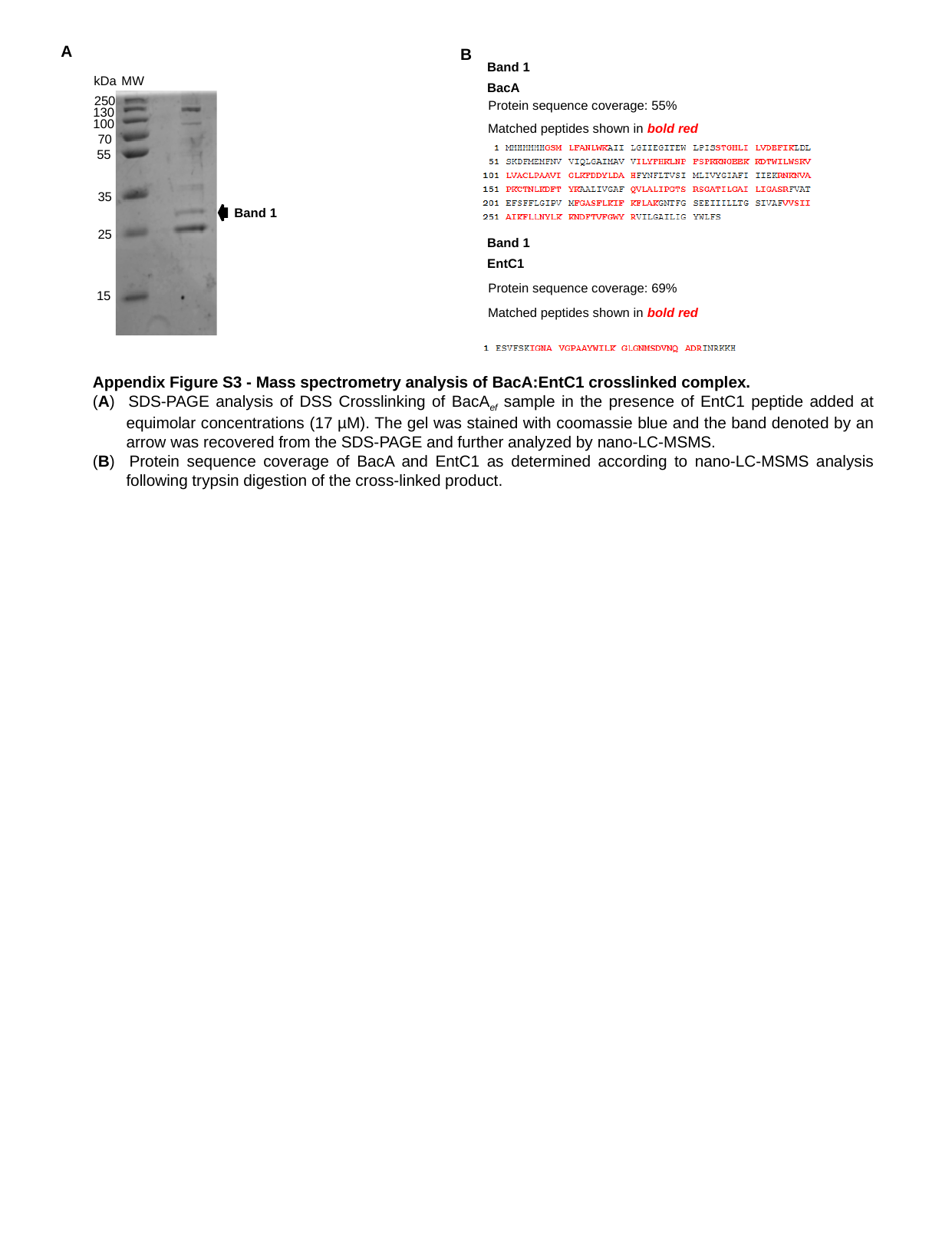

A
B
Band 1
kDa
MW
250
130
100
70
55
35
25
15
BacA
Protein sequence coverage: 55%
Matched peptides shown in bold red
Band 1
Band 1
EntC1
Protein sequence coverage: 69%
Matched peptides shown in bold red
Appendix Figure S3 - Mass spectrometry analysis of BacA:EntC1 crosslinked complex.
(A) 	SDS-PAGE analysis of DSS Crosslinking of BacA_ef_ sample in the presence of EntC1 peptide added at equimolar concentrations (17 µM). The gel was stained with coomassie blue and the band denoted by an arrow was recovered from the SDS-PAGE and further analyzed by nano-LC-MSMS.
(B) 	Protein sequence coverage of BacA and EntC1 as determined according to nano-LC-MSMS analysis following trypsin digestion of the cross-linked product.

### Slide 5
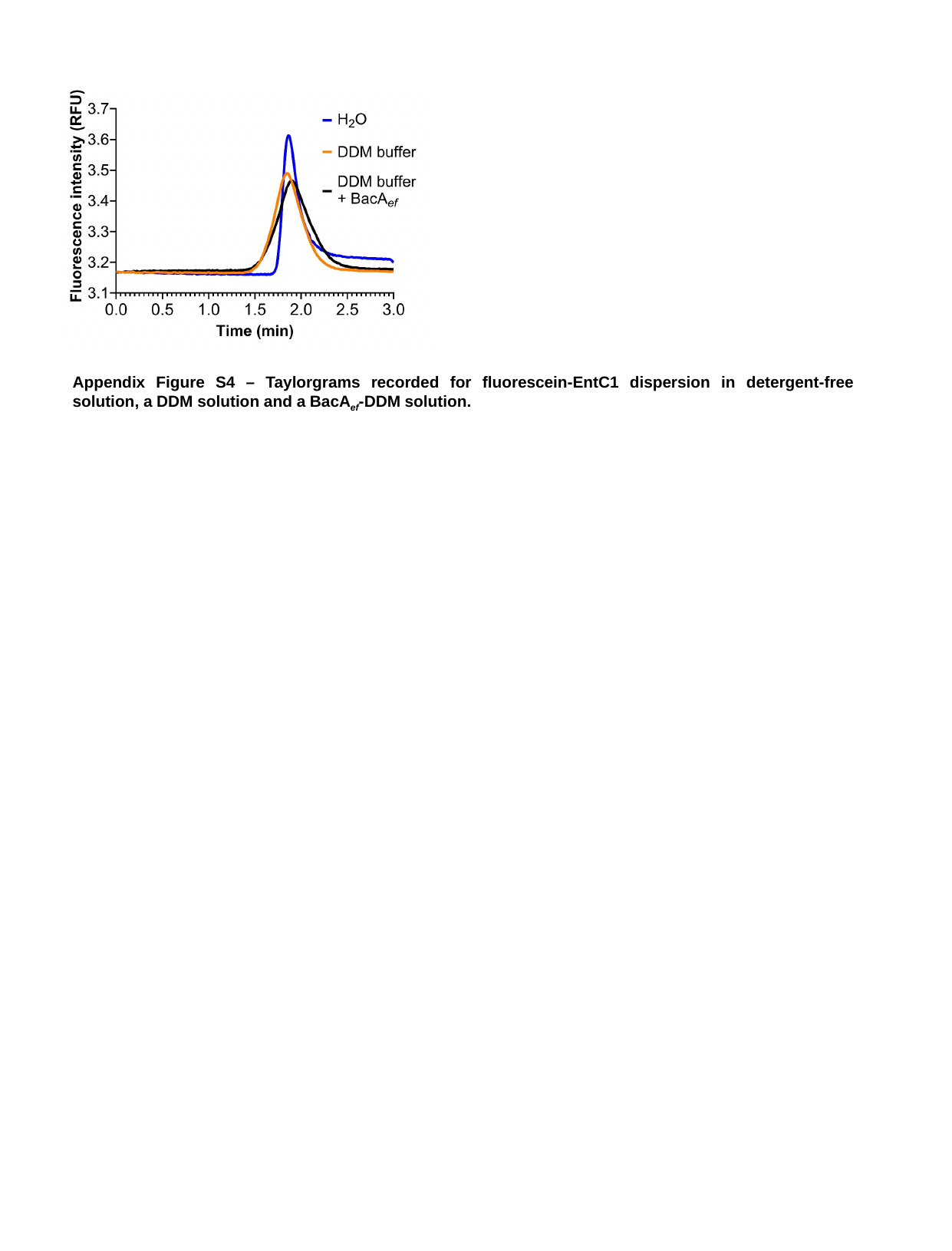

Appendix Figure S4 – Taylorgrams recorded for fluorescein-EntC1 dispersion in detergent-free solution, a DDM solution and a BacA_ef_-DDM solution.

### Slide 6
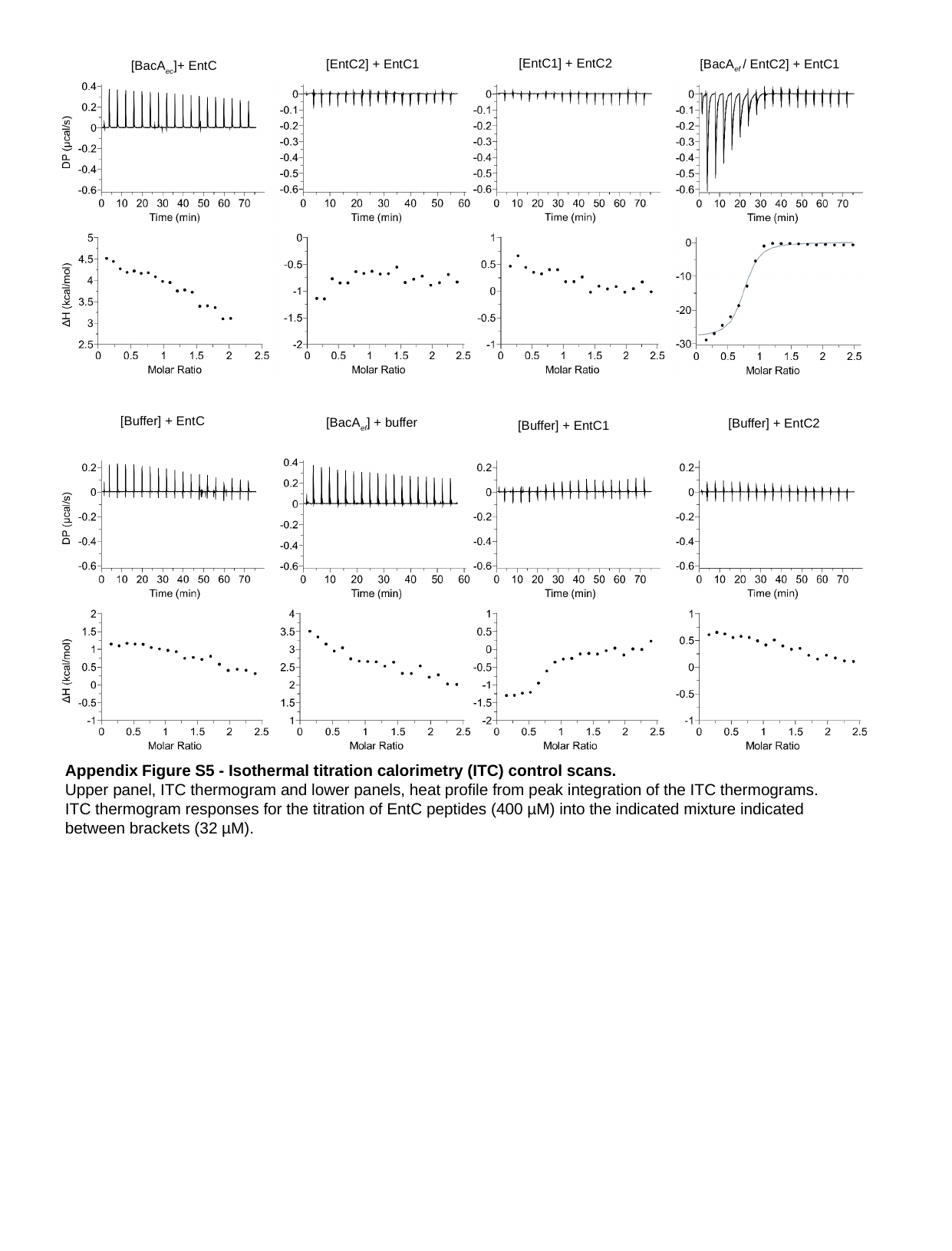

[EntC1] + EntC2
[BacA_ef_ / EntC2] + EntC1
[EntC2] + EntC1
[BacA_ec_]+ EntC
No binding
[Buffer] + EntC
[BacA_ef_] + buffer
[Buffer] + EntC2
[Buffer] + EntC1
Appendix Figure S5 - Isothermal titration calorimetry (ITC) control scans.
Upper panel, ITC thermogram and lower panels, heat profile from peak integration of the ITC thermograms.
ITC thermogram responses for the titration of EntC peptides (400 µM) into the indicated mixture indicated between brackets (32 µM).

### Slide 7
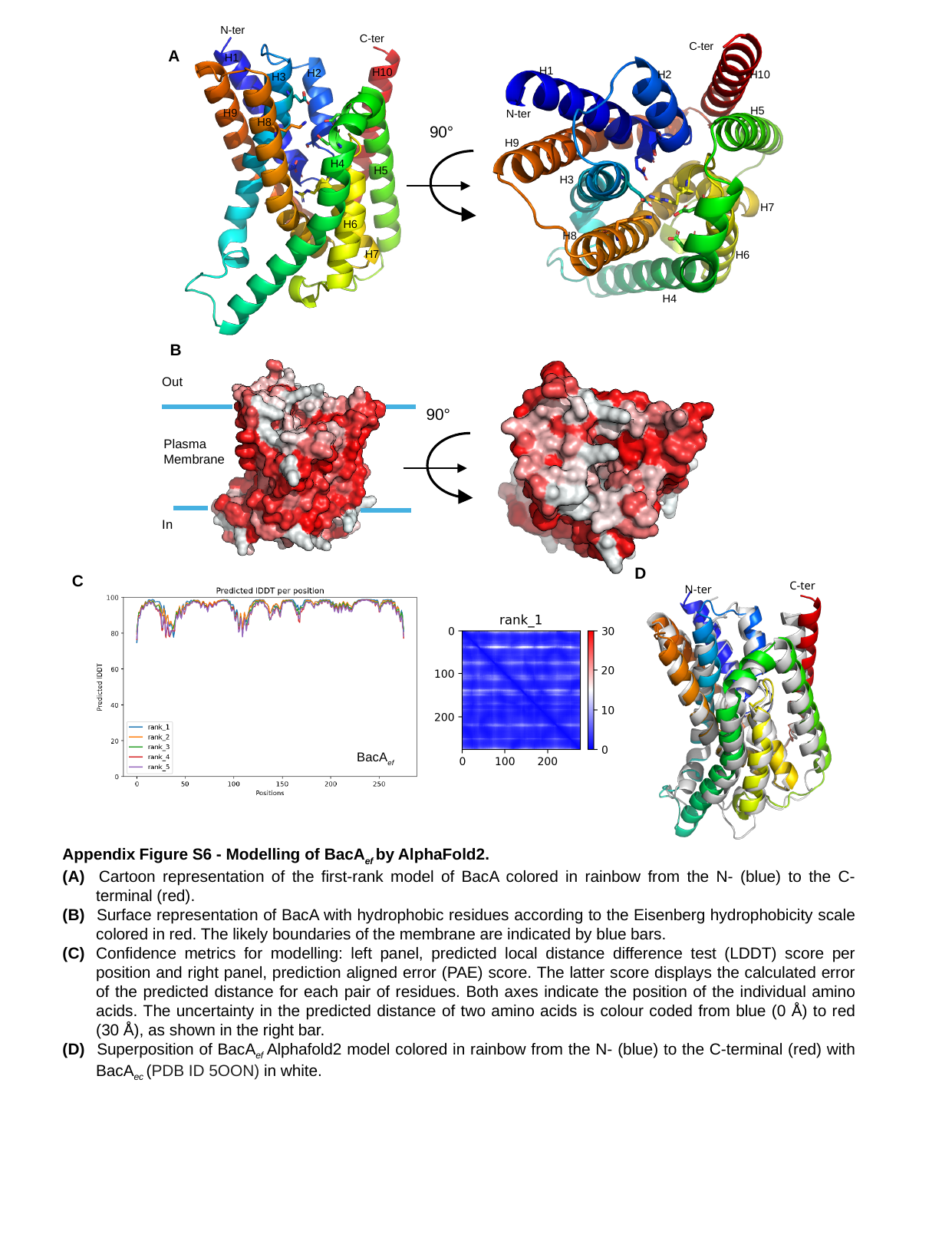

N-ter
C-ter
H1
H10
H2
H3
H9
H8
H4
H5
H6
H7
C-ter
H1
H2
H10
H5
N-ter
H9
H3
H7
H8
H6
H4
A
90°
B
Out
90°
Plasma
Membrane
In
D
C
C-ter
N-ter
BacA_ef_
Appendix Figure S6 - Modelling of BacA_ef_ by AlphaFold2.
(A) 	Cartoon representation of the first-rank model of BacA colored in rainbow from the N- (blue) to the C-terminal (red).
(B) 	Surface representation of BacA with hydrophobic residues according to the Eisenberg hydrophobicity scale colored in red. The likely boundaries of the membrane are indicated by blue bars.
(C)	Confidence metrics for modelling: left panel, predicted local distance difference test (LDDT) score per position and right panel, prediction aligned error (PAE) score. The latter score displays the calculated error of the predicted distance for each pair of residues. Both axes indicate the position of the individual amino acids. The uncertainty in the predicted distance of two amino acids is colour coded from blue (0 Å) to red (30 Å), as shown in the right bar.
(D) 	Superposition of BacA_ef_ Alphafold2 model colored in rainbow from the N- (blue) to the C-terminal (red) with BacA_ec_ (PDB ID 5OON) in white.

### Slide 8
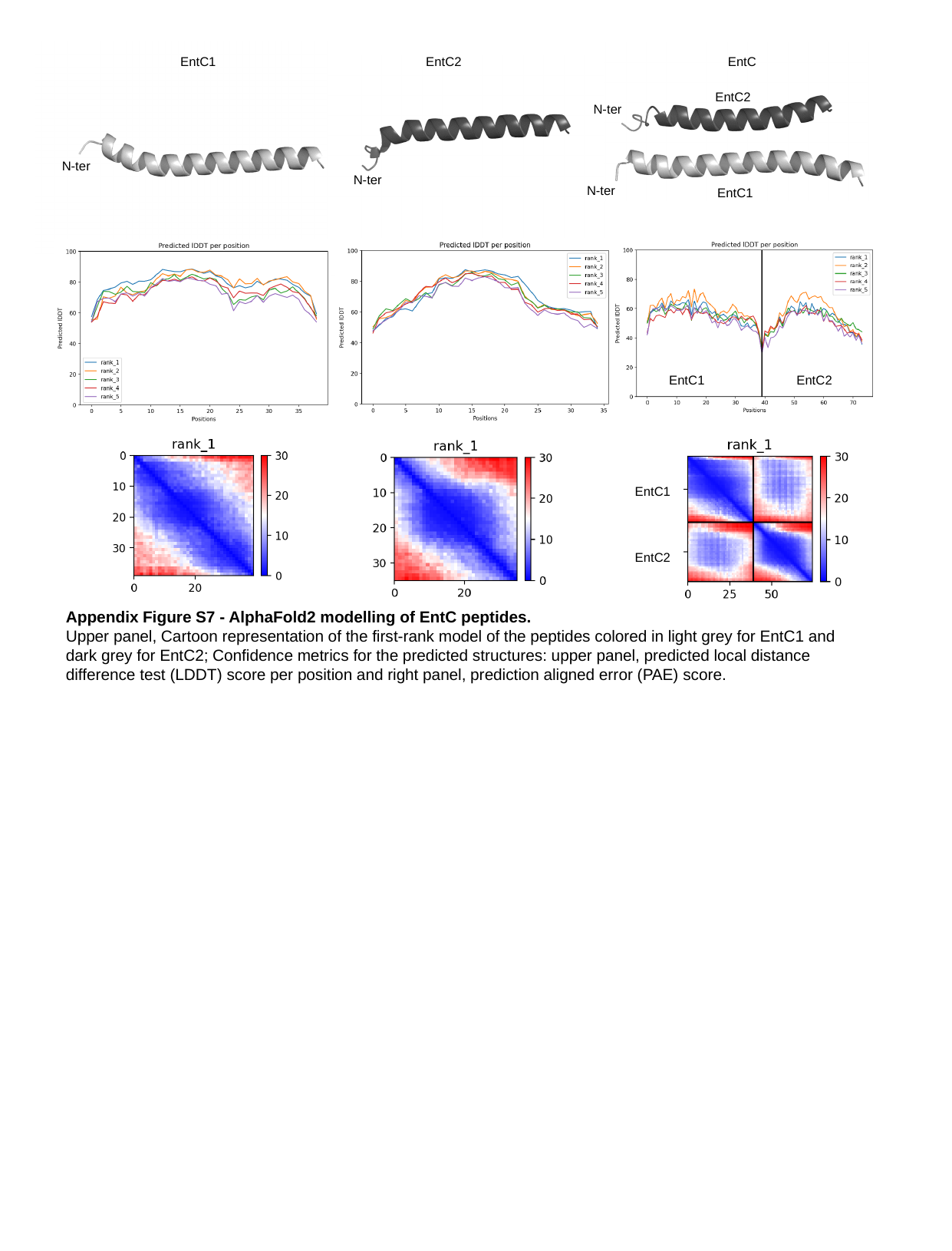

EntC1
EntC2
EntC
EntC2
N-ter
N-ter
N-ter
N-ter
EntC1
EntC1
EntC2
EntC1
EntC2
Appendix Figure S7 - AlphaFold2 modelling of EntC peptides.
Upper panel, Cartoon representation of the first-rank model of the peptides colored in light grey for EntC1 and dark grey for EntC2; Confidence metrics for the predicted structures: upper panel, predicted local distance difference test (LDDT) score per position and right panel, prediction aligned error (PAE) score.

### Slide 9
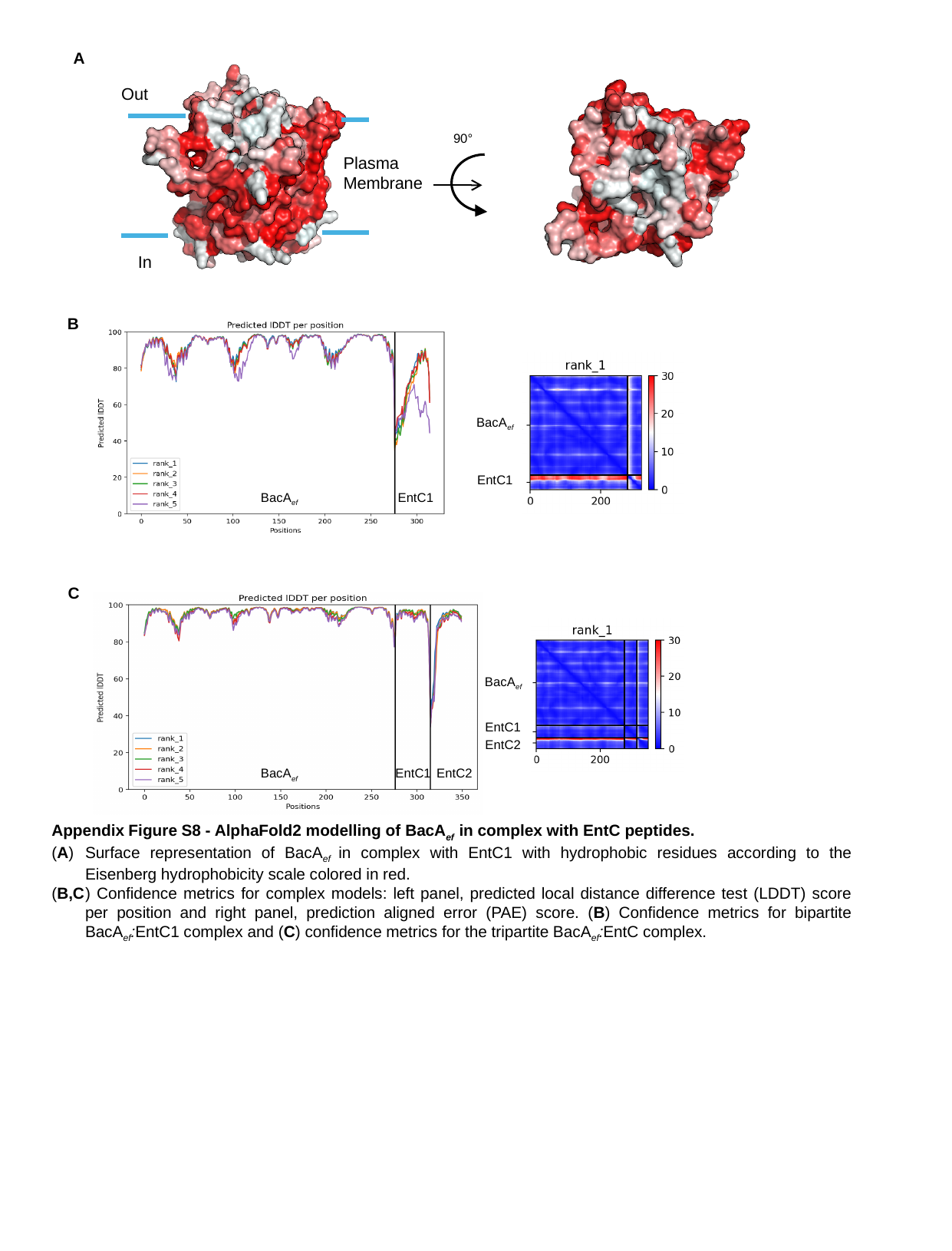

A
Out
Plasma
Membrane
In
90°
B
BacA_ef_
EntC1
BacA_ef_
EntC1
C
BacA_ef_
EntC1
EntC2
BacA_ef_
EntC1
EntC2
Appendix Figure S8 - AlphaFold2 modelling of BacA_ef_ in complex with EntC peptides.
(A)	Surface representation of BacA_ef_ in complex with EntC1 with hydrophobic residues according to the Eisenberg hydrophobicity scale colored in red.
(B,C	) Confidence metrics for complex models: left panel, predicted local distance difference test (LDDT) score per position and right panel, prediction aligned error (PAE) score. (B) Confidence metrics for bipartite BacA_ef_:EntC1 complex and (C) confidence metrics for the tripartite BacA_ef_:EntC complex.

### Slide 10
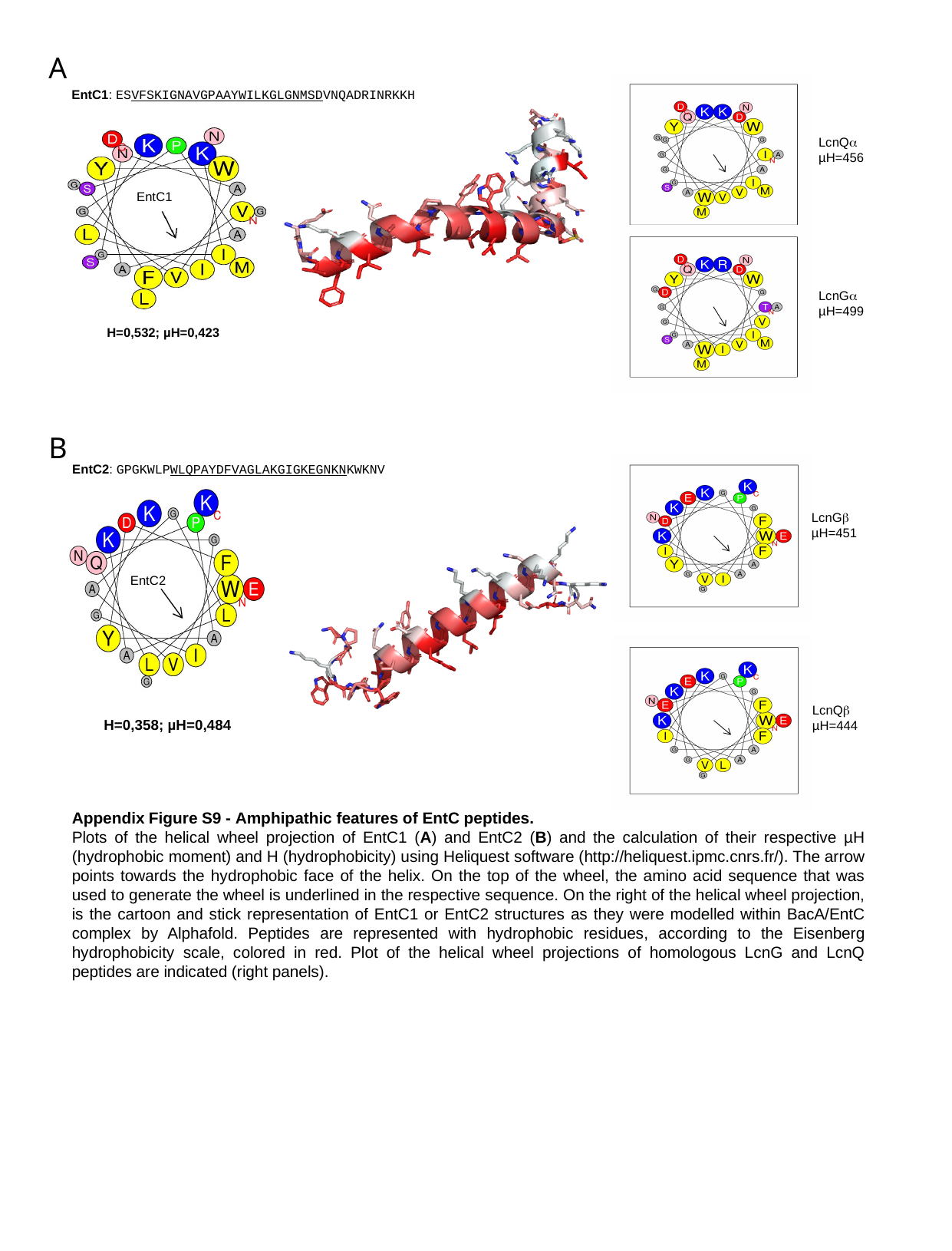

A
B
EntC1: ESVFSKIGNAVGPAAYWILKGLGNMSDVNQADRINRKKH
LcnQα
µH=456
EntC1
LcnGα
µH=499
H=0,532; µH=0,423
EntC2: GPGKWLPWLQPAYDFVAGLAKGIGKEGNKNKWKNV
LcnGβ
µH=451
EntC2
LcnQβ
µH=444
H=0,358; µH=0,484
Appendix Figure S9 - Amphipathic features of EntC peptides.
Plots of the helical wheel projection of EntC1 (A) and EntC2 (B) and the calculation of their respective µH (hydrophobic moment) and H (hydrophobicity) using Heliquest software (http://heliquest.ipmc.cnrs.fr/). The arrow points towards the hydrophobic face of the helix. On the top of the wheel, the amino acid sequence that was used to generate the wheel is underlined in the respective sequence. On the right of the helical wheel projection, is the cartoon and stick representation of EntC1 or EntC2 structures as they were modelled within BacA/EntC complex by Alphafold. Peptides are represented with hydrophobic residues, according to the Eisenberg hydrophobicity scale, colored in red. Plot of the helical wheel projections of homologous LcnG and LcnQ peptides are indicated (right panels).

### Slide 11
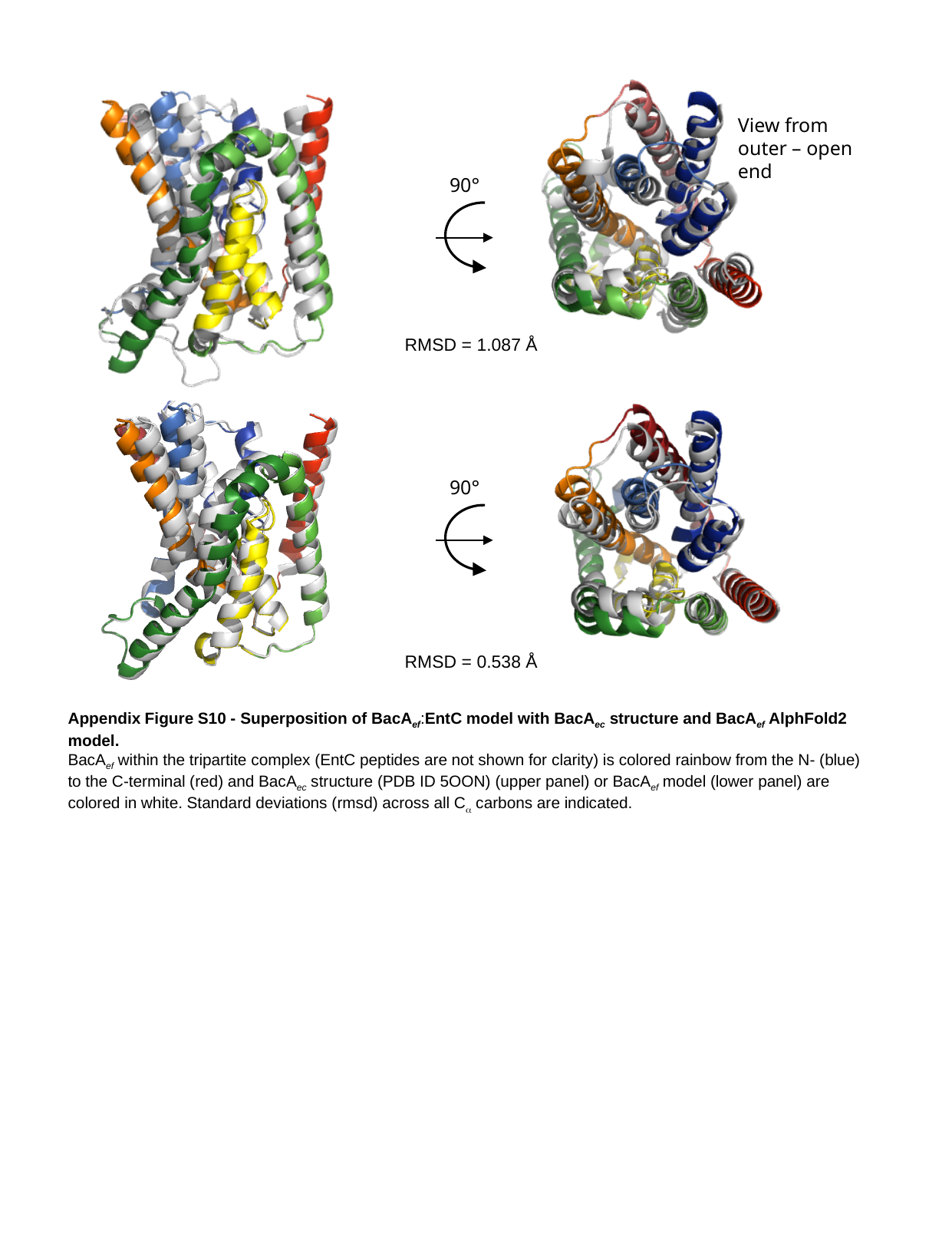

View from outer – open end
90°
RMSD = 1.087 Å
90°
RMSD = 0.538 Å
Appendix Figure S10 - Superposition of BacA_ef_:EntC model with BacA_ec_ structure and BacA_ef_ AlphFold2 model.
BacA_ef_ within the tripartite complex (EntC peptides are not shown for clarity) is colored rainbow from the N- (blue) to the C-terminal (red) and BacA_ec_ structure (PDB ID 5OON) (upper panel) or BacA_ef_ model (lower panel) are colored in white. Standard deviations (rmsd) across all C_α_ carbons are indicated.

### Slide 12
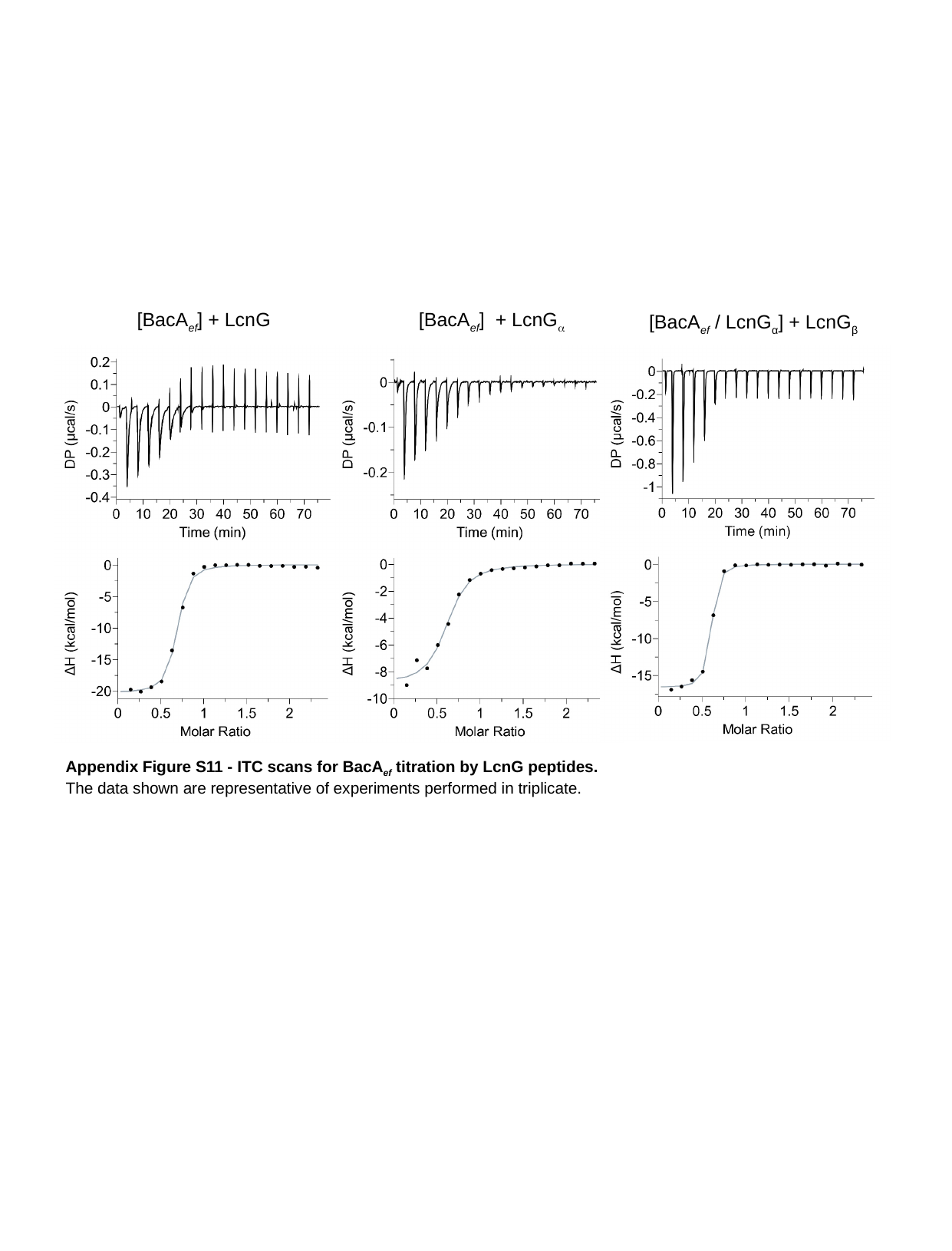

[BacA_ef_] + LcnG
[BacA_ef_] + LcnG_α_
[BacA_ef_ / LcnG_α_] + LcnG_β_
Appendix Figure S11 - ITC scans for BacA_ef_ titration by LcnG peptides.
The data shown are representative of experiments performed in triplicate.

### Slide 13
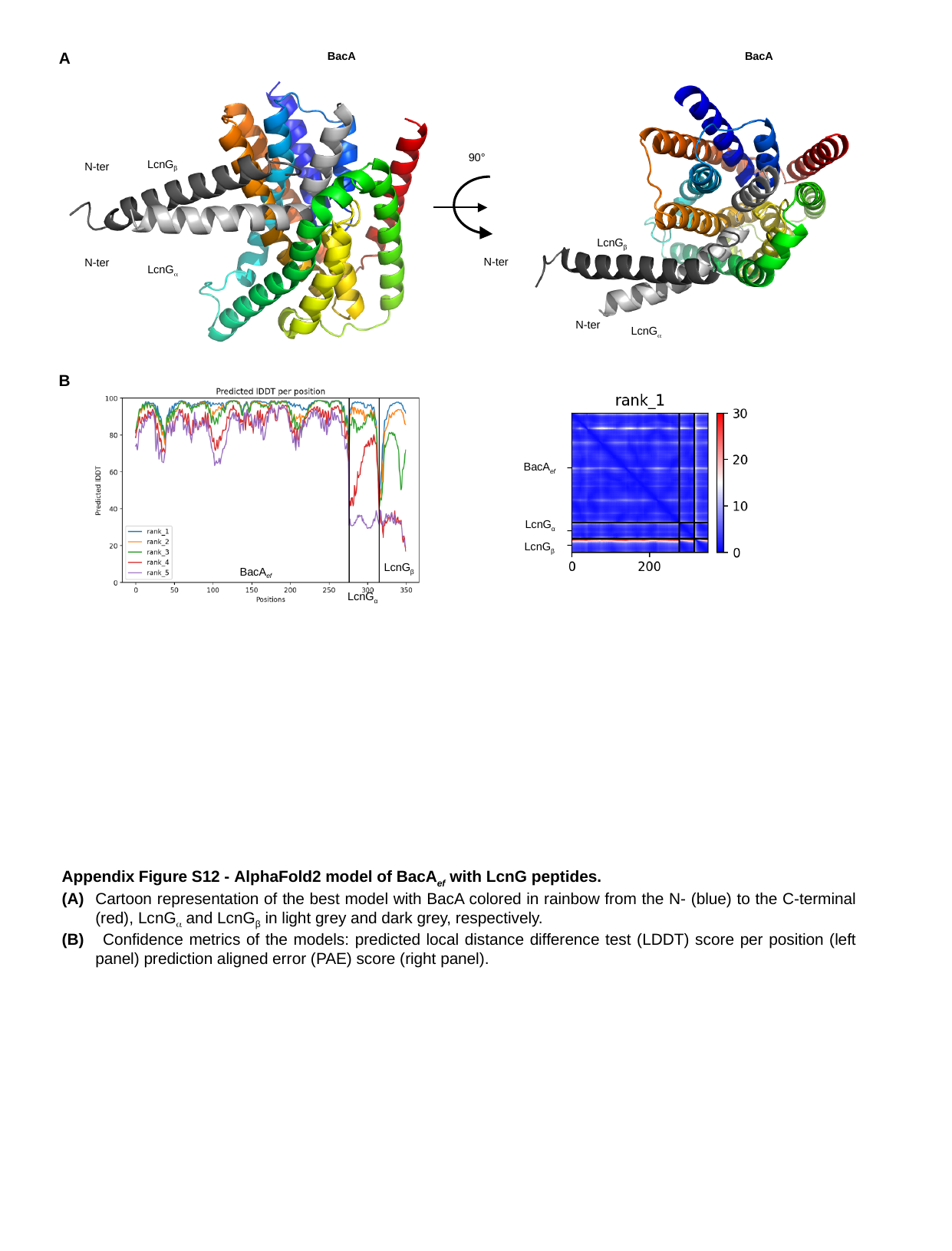

A
BacA
LcnG_β_
N-ter
N-ter
LcnG_α_
BacA
LcnG_β_
N-ter
N-ter
LcnG_α_
90°
B
LcnG_β_
BacA_ef_
LcnG_α_
BacA_ef_
LcnG_α_
LcnG_β_
Appendix Figure S12 - AlphaFold2 model of BacA_ef_ with LcnG peptides.
(A)	Cartoon representation of the best model with BacA colored in rainbow from the N- (blue) to the C-terminal (red), LcnG_α_ and LcnG_β_ in light grey and dark grey, respectively.
(B) 	 Confidence metrics of the models: predicted local distance difference test (LDDT) score per position (left panel) prediction aligned error (PAE) score (right panel).

### Slide 14
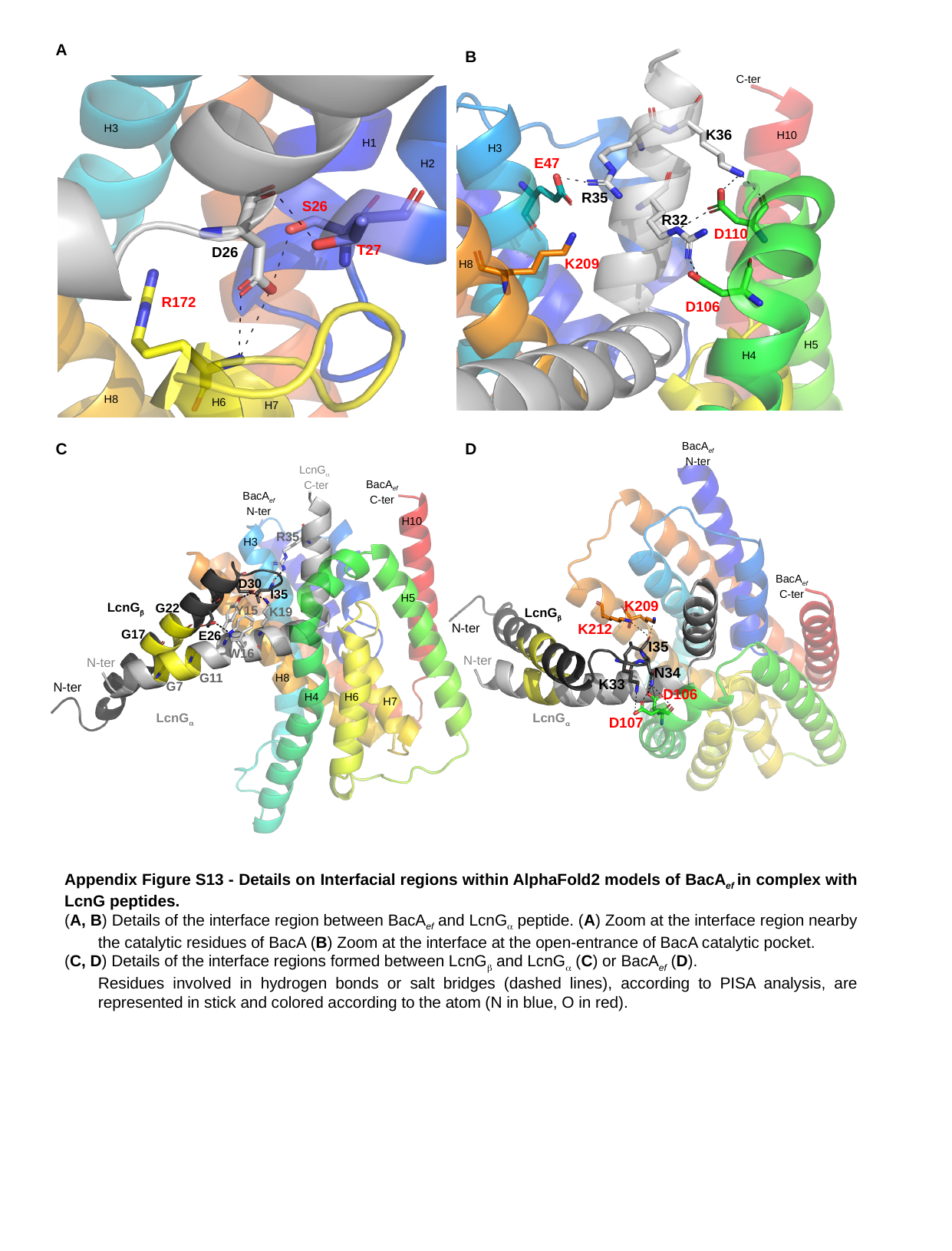

C-ter
K36
H10
H3
E47
R35
R32
D110
K209
H8
D106
H5
H4
A
B
H3
H1
H2
S26
T27
D26
R172
H8
H6
H7
K209
LcnG_β_
K212
I35
N34
K33
D106
LcnG_α_
D107
BacA_ef_
N-ter
LcnG_α_
C-ter
BacA_ef_
C-ter
BacA_ef_
N-ter
H10
R35
H3
BacA_ef_
C-ter
D30
I35
H5
LcnG_β_
G22
Y15
K19
N-ter
G17
E26
W16
N-ter
N-ter
G11
H8
G7
N-ter
H4
H6
H7
LcnG_α_
C
D
Appendix Figure S13 - Details on Interfacial regions within AlphaFold2 models of BacA_ef_ in complex with LcnG peptides.
(A, B) Details of the interface region between BacA_ef_ and LcnG_α_ peptide. (A) Zoom at the interface region nearby the catalytic residues of BacA (B) Zoom at the interface at the open-entrance of BacA catalytic pocket.
(C, D) Details of the interface regions formed between LcnG_β_ and LcnG_α_ (C) or BacA_ef_ (D).
	Residues involved in hydrogen bonds or salt bridges (dashed lines), according to PISA analysis, are represented in stick and colored according to the atom (N in blue, O in red).

### Slide 15
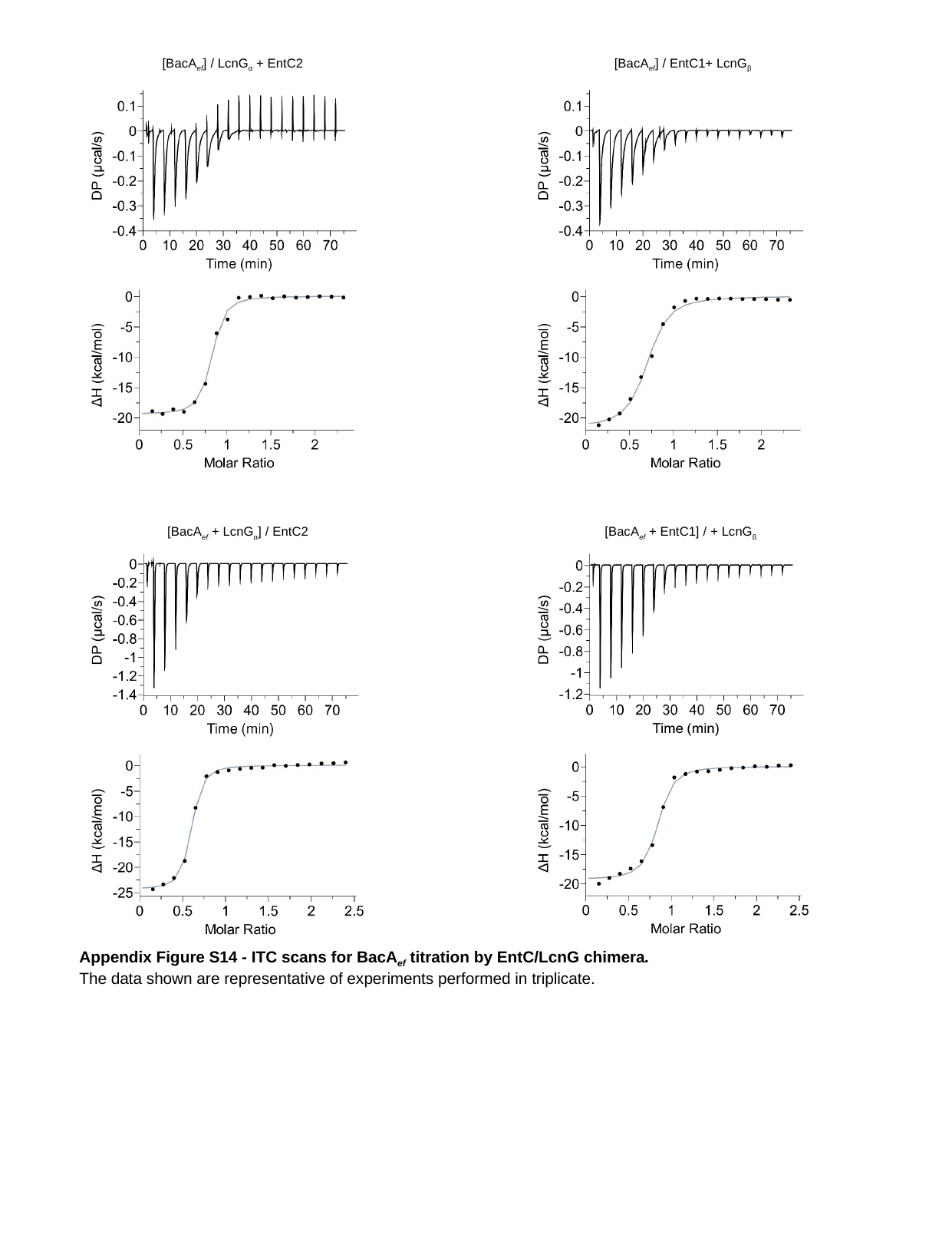

[BacA_ef_] / LcnG_α_ + EntC2
[BacA_ef_] / EntC1+ LcnG_β_
[BacA_ef_ + LcnG_α_] / EntC2
[BacA_ef_ + EntC1] / + LcnG_β_
Appendix Figure S14 - ITC scans for BacA_ef_ titration by EntC/LcnG chimera.
The data shown are representative of experiments performed in triplicate.

### Slide 16
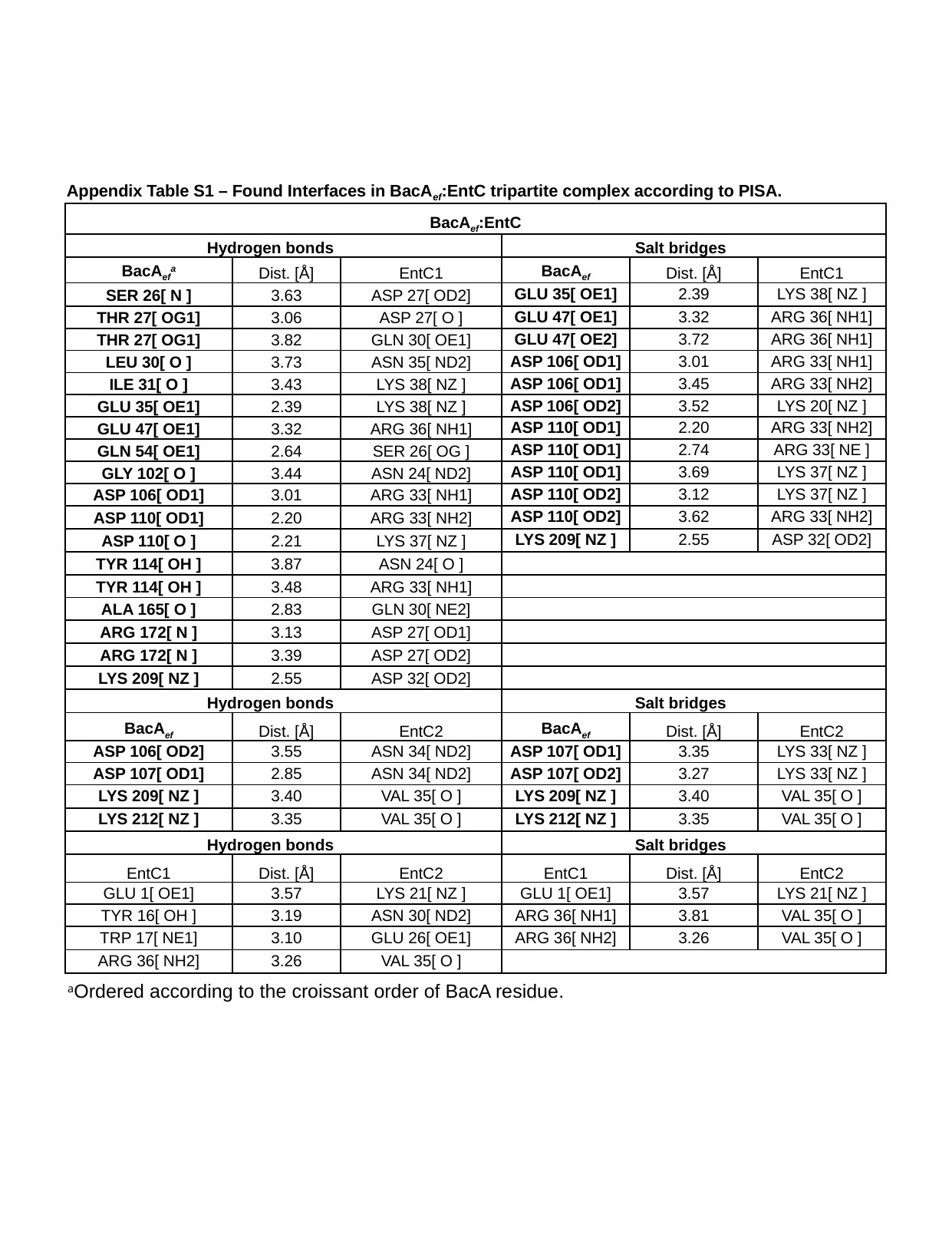

Appendix Table S1 – Found Interfaces in BacA_ef_:EntC tripartite complex according to PISA.
| BacA_ef_:EntC | | | | | |
| --- | --- | --- | --- | --- | --- |
| Hydrogen bonds | | | Salt bridges | | |
| BacA_ef_^a^ | Dist. [Å] | EntC1 | BacA_ef_ | Dist. [Å] | EntC1 |
| SER 26[ N ] | 3.63 | ASP 27[ OD2] | GLU 35[ OE1] | 2.39 | LYS 38[ NZ ] |
| THR 27[ OG1] | 3.06 | ASP 27[ O ] | GLU 47[ OE1] | 3.32 | ARG 36[ NH1] |
| THR 27[ OG1] | 3.82 | GLN 30[ OE1] | GLU 47[ OE2] | 3.72 | ARG 36[ NH1] |
| LEU 30[ O ] | 3.73 | ASN 35[ ND2] | ASP 106[ OD1] | 3.01 | ARG 33[ NH1] |
| ILE 31[ O ] | 3.43 | LYS 38[ NZ ] | ASP 106[ OD1] | 3.45 | ARG 33[ NH2] |
| GLU 35[ OE1] | 2.39 | LYS 38[ NZ ] | ASP 106[ OD2] | 3.52 | LYS 20[ NZ ] |
| GLU 47[ OE1] | 3.32 | ARG 36[ NH1] | ASP 110[ OD1] | 2.20 | ARG 33[ NH2] |
| GLN 54[ OE1] | 2.64 | SER 26[ OG ] | ASP 110[ OD1] | 2.74 | ARG 33[ NE ] |
| GLY 102[ O ] | 3.44 | ASN 24[ ND2] | ASP 110[ OD1] | 3.69 | LYS 37[ NZ ] |
| ASP 106[ OD1] | 3.01 | ARG 33[ NH1] | ASP 110[ OD2] | 3.12 | LYS 37[ NZ ] |
| ASP 110[ OD1] | 2.20 | ARG 33[ NH2] | ASP 110[ OD2] | 3.62 | ARG 33[ NH2] |
| ASP 110[ O ] | 2.21 | LYS 37[ NZ ] | LYS 209[ NZ ] | 2.55 | ASP 32[ OD2] |
| TYR 114[ OH ] | 3.87 | ASN 24[ O ] | | | |
| TYR 114[ OH ] | 3.48 | ARG 33[ NH1] | | | |
| ALA 165[ O ] | 2.83 | GLN 30[ NE2] | | | |
| ARG 172[ N ] | 3.13 | ASP 27[ OD1] | | | |
| ARG 172[ N ] | 3.39 | ASP 27[ OD2] | | | |
| LYS 209[ NZ ] | 2.55 | ASP 32[ OD2] | | | |
| Hydrogen bonds | | | Salt bridges | | |
| BacA_ef_ | Dist. [Å] | EntC2 | BacA_ef_ | Dist. [Å] | EntC2 |
| ASP 106[ OD2] | 3.55 | ASN 34[ ND2] | ASP 107[ OD1] | 3.35 | LYS 33[ NZ ] |
| ASP 107[ OD1] | 2.85 | ASN 34[ ND2] | ASP 107[ OD2] | 3.27 | LYS 33[ NZ ] |
| LYS 209[ NZ ] | 3.40 | VAL 35[ O ] | LYS 209[ NZ ] | 3.40 | VAL 35[ O ] |
| LYS 212[ NZ ] | 3.35 | VAL 35[ O ] | LYS 212[ NZ ] | 3.35 | VAL 35[ O ] |
| Hydrogen bonds | | | Salt bridges | | |
| EntC1 | Dist. [Å] | EntC2 | EntC1 | Dist. [Å] | EntC2 |
| GLU 1[ OE1] | 3.57 | LYS 21[ NZ ] | GLU 1[ OE1] | 3.57 | LYS 21[ NZ ] |
| TYR 16[ OH ] | 3.19 | ASN 30[ ND2] | ARG 36[ NH1] | 3.81 | VAL 35[ O ] |
| TRP 17[ NE1] | 3.10 | GLU 26[ OE1] | ARG 36[ NH2] | 3.26 | VAL 35[ O ] |
| ARG 36[ NH2] | 3.26 | VAL 35[ O ] | | | |
^a^Ordered according to the croissant order of BacA residue.

### Slide 17
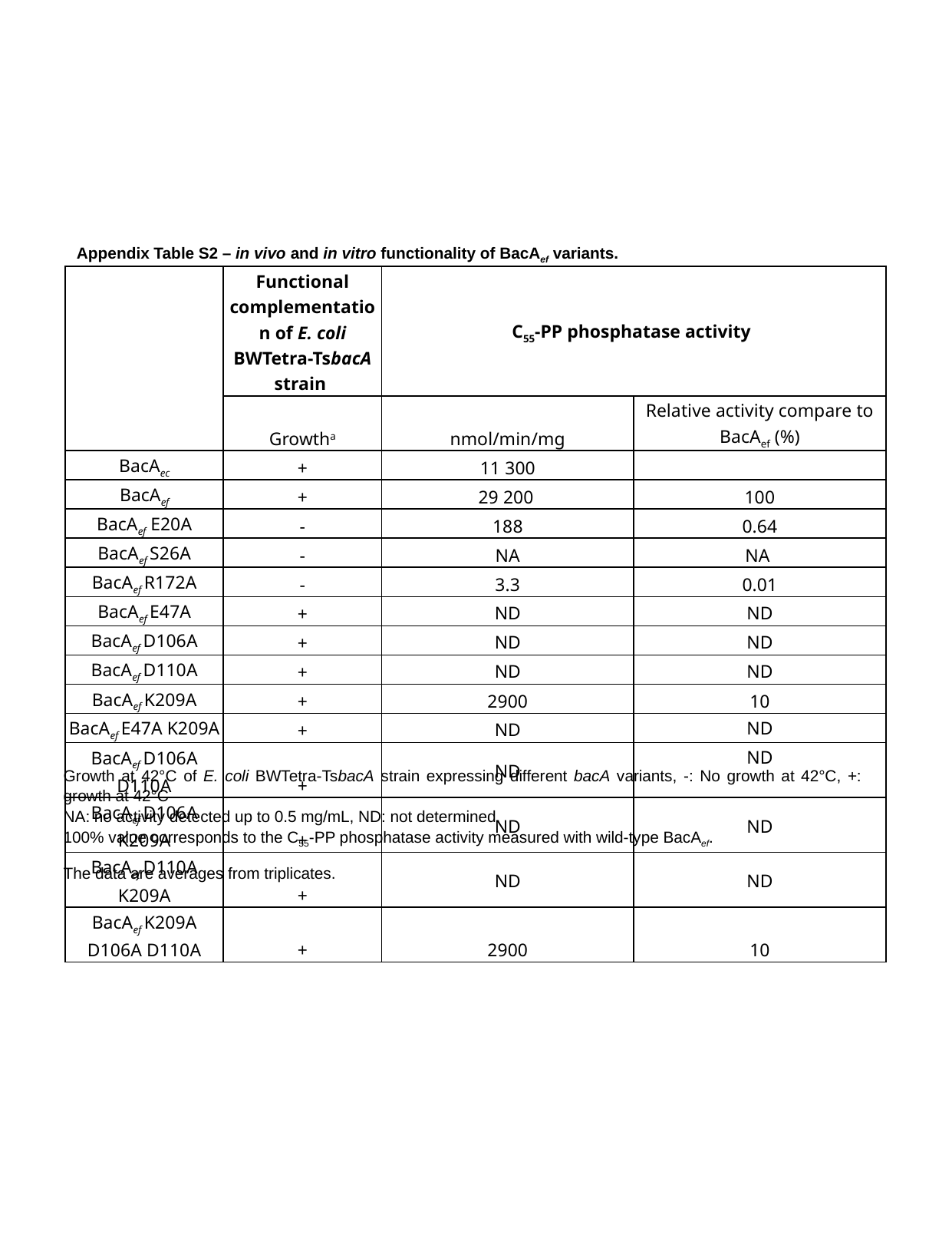

Appendix Table S2 – in vivo and in vitro functionality of BacA_ef_ variants.
| | Functional complementation of E. coli BWTetra-TsbacA strain | C_55_-PP phosphatase activity | |
| --- | --- | --- | --- |
| | Growth^a^ | nmol/min/mg | Relative activity compare to BacA_ef_ (%) |
| BacA_ec_ | + | 11 300 | |
| BacA_ef_ | + | 29 200 | 100 |
| BacA_ef_ E20A | - | 188 | 0.64 |
| BacA_ef_ S26A | - | NA | NA |
| BacA_ef_ R172A | - | 3.3 | 0.01 |
| BacA_ef_ E47A | + | ND | ND |
| BacA_ef_ D106A | + | ND | ND |
| BacA_ef_ D110A | + | ND | ND |
| BacA_ef_ K209A | + | 2900 | 10 |
| BacA_ef_ E47A K209A | + | ND | ND |
| BacA_ef_ D106A D110A | + | ND | ND |
| BacA_ef_ D106A K209A | + | ND | ND |
| BacA_ef_ D110A K209A | + | ND | ND |
| BacA_ef_ K209A D106A D110A | + | 2900 | 10 |
Growth at 42°C of E. coli BWTetra-TsbacA strain expressing different bacA variants, -: No growth at 42°C, +: growth at 42°C
NA: no activity detected up to 0.5 mg/mL, ND: not determined.
100% value corresponds to the C_55_-PP phosphatase activity measured with wild-type BacA_ef_.
The data are averages from triplicates.

### Slide 18
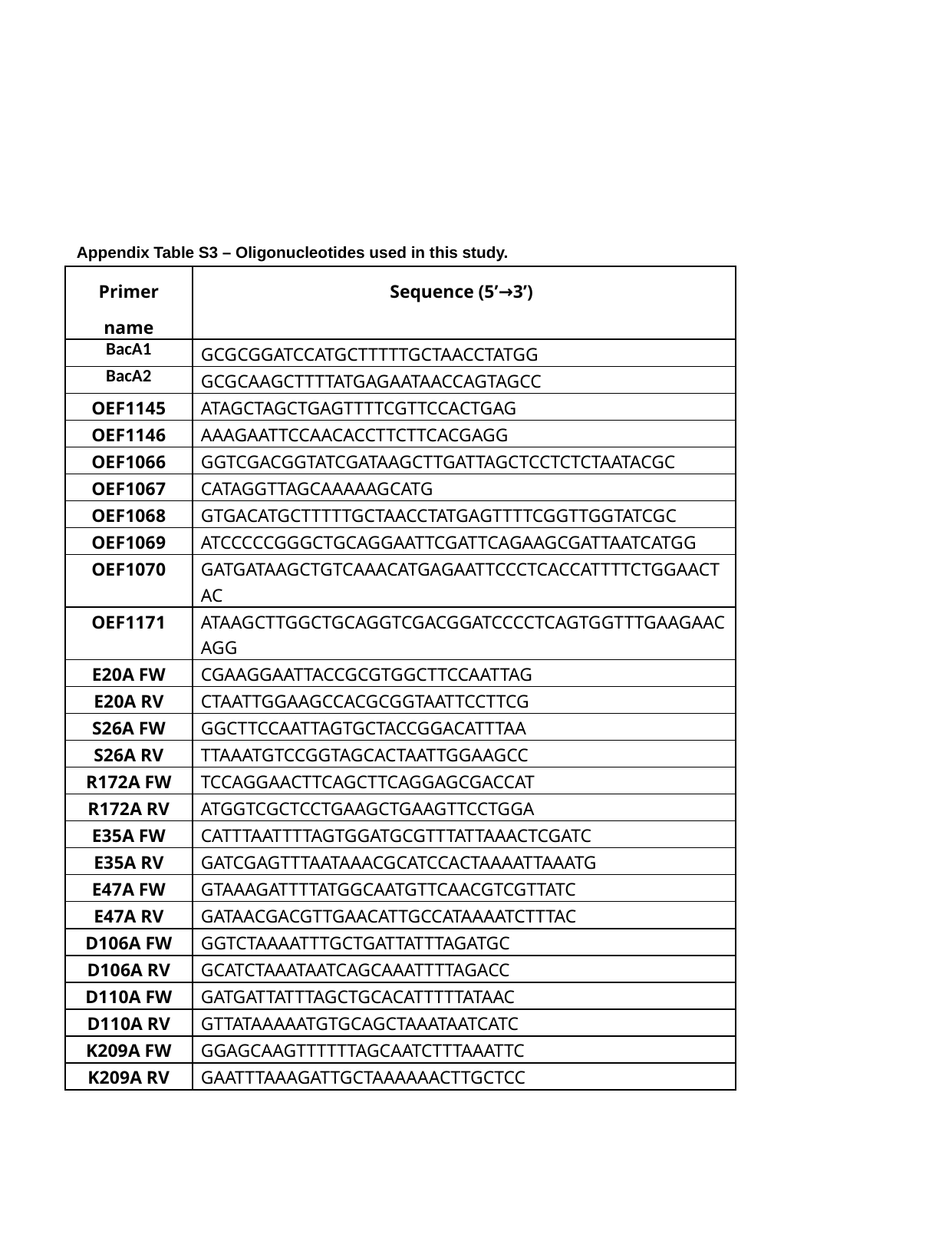

Appendix Table S3 – Oligonucleotides used in this study.
| Primer name | Sequence (5’→3’) |
| --- | --- |
| BacA1 | GCGCGGATCCATGCTTTTTGCTAACCTATGG |
| BacA2 | GCGCAAGCTTTTATGAGAATAACCAGTAGCC |
| OEF1145 | ATAGCTAGCTGAGTTTTCGTTCCACTGAG |
| OEF1146 | AAAGAATTCCAACACCTTCTTCACGAGG |
| OEF1066 | GGTCGACGGTATCGATAAGCTTGATTAGCTCCTCTCTAATACGC |
| OEF1067 | CATAGGTTAGCAAAAAGCATG |
| OEF1068 | GTGACATGCTTTTTGCTAACCTATGAGTTTTCGGTTGGTATCGC |
| OEF1069 | ATCCCCCGGGCTGCAGGAATTCGATTCAGAAGCGATTAATCATGG |
| OEF1070 | GATGATAAGCTGTCAAACATGAGAATTCCCTCACCATTTTCTGGAACTAC |
| OEF1171 | ATAAGCTTGGCTGCAGGTCGACGGATCCCCTCAGTGGTTTGAAGAACAGG |
| E20A FW | CGAAGGAATTACCGCGTGGCTTCCAATTAG |
| E20A RV | CTAATTGGAAGCCACGCGGTAATTCCTTCG |
| S26A FW | GGCTTCCAATTAGTGCTACCGGACATTTAA |
| S26A RV | TTAAATGTCCGGTAGCACTAATTGGAAGCC |
| R172A FW | TCCAGGAACTTCAGCTTCAGGAGCGACCAT |
| R172A RV | ATGGTCGCTCCTGAAGCTGAAGTTCCTGGA |
| E35A FW | CATTTAATTTTAGTGGATGCGTTTATTAAACTCGATC |
| E35A RV | GATCGAGTTTAATAAACGCATCCACTAAAATTAAATG |
| E47A FW | GTAAAGATTTTATGGCAATGTTCAACGTCGTTATC |
| E47A RV | GATAACGACGTTGAACATTGCCATAAAATCTTTAC |
| D106A FW | GGTCTAAAATTTGCTGATTATTTAGATGC |
| D106A RV | GCATCTAAATAATCAGCAAATTTTAGACC |
| D110A FW | GATGATTATTTAGCTGCACATTTTTATAAC |
| D110A RV | GTTATAAAAATGTGCAGCTAAATAATCATC |
| K209A FW | GGAGCAAGTTTTTTAGCAATCTTTAAATTC |
| K209A RV | GAATTTAAAGATTGCTAAAAAACTTGCTCC |
